# *Malat1* regulates female Th2 cell cytokine expression through controlling early differentiation and response to IL2

**DOI:** 10.1101/2024.10.23.619648

**Authors:** Mags Gwynne, Katie A. West, Stijn van Dongen, Ioannis Kourtzelis, Dawn Coverley, Sarah A. Teichmann, Kylie R. James, James P. Hewitson, Dimitris Lagos

## Abstract

Identifying cell intrinsic regulators of immune sexual dimorphism is critical for treatment of several immunopathologies. We show that *Malat1* is required for appropriate cytokine expression in female but not male Th2 cells. *Malat1* deficiency impairs *in vitro* Th2 differentiation of naïve CD4^+^ T cells from female mice, characterised by transcriptome-wide effects and suppression of cytokine expression, particularly IL10. Upon IL10R blockade a pronounced effect is also seen on IL4 and IL13. Mechanistically, naïve CD4^+^ T cells from *Malat1^-/-^* female mice demonstrate altered early activation kinetics and impaired early differentiation gene expression, including up-regulation of an interferon stimulated gene (ISG) module. This is followed by suppression of IL2Rα and IL2Rγ expression and IL2-mediated differentiation. Mimicking the effect of *Malat1* loss by maintaining early ISG expression in *WT* cells with IFNβ treatment partially phenocopies the effects of *Malat1* deficiency. A subset of the effects of *Malat1* loss in female cells is also observed in male cells. However, this does not affect endpoint Th2 differentiation. Male CD4^+^ T cells demonstrate stronger early activation, higher ISG expression during early differentiation, maintenance of IL2Rα expression independently of *Malat1*, and lower sensitivity to exogenous IL2 during late differentiation compared to female cells. *In vivo*, female, but not male, *Malat1^-/-^* mice demonstrate altered Th2 cytokine expression characterised by a reduction in IL10^+^ Th2 cells in both lung and spleen following priming and challenge with *Schistosoma mansoni* eggs, a model of lung type 2 inflammation. Overall, these findings reveal *Malat1* as a novel determinant of immune sexual dimorphism.

## Introduction

Biological sex is a known determinant of immune responses (1–4). Higher incidence or severity of autoimmune diseases (5–7) in females and infectious diseases in men (8) are commonly reported. The main drivers of immune sexual dimorphism are sex hormones (extrinsic) and X or Y chromosome-linked genes (intrinsic). For example, it is thought that females favour Th2 responses and males Th1 (2) and that this is due to hormonal regulation of immune genes such as IL4 (2, 9), and function of X chromosome-linked genes including the long intergenic non-coding RNA (lincRNA) *Xist* (10), which was recently shown to form ribonucleoprotein complexes that are directly involved in autoimmunity (11). Other mechanisms including those occurring at the post-transcriptional level, have been proposed, yet very little has been experimentally tested and validated. Overall, little is understood about cell intrinsic mediators of sexual dimorphism in Th cells that are not directly linked to sex hormone-mediated or sex chromosome-driven regulation.

Thousands of long non-coding transcripts, including lincRNAs, have been discovered in mammalian transcriptomes (12). In most cases, these are medium to lowly expressed transcripts displaying poor conservation across mammals. Our studies have focused on *Malat1*, a 7.5kb-long transcript discovered through an association with cancer progression and metastasis (13). *Malat1* is localised in nuclear speckles (14), which are nuclear foci enriched in factors involved in pre-mRNA splicing and transcription (15). It was initially thought that *Malat1* controls in mRNA splicing (14, 16). However, *Malat1^-/-^* mice develop normally and are viable and fertile without apparent defects in splicing (17–19). We previously discovered that *Malat1* is a key regulator of Th1 cell responses (20). We showed that *Malat1* is down-regulated upon initiation of Th cell differentiation, yet its complete loss results in stronger immune responses through suppression of IL10 expression in mouse models of type 1 immunity to parasitic infection, including visceral leishmaniasis and malaria. In humans, *Malat1* downregulation is a hallmark of proliferative CD4^+^ and CD8^+^ T cells (21). Others have linked *Malat1* to CD4^+^ T cell function, particularly in Th17 cells (22, 23), and to CD8^+^ T cell function (24).

In our previous report, which focused on Th1 cells, we showed that *Malat1* loss also affected IL10 expression in *in vitro* differentiated Th2 cells (20). Although expressed by all effector Th cells, IL10 expression is highest in Th2 cells (25), which we have also shown previously (20). However, the role of *Malat1* in Th2 cell differentiation and function remain under-explored. Here, we aimed to test how loss of *Malat1* affects Th2 differentiation *in vitro* and in an *in vivo* model of type 2 inflammation, *Schistosoma mansoni* (*S. mansoni*) egg injection (26, 27). We report that *Malat1* deficiency leads to impaired Th2 differentiation with widespread effects across the transcriptome and a notable suppression of IL10 expression only observed in female-derived Th2 cells. Although some effects caused by *Malat1* deficiency are shared between male- and female-derived CD4^+^ T cells, *Malat1* loss has more profound effects on female Th2 cells. This is due to a female-specific impaired regulation of an early differentiation interferon stimulated gene (ISG) expression programme, suppression of IL2 receptor (IL2R) expression in *Malat1^-/-^* cells, and increased sensitivity of female cells to IL2-driven cytokine production.

## Materials and Methods

### Animals and ethics

Animal care and experimental procedures were regulated under the Animals (Scientific Procedures) Act 1986 (revised under European Directive 2010/63/EU) and were performed under U.K. Home Office License (project license number PP0841992 for breeding and PP9423191for *S. mansoni* egg injections) with approval from the University of York Animal Welfare and Ethical Review Body. Animal experiments conformed to Animal Research: Reporting of In Vivo Experiments guidelines (28).

### *S. mansoni* egg injection

C57BL/6 CD45.2 *wildtype* (*WT*) mice were obtained from Charles River Laboratories. *Malat1^−/−^* mice (complete knockouts) were obtained from the Riken Institute (18). All mice were bred in-house, maintained under specific pathogen-free conditions, and used at 6–12 weeks of age. Schistosome eggs were recovered from the livers of C57BL/6 mice at week 7 post-infection following exposure to 100 *S. mansoni* cercariae. Cercariae were obtained from schistosome-infected *Biomphalaria glabrata* snails provided by the Barrett Centre for Helminth Control (Aberystwyth University, UK). Livers were digested overnight at 37°C with shaking with 0.2U/ml collagenase D (Roche) in the presence of 5000U/ml polymyxin B (Merck). Eggs were purified by centrifugation through 10ml percoll (GE Healthcare) / 20ml 0.25M sucrose (450 x*g*, 5min, RT), washed in PBS and stored at -20°C before usage. 5000 dead *S. mansoni* eggs in 200 μl PBS were delivered via intraperitoneal (ip) injection into mice. Two weeks later, the mice were intravenously (iv) challenged with another 5000 eggs in 200 μl PBS. After another week the mice were sacrificed and the lungs and spleen were extracted and processed. Lungs were digested with 0.4 U/ml Liberase TL (Roche) and 80 U/ml DNase I type IV in HBSS (both Sigma-Aldrich) for 45 min at 37°C. Enzyme activity was inhibited with 10 mM EDTA (pH 7.5) and single-cell suspensions were created with 100-μm nylon filters (Falcon) in complete RPMI 1640 (Thermo Fisher Scientific) supplemented with 10% heat-inactivated FCS (HyClone), 100 U/ml penicillin, 100 μg/ml streptomycin, and 2 mM l-glutamine (all Thermo Fisher Scientific), then cleaned via Percoll gradient and 3ml Ammonium-Chloride-Potassium (ACK) red blood cell lysis buffer. Spleen single cell suspensions were generated in the same manner and only passed through a 70 μm filter and treated with 3 ml ACK buffer.

### *In vitro* Th2 differentiation and *Malat1* knockdown

Spleens and axillary, brachial, mesenteric, and inguinal lymph nodes of *WT* or *Malat1^-/-^*mice were extracted, homogenised, through a 70 μm filter in RPMI 1640. Resulting cell pellets were then treated with 3ml ACK lysis buffer. Naïve CD4^+^ T cells were then purified via MACS column isolation (Miltenyi Naive CD4^+^ T Cell Isolation Kit, mouse, catalogue number 130-104-453) normally resulting in purity of ∼95%. For Th2 polarisation, purified naïve CD4^+^ T cells (500,000 cells per well) were stimulated with 10 μg/ml plate-bound anti-CD3ε (clone 145-2C11) and 4 μg/ml soluble anti-CD28 (37.51) in RPMI 1640 in flat-bottom 96-well plates in the presence of 25 ng/ml mouse rIL4 and 5 μg/ml anti–IFN-γ (XMG1.2). Anti-CD3/anti-CD28–dependent activation (4 days) was followed by rest in 10 U/ml human recombinant IL2 for 2 days. Titrations of anti-CD3ε antibody or recombinant IL2 were performed as indicated. Recombinant cytokines were from PeproTech. For *Malat1* knockdown experiments, control or *Malat1*-targeting antisense oligonucleotide GapmeRs were from QIAGEN (Hilden, Germany; LG00000002-DDA and LG00000008-DDA, respectively) and were added to naive CD4^+^ T cells on day 0, or differentiating Th2 cells on day 4, at a final concentration of 100 nM. For IL10 receptor blockade experiments, CD4^+^ T cells were treated either with anti-IL10R (Clone: 1B1.3A from Bio X Cell) or rat IgG at 10 μg/ml as a control, at both day 0 and day 4. For IFNβ treatment experiments, cells were treated with 5000 U/ml mouse IFNβ (CYT-651, from Prospec), at day 0.

### Flow Cytometry

For flow cytometry analysis, single cell suspensions were generated. For live/dead discrimination, cells were washed twice in PBS, then stained with Zombie Aqua (BioLegend) in PBS before resuspension in FACS (Fluorescence-Activated Cell Sorting) buffer (PBS containing 0.5% BSA and 0.05% azide). Fc receptors were blocked with 100 μg/ml rat IgG (Sigma-Aldrich) for 10 min at 4°C before surface staining for 30 min at 4°C. The following anti-mouse Abs from Biolegend were used: CD45.2 BV786 (clone 104); TCRβ PE-Cy7 (H57-597); CD19 APC-Cy7 (6D5); MHC class II (MHCII) Alexa Fluor 700 (M5/114.15.2); Ly-6G APC-Cy7 (1A8); Ly6C BV605 (HK1.4); CD64 PE (X54-5/7.1); CD11b Pacific Blue (M1/70); CD44 FITC (IM7); CD62L PE (MEL-14); CD8α PB (53-6.7); CD4 PerCP/Cy5.5 (RM4-5); IFN-γ FITC (XMG1.2); IL10 PE (JES5-16E3); IL4 PE-Dazzle or APC (11B11); CD25 (IL-2Ra) PerCP- Cy5.5 or APC (PC6.1); CD69 APC (H1.2F3); CD132 (IL-2Rg) PE (TUGm2); GATA3 PE-Dazzle 594 (16E10A23); IFNAR1-PE (MAR1-5A3) and streptavidin PE-Cy7. These were used in combination with abs SiglecF PerCP-eFluor 710 (1RNM44N), iNOS PE-eFluor 610 (CXNET) and goat anti-rabbit A647 from ThermoFisher Scientific, biotinylated Ym1 from R&D systems, anti-murine RELMα from PeproTech, and ; IL-13 PB (eBio13A) from eBioscience. To measure intracellular cytokines in T cells following *ex vivo* stimulation, cells were first stimulated in complete RPMI 1640 for 4 h at 37°C with 500 ng/ml PMA, 1 μg/ml ionomycin, and 10 μg/ml brefeldin A (all Sigma-Aldrich). For all intracellular cytokine staining, surface stained cells were fixed and permeabilized (20 min at 4°C) using Fixation/Permeabilization Solution before washes in Perm/Wash buffer (both BD Biosciences). Cells were then stained with intracellular Abs as above except in Perm/Wash buffer. For transcription factor staining, surface stained cells were fixed and permeabilized (1h or overnight at 4°C) using the FoxP3 Fixation/Permeabilization solution before washes in 1x Permeabilization Buffer (eBioscience). Cells were stained with intracellular Abs as above except in Permeabilization Buffer. Appropriate isotype or fluorescence minus one (FMO) controls were used to draw gates for populations of interest. Events were acquired on an LSRFortessa (BD Biosciences) with analysis by FlowJo (FlowJo) or a Cytoflex LX (Beckman Coulter) before analysis with or CytExpert.

### RNA extraction, cDNA synthesis, and quantitative RT-PCR

RNA was extracted from purified cell populations using QIAzol and miRNeasy RNA extraction kits (QIAGEN) according to manufacturer’s instructions. For mRNA transcripts, reverse transcriptions were carried out with Superscript III (Thermo Fisher Scientific) and random hexamer primers (Promega) and measured with Fast SYBR Green Master Mix (Thermo Fisher Scientific). Quantitative PCR was performed using a StepOnePlus Real Time PCR System or a QuantStudio 3 (both Thermo Fisher Scientific), and relative transcript levels were determined using the ΔΔCt method. The primers used for the study are show in **Supplemental Table S1**.

### RNA sequencing (RNAseq) and analysis

We used short-read (Illumina) RNAseq to compare naïve CD4^+^ T cells and *in vitro* differentiated Th2 cells from female and male *WT* and *Malat1^-/-^* mice. Pooled cDNA libraries were sequenced on one lane of an Illumina Novaseq 6000 S4 flow-cell at 100PE. The raw sequencing reads were aligned to the mouse genome version Ensembl GRCm38.92 with STAR 2.5.4a. We sequenced four replicates per condition. Samples with assigned sequencing reads lower than 500,000 were removed from downstream analysis. Of the remaining samples (N=4 for all *Malat1^-/-^* samples and *WT* male Th2, N=3 for *WT* female and male naïve CD4^+^ T cells and *WT* female Th2), we obtained an average of 65 million reads per sample (range: 52– 91 million). Analysis was performed in RStudio (version 1.4.1106) with R (v 4.1). Data was normalised (‘calcNormFactors’ and ‘estimateDisp’ functions of edgeR (version 3.14.0)) and a quasi-likelihood negative binomial generalized log-linear model (‘glmQLFit’ and ‘glmQLFTest’ functions with coefficient = 2) was used to determine differentially expressed genes. Adjusted p values were calculated using the Benjamini and Hochberg method.

We used Oxford Nanopore Technologies (ONT) long-read RNAseq to compare naïve CD4^+^ T cells and cells after 2 days of *in vitro* differentiation from female *WT* and *Malat1^-/-^* mice (N=4 per group). Full length cDNA libraries were prepared using the ONT cDNA-PCR Sequencing V14 - Barcoding kit (SQK-PCB114.24). Briefly, cDNA RT adapters were ligated to 3’ ends (polyA tails) of transcripts, prior to stand switching cDNA synthesis and a 13 cycle PCR reaction with barcoded primers, labelling each cDNA sample with a unique DNA barcode. Barcoded cDNAs were pooled at equimolar ratios before final adapter ligation and sequencing on R10.4.1 flowcells in ONT PromethION sequencer (8 samples per flow cell). Live superaccuracy basecalling and barcode demultiplexing were performed in MinKNOW software (version 24.06.10) at the time of the run. We acquired 0.745-1.121 million reads per sample, with 604-840 bases mean read length. Demultiplexed reads were analyzed through the workflow Transcriptomes from Epi2Me^TM^ application, specifically designed by ONT. The workflow generates differential gene expression data (with edgeR), as well as differential transcript usage (DTU) data (with Dexseq), using a reference transcriptome. The Viking cluster, provided by the University of York, was used for the analyses.

### Statistics

Experiments were typically performed in 2 – 3 independent replicates in groups of N = 3 – 5 mice per condition and replicate. Statistical analyses were carried out as indicated with Prism 5 (GraphPad Software). Two-way comparisons used paired or unpaired t tests as indicated, and multiple comparisons used one-way ANOVA, followed by Sidak’s multiple comparisons test for comparison of samples with biological relevance (e.g. *WT* female to *Malat1^-/-^*female, *WT* male to *Malat1^-/-^* male, *WT* female to *WT* male). Confidence levels were set to 0.05 for significance. In merged data where n>9, consensus between Anderson-Darling, Shapiro-Wilk and D’Agostino & Pearson tests were used to determine normality. If all samples were normally distributed one-way ANOVA was performed as described. If one or more samples did not follow a normal distribution, a Kruskal-Wallis test followed by Dunn’s multiple comparison test were performed to identify significance, with confidence levels of 0.05. P values are displayed as asterisks representing P value classification, where * ≤ 0.05, ** ≤ 0.01, ***≤ 0.001 and **** ≤ 0.0001, unless the results are borderline non-significant, in which case the p value is shown.

### Data availability

RNAseq data have been deposited in the National Center for Biotechnology (NCBI) Gene Expression Omnibus (GEO) database. Accession numbers: GSE279185 for Illumina RNAseq and GSE278413 for Oxford Nanopore Technologies RNAseq data.

## Results

### *Malat1* loss impairs *in vitro* Th2 differentiation of CD4^+^ T cells from female mice

We first assessed the effect *of Malat1* deletion on Th2 differentiation *in vitro* in response to stimulation with anti-CD3 and anti-CD28 antibodies in the presence of recombinant IL4 and anti-IFNγ antibody for 4 days, followed by IL2 treatment for 2 days. This protocol allowed us to dissect potential effects on naïve CD4^+^ T cell activation (0-24h), early differentiation (2-4 days), and terminal differentiation in response to exogenous IL2 (**Fig. 1A**). Following *in vitro* differentiation of naive CD4+ T cells from female mice to Th2 cells, *Malat1^-/-^* cells displayed significantly lower percentages of IL10^+^ and IL4^+^/IL10^+^ cells at day 6 of differentiation, with no differences in IL4^+^/IL10^-^ cells, and no differences at day 4 (**Fig. 1B – F** and **Supplemental Fig. 1A**). This decrease in cytokine expression was not observed from cells derived from male mice. Under these conditions, differentiation of *WT* CD4^+^ T cells from male mice was less efficient compared to female *WT* cells, and *Malat1* deletion did not affect this further (**Fig. 1B – F**). RNAseq analysis of *WT* and *Malat1^-/-^* naïve CD4^+^ T cells and *in vitro* differentiated Th2 cells revealed only a small number of genes were statistically significantly differentially expressed between *WT* and *Malat1^-/-^* cells (**Supplemental Table S2**). As shown before (20), IL10 levels were suppressed in *Malat1^-/-^* Th2 cells compared to *WT* controls (Log2 fold change LFC = - 0.882, p = 0.034, for female-derived cells and LFC = 0.03, p = 0.932 for male-derived cells), although this did not reach significance after multiple testing correction. This was further confirmed by qRTPCR for the sequenced samples (**Supplemental Fig. S1B**), suggesting that this bulk RNAseq analysis provided a conservative representation of the effect of *Malat1* deficiency on Th2 gene expression. Focusing on genes that were differentially expressed between Th2 and naïve CD4^+^ T cells from *WT* mice (Th2-differentiation associated genes) we observed a transcriptome-wide blunting of Th2 gene expression upon *Malat1* loss in female but not male cells (**Fig. 1G – H** and **Supplemental Tables S3 – S5**). Analysis of all (FDR<0.001) or the top (by FDR) 500 differentially expressed genes (DEGs) between *WT* Th2 and naïve CD4^+^ T cells demonstrated that most genes that were up-regulated during *WT* Th2 differentiation were overall expressed at lower levels when comparing *Malat1^-/-^* Th2 cells to *WT* Th2 cells, and, conversely, genes that were suppressed upon *WT* Th2 differentiation were overall more highly expressed in *Malat1^-/-^* Th2 cells (**Fig. 1G** and **Supplemental Tables S3 – S5**). This transcriptome-wide trend was not observed in male cells (**Fig. 1H**). Taking a stricter threshold for DEGs between Th2 and naïve CD4^+^ T cells from female mice (FDR<0.001 and absolute LFC>2), we found 524 Th2-differentiation associated DEGs in both *WT* and *Malat1^- /-^*, 469 DEGs only in *WT*, and 139 DEGs only in *Malat1^-/-^* Th2 cells. We noted that common and *WT* only Th2 DEGs were mainly up-regulated, whereas *Malat1^-/-^* only Th2 DEGs were mainly down-regulated (**Supplemental Fig. S1C – D**, **Supplemental Tables S3 – S5**).

**Figure 1.**
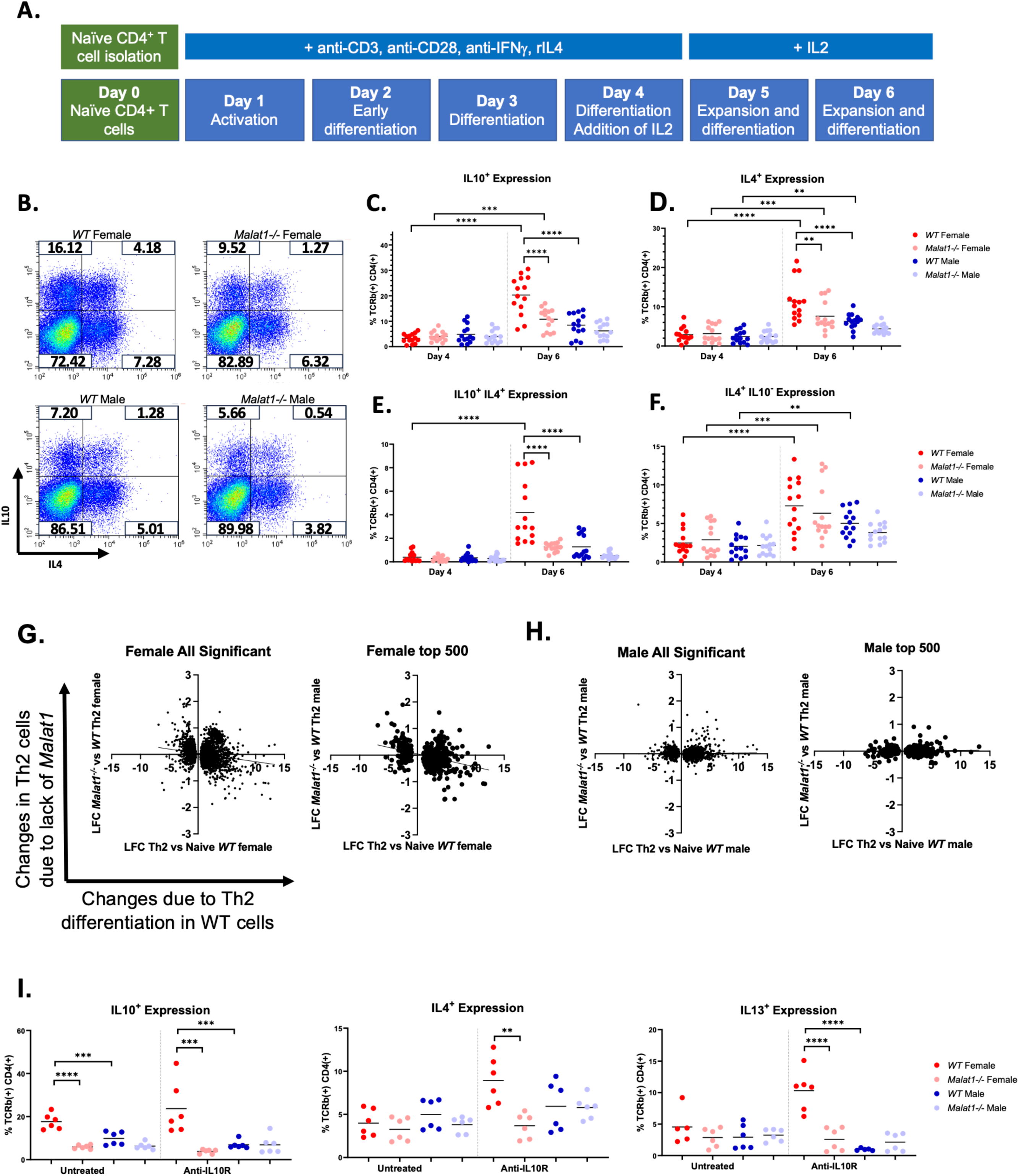
*Malat1* loss impairs *in vitro* Th2 differentiation only in female cells. A. Schematic of *in vitro* Th2 differentiation protocol. B. Representative FACS plots of IL10 and IL4 expression in *WT* and *Malat1^-/-^* female and male *in vitro* differentiated Th2 cells (Day 6). C. Percentage of IL10^+^ live TCRb^+^ CD4^+^ in *in vitro* differentiated Th2 cells derived from female and male mice at day 4 (prior to IL2 addition) and day 6 (experimental end-point). Levels determined by intracellular cytokine staining (N=14 per condition). D. As in **B**, but for IL4^+^ live TCRb^+^ CD4^+^ cells. E. As in **B**, but for IL4^+^/IL10^+^ live TCRb^+^ CD4^+^ cells. F. As in **B**, but for IL4^+^/IL10^-^ live TCRb^+^ CD4^+^ cells. G. Log2 fold change (LFC) in gene expression between *WT* female naïve and *in vitro* differentiated Th2 cells against LFC in gene expression between *WT* and *Malat1^-/-^* female *in vitro* differentiated Th2 cells. Data shown for all or the top 500 DEGs between Th2 and naïve CD4^+^ T cells from *WT* female mice. H. As in **E**, but for cells derived from male mice. I. Percentage of IL10^+^ live TCRb^+^ CD4^+^ *in vitro* differentiated Th2 cells derived from *WT* or *Malat1^-/-^* female and male mice at day 6, with or without treatment with 10 μg/ml of anti-IL10 receptor antibody. Levels determined by intracellular cytokine staining (N=6 per condition).

Furthermore, when looking at common DEGs, we found that the vast majority of up-regulated genes showed a smaller LFC in *Malat1^-/-^* Th2 cells compared to *WT* (**Supplemental Fig. S1E**). To distinguish the effect of *Malat1* from that of IL10, we blocked the IL10 receptor (IL10R) between days 4 and 6 of differentiation. As above, in control conditions, the effect of *Malat1* on IL4 and IL13 effector Th2 cytokines was modest. However, upon blockade of IL10R, *Malat1* loss resulted in dramatic and statistically significant decrease in IL4 and IL13 expression in female-derived Th2 cells, with no effect on male-derived cells (**Fig. 1I**). Overall, these results indicated that *Malat1* loss impairs Th2 differentiation of female- but not male-derived naïve CD4^+^ T cells, characterised by suppression of IL10 expression and an impairment of the Th2 gene expression programme.

### The effect of *Malat1* depletion on IL10 expression is independent of activation strength and occurs during the early stages of differentiation

We next tested whether *Malat1* deficiency affected Th2 polarization under weaker activation conditions. We found that *Malat1^-/-^*cells displayed lower IL10 expression under suboptimal differentiation conditions, whereas IL4 levels were not statistically significantly different (**Fig. 2A – B**, **Supplemental Fig. S2A – B**). Of note, at lower activation levels there were no differences in IL10 expression between *in vitro* Th2 differentiation of male and female cells. This indicated that *Malat1* deficiency can affect Th2 cytokine expression independently of activation strength. Under suboptimal differentiation conditions, IL10R blockade did not enhance the effect of *Malat1* deficiency on IL4 or IL13 expression, potentially due to the low cytokine expression levels under these conditions (**Supplemental Fig. S2C**).

**Figure 2.**
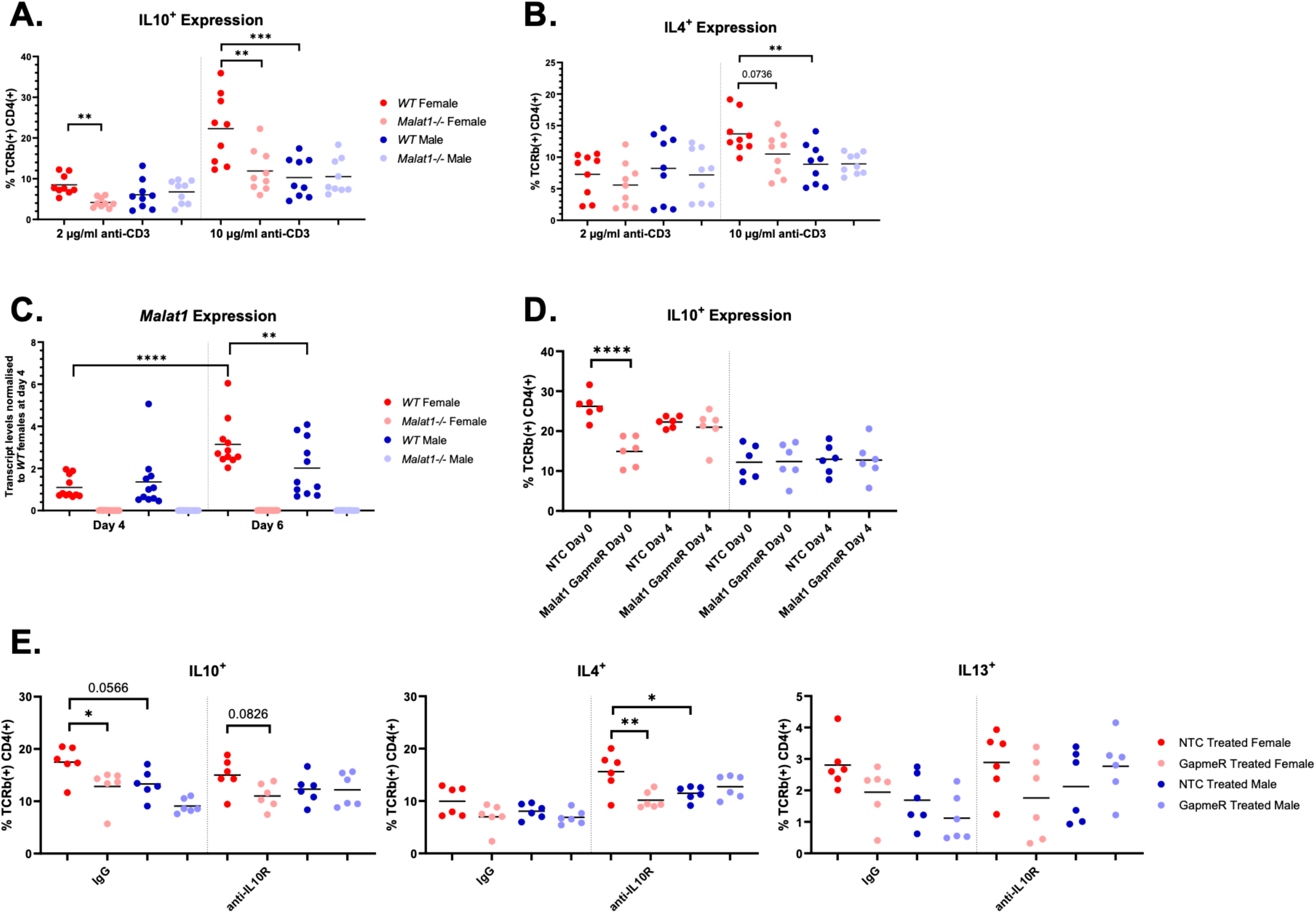
The effect of *Malat1* depletion on IL10 expression is independent of activation strength and occurs during the early stages of differentiation. **A.** Percentage of IL10^+^ live TCRb^+^ CD4^+^ *WT* or *Malat1^-/-^ in vitro* differentiated Th2 cells derived from female and male mice, with 10 μg/ml or 2 μg/ml of anti-CD3 antibody (see Methods). Levels determined by intracellular cytokine staining (N=9). **B.** As in **A**, but for IL4^+^ live TCRb^+^ CD4^+^ cells. **C.** *Malat1* levels in *in vitro* differentiated Th2 cells at day 4 (prior to resuspension in IL2) and at day 6 (experimental end point), determined by qRTPCR. Levels normalised to U6 and average levels of *WT* females at day 4 (N=11). **D.** Percentage of IL10^+^ live TCRb^+^ CD4^+^ *in vitro* differentiated *WT* Th2 cells derived from female and male mice, treated with either non-targeting control (NTC) GapmeRs or *Malat1* targeting GapmeRs. Levels determined by intracellular cytokine staining (N=6). **E.** Percentage of IL10^+^, IL4^+^, and IL13^+^ live TCRb^+^ CD4^+^ *in vitro* differentiated *WT* Th2 cells derived from female and male mice treated with NTC or *Malat1* targeting GapmeRs, with and without treatment with an anti-IL10 receptor antibody. Levels determined by intracellular cytokine staining (n=6).

We next tested whether *Malat1* knockdown rather than genetic deletion had a sex-specific effect on Th2 differentiation. As described previously (20), we used GapmeR oligonucleotides to suppress *Malat1* expression (**Supplemental Fig. S2D**). Although, Th2 cells express lower *Malat1* levels than naïve CD4^+^ T cells ((20) and **Supplemental Fig. S2E**), we found that *Malat1* levels increase between day 4 and 6 of differentiation, upon cessation of CD3 activation and addition of exogenous IL2 (**Fig. 2C**). As for cytokine expression (**Fig. 2A-B**), the increase in *Malat1* levels in response to IL2 (day 4 to day 6) was more profound in female cells. We note that there were no differences in *Malat1* expression between *WT* male and female naïve **(Supplemental Fig. S2E)** or differentiating (day 4; **Fig. 2C**) CD4^+^ T cells prior to IL2 stimulation.

Based the observation that *Malat1* deficiency impaired cytokine induction between days 4 and 6 (**Fig. 1C – F**), we knocked down *Malat1* either on day 0, or by adding GapmeRs on day 4, at the same time with exogenous IL2 stimulation. *Malat1* knockdown reduced IL10 expression in female Th2 cells only if GapmeRs were added to naïve CD4^+^ T cells, but not when added concurrently with exogenous IL2 stimulation. No effects on cytokine expression were observed upon *Malat1* knockdown in male cells (**Fig. 2D**). Partial knockdown of *Malat1* in cells from female mice did not affect IL4 expression (**Supplemental Fig. S2F)**. As in the case of *Malat1^-^*

*^/-^* cells, IL10R blockade enhanced the effect of *Malat1* GapmeR-mediated knockdown on IL4 (**Fig. 2E**). This was not observed for IL13, potentially due to low IL13 expression under these conditions (**Fig. 2E**). We noticed that treatment with non-targeting control GapmeRs consistently resulted in lower IL13 cytokine expression in Th2 cells (compare **Fig. 2E** with **Fig. 1I**). Overall, this demonstrated that the effect of *Malat1* on Th2 cytokine expression required its activity during the early differentiation stages.

### *Malat1* loss alters the kinetics of naïve CD4^+^ T cell activation and impairs early Th2 differentiation

Based on the above, we tested the effect of *Malat1* deletion on the early stages of Th2 differentiation. First, we studied the first 24h of naïve CD4^+^ T cell activation by measuring levels of T cell early activation marker CD69 (**Supplemental Fig. S3A**). As expected, *Malat1* levels were suppressed within 24h in both male and female *WT* mice (**Supplemental Fig. S3B**). In female cells, we found that compared to *WT* cells CD69 protein expression was higher in *Malat1^-/-^*cells at 3h post activation, the difference reaching statistical significance at 7h post activation. However, CD69 expression was significantly lower in *Malat1^-/-^* cells at 24h (**Fig. 3A**). At the transcript level, *Cd69* levels peaked at 3h post activation, with significantly higher levels in *Malat1^-/-^* cells (**Supplemental Fig. S3C**). In male cells, we did not observe any statistically significant differences between *WT* and *Malat1^-/-^* cells, although we noted that male *WT* cells showed statistically significant higher CD69 levels compared to female counterparts at 3h post activation (**Fig. 3A**). We also measured levels of IL2Rα (CD25), which was induced upon activation but found no differences between genotypes or biological sexes (**Supplemental Fig. S3D**).

**Figure 3.**
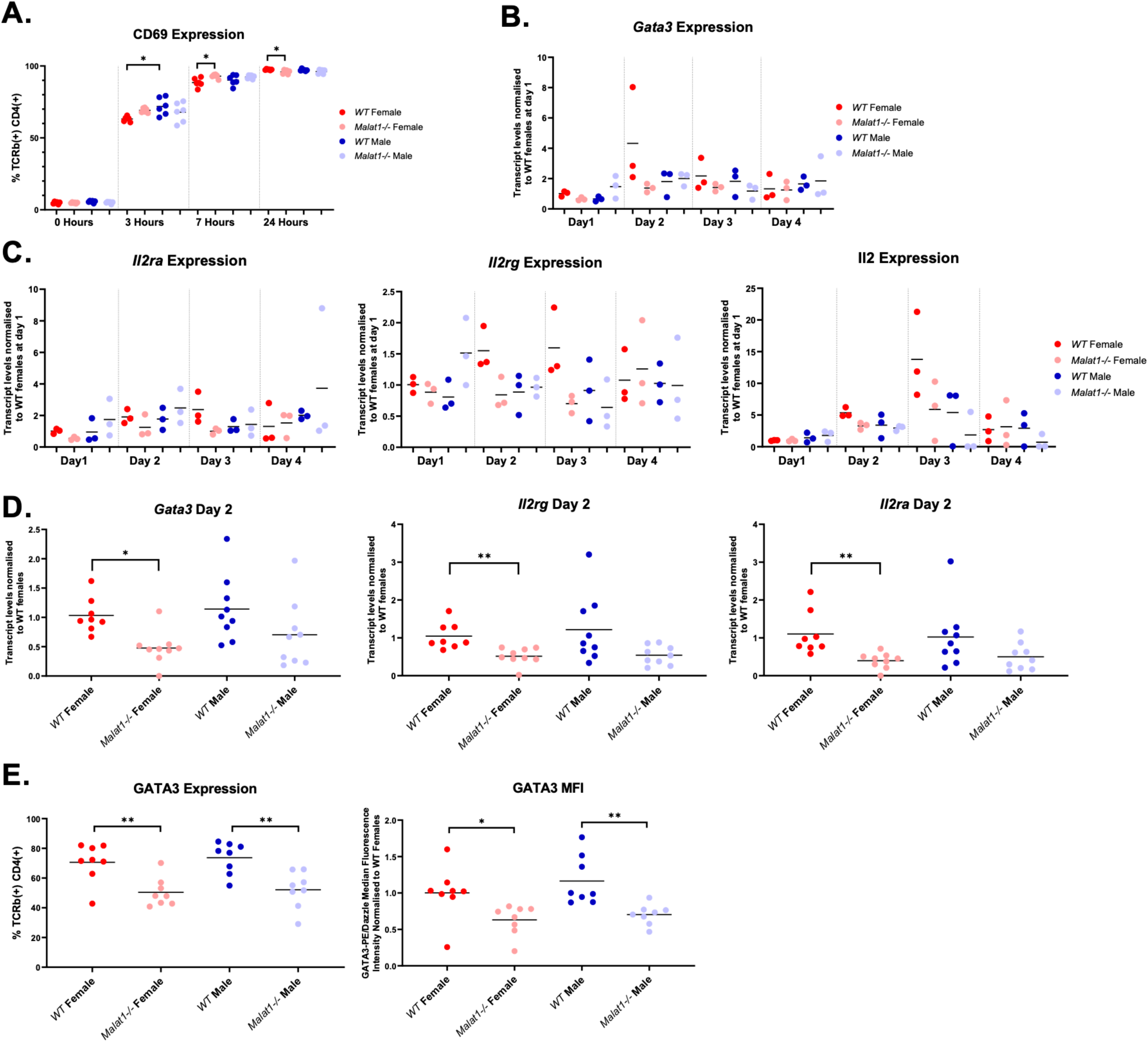
*Malat1* loss enhances naïve CD4^+^ T cell activation and impairs early Th2 differentiation. **A.** Percentage of CD69^+^ live TCRb^+^ CD4^+^ *WT* or *Malat1^-/-^ in vitro* activated CD4^+^ T cells derived from female or male mice at 0, 3, 7, and 24 hours post-activation. Levels determined by surface staining (n=6). **B.** *Gata3* mRNA levels at days 1, 2, 3 and 4 of Th2 differentiation determined by qRTPCR (n=3). Levels normalised to U6 and average levels of *WT* females at day 1. **C.** *Il2ra*, *Il2rg*, and *Il2* mRNA levels at days 1, 2, 3 and 4 of Th2 differentiation determined by qRTPCR (n=3). Levels normalised to U6 and average levels of *WT* females at day 1. **D.** *Il2ra*, *Il2rg* and *Gata3* mRNA levels at day 2 of differentiation determined by qRTPCR (n=9). Levels normalised to *U6* and average levels of *WT* females. **E.** Percentage and Median Fluorescence Intensity (MFI) of GATA3^+^ live TCRb^+^ CD4^+^ *WT* or *Malat1^-/-^* cells derived from female or male mice at day 2 of differentiation (n=5 per condition).

To explore the consequences of the altered activation kinetics in female *Malat1^-/-^* cells, we performed a daily timecourse (days 1 – 4), measuring transcript levels of key transcription factors (*Gata3*, *Maf*, and *Tbet*) and differentiation-associated genes (*Il2, Il2ra, Il2rg*). We focused on *Il2ra* and *Il2rg* as the former is induced during differentiation and the latter is an X-linked gene, given the sex-specificity of the observed effects. We noticed suppression for multiple of the measured transcripts in female *Malat1^-/-^* differentiating cells, including *Il2*, *Il2ra*, *Il2rg*, and *Gata3* on days 2 and 3 of differentiation (**Fig. 3B – C,** and **Supplemental Fig. S3E**). These initial findings were confirmed in independent experiments for *Il2ra*, *Il2rg*, and *Gata3* on day 2 (**Fig. 3D**). We chose day 2 for the validation experiments as this was the earliest timepoint the suppression of the Th2 programme was observed. In male *Malat1^-/-^* cells, we observed higher variability and similar but non-statistically significant trends (**Fig. 3D**). GATA3 protein levels were suppressed in *Malat1^-/-^* cells at day 2 of the differentiation in both male and female *Malat1^-/-^* cells (**Fig. 3E** and **Supplemental Fig, S3F**). At this point we did not find any significant differences in the levels of the IL2R subunits (**Supplemental Fig. S3G**). These results suggested that *Malat1* was necessary for appropriate early Th2 differentiation.

### Impairment of early differentiation in *Malat1^-/-^* cells is characterised by up-regulation of an ISG signature that suppresses differentiation

To further explore the role of *Malat1* in early Th2 differentiation we performed long-read RNAseq of *WT* and *Malat1^-/-^*naïve CD4^+^ T cells and differentiating cells at day 2 of differentiation. As *Malat1* has been suggested to be involved in regulation of mRNA splicing (14), we opted for long-read RNAseq to facilitate concurrent assessment of differential gene expression and differential transcript usage (DTU). As in the case of our short read RNAseq analysis, there were only few differences between *WT* and *Malat1^-/-^*naïve CD4^+^ T cells (16 DEGs at FDR<0.1, **Supplemental Fig. S4A** and **Supplemental Table S6**). However, comparison of *WT* and *Malat1^-/-^* cells on day 2 of differentiation revealed 239 DEGs, 199 of which were up-regulated in *Malat1^-/-^* cells (**Fig. 4A** and **Supplemental Table S7**). Gene set enrichment analysis (GSEA; (29)), revealed a profound over-representation of interferon alpha and interferon gamma response gene signatures amongst the up-regulated genes in *Malat1^-/-^* cells on day 2 (**Fig. 4B**), whereas enriched terms amongst down-regulated genes, including hypoxia, only reached modest significance (**Supplemental Fig. S4B**). We note that in agreement with our qRTPCR analysis (**Fig. 3D**), trends towards down-regulation were observed for *Il2ra* (LFC = -0.589) and *Gata3* (LFC = -0.401) but these did not reach statistical significance. The most prominent feature of *Malat1^-/-^* cells, the up-regulated interferon gene cluster (**Supplemental Fig. S4C** and **Supplemental Table S8**; clusters identified by k-means clustering) included ISGs (e.g. *Ifit1bl1*, *Ifit3*, *Ifit1*, *Oas2*, and others) and transcription factors including *Irf7*, *Irf1*, and *Stat1* (**Fig. 4C**, **Supplemental Fig. S4C** and **Supplemental Tables S7 and S8**). Most of the day 2 *Malat1*-associated DEGs were down-regulated during differentiation of *WT* cells both when comparing day 2 with naïve cells (**Fig. 4D**) and when comparing *WT* Th2 and naïve CD4^+^ T cells using our short-read sequencing data (**Fig. 4E**). This was also the case specifically for the ISG module (**Supplemental Fig. S4D**). As in the case of Th2 cells (**Fig. 1G**) *Malat1* deficiency had a transcriptome-wide effect by blunting changes in gene expression occurring in *WT* cells within the first two days of differentiation (**Supplemental Fig. S4E**).

**Figure 4.**
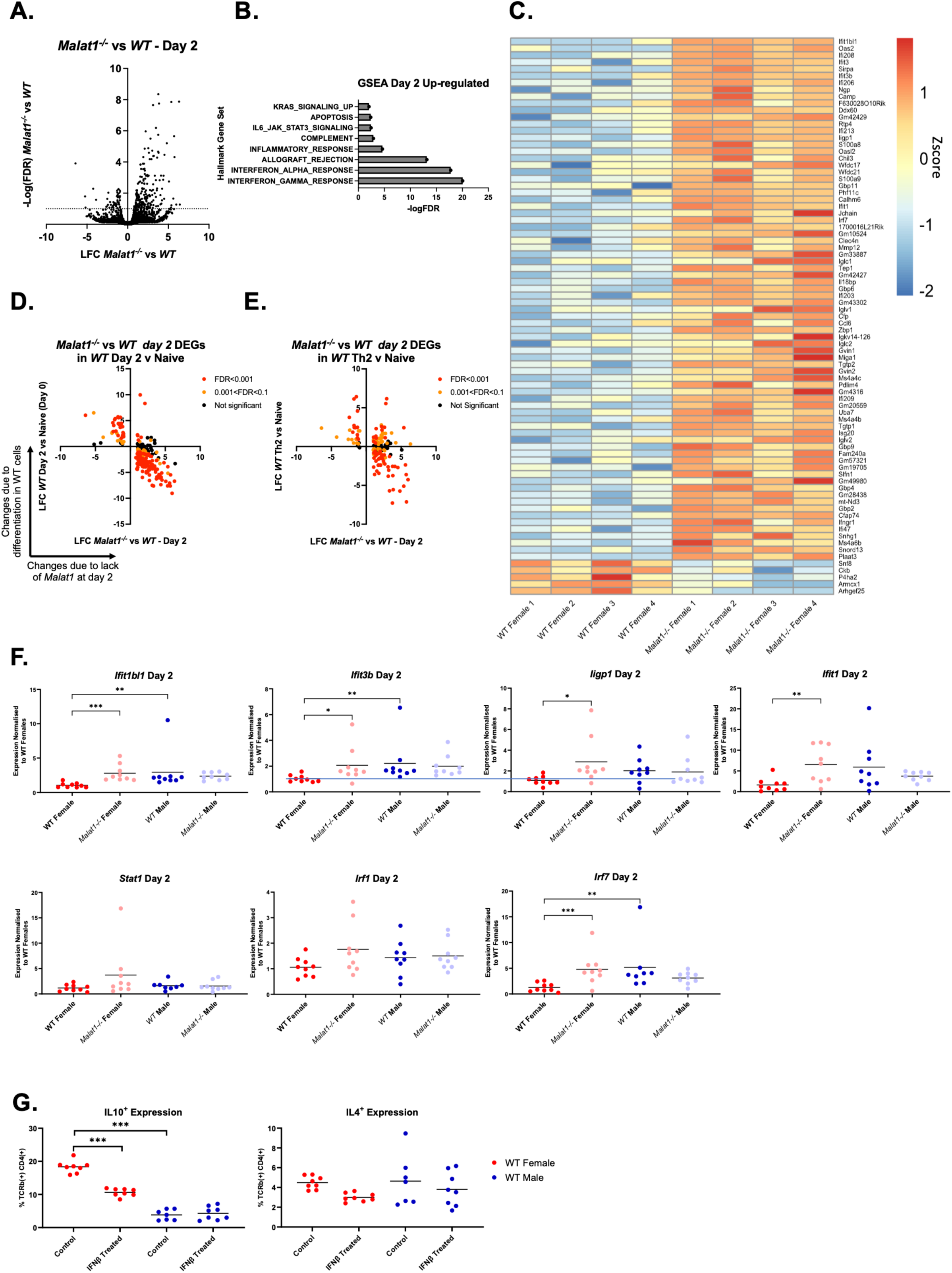
Impaired suppression of an interferon gene module during early differentiation of female *Malat1^-/-^*Th2 cells. A. Volcano plot displaying LFC in gene expression between *WT* and *Malat1^-/-^*female cells (determined by Nanopore long-read RNAseq, N=4 per group) against -logFDR at day 2 of *in vitro* differentiation. B. Gene set enrichment analysis (GSEA) hallmark gene set terms enriched within significantly up-regulated genes in *Malat1^-/-^* cells at day 2 of *in vitro* differentiation. C. Heatmap displaying Z score of log2CPM for top DEGs (FDR<0.01) between *WT* and *Malat1^-/-^* cells at day 2 of differentiation, in descending order of LFC. D. LFC in gene expression between female *WT* naïve and differentiating cells at day 2 of *in vitro* differentiation against LFC between *WT* and *Malat1^-/-^* female cells at day 2 of *in vitro* Th2 differentiation. Data shown for all statistically significant DEGs between *WT* and *Malat1^-/-^* cells at day 2. E. LFC in gene expression between female *WT* naïve and Th2 cells (as in Figure 1) against LFC between *WT* and *Malat1^-/-^* female cells at day 2 of *in vitro* Th2 differentiation. Data shown for all statistically significant DEGs between *WT* and *Malat1^-/-^* cells at day 2. Colours indicate level of significance in differential expression when comparing day 2 vs naïve CD4^+^ T cells (left) or Th2 vs naïve CD4^+^ T cells (right). F. Levels of interferon induced genes (*Ifit1bl1*, *Ifit3b*, *Iigp1*, *Ifit1*) and transcription factors (*Stat1, Irf1, Irf7*) in *WT* and *Malat1^-/-^ in vitro* differentiated female or male cells at day 2 of differentiation. mRNA levels determined by qRTPCR (N=9) and normalised to *U6* and average levels of *WT* cells from female mice. G. Percentage of IL10^+^ and IL4^+^ live TCRb^+^ CD4^+^ *in vitro* differentiated Th2 cells derived from *WT* or *Malat1^-/-^* female and male mice at day 6, treated with 0 (control) or 5000 U/ml IFNβ. Levels determined by intracellular cytokine staining (N=8 for all conditions, except control treated *WT* male, which is n=7).

Validation of the long-read sequencing data by qRTPCR for selected genes confirmed up- regulation of *Ifit1bl1*, *Ifit3b*, *ligp1*, *Ifit1*, and *Irf7* (**Fig. 4F**). Up-regulation trends for Irf1 and Stat1 did not reach statistical significance. Notably, no effects due to *Malat1* deficiency were observed in male CD4^+^ T cells, and levels of *Ifit1bl1*, *Ifit3b*, and *Irf7* were significantly higher in male *WT* cells compared to female cells on day 2 (**Fig. 4F**). Comparing day 2 to naïve CD4^+^ T cells, we found significant down-regulation for *Ifit1bl1*, *Ifit1*, and *Irf7* for all conditions.

Statistically significant *Stat1* down-regulation on day 2 was only observed for female *WT* cells (**Supplemental Fig. S4F**). *Malat1* deficiency had weak effects on DTU affecting only 25 genes (FDR<0.1, **Supplemental Table S9**) including some cytokine genes, but demonstrating modest effects. In some instances, as for *Kdm2a* and *Trim12a* isoform usage changes due to *Malat1* loss showed opposing patterns to those induced by early differentiation of *WT* cells (**Supplemental Fig. S5**).

To explore the link between sustained early ISG expression and endpoint Th2 differentiation, we treated *WT* or *Malat1^-/-^*, female or male, naïve CD4^+^ T cells with IFNβ. This resulted in enhanced mRNA levels in *WT* female-derived cells of several genes identified as up-regulated in *Malat1^-/-^* CD4^+^ T cells at Day 2 of differentiation including *Ifit1bl1*, *Ifit3b*, and *Irf7* (**Supplemental Fig. S6A**). We note that although overall ISG expression trends were similar, there were differences between IFNβ-treated and *Malat1^-/-^*cells at day 2, for example upregulation of *Stat1* reached statistical significance in IFNβ-treated cells (**Supplemental Fig. S6A**) but not when comparing *Malat1^-/-^* to *WT* cells (**Fig. 4F**). Critically, IFNβ treatment resulted in a statistically significant suppression of IL10 expression and a modest decrease in IL4 expression in Th2 cells (**Fig. 4G**) phenocopying the effect of *Malat1* loss. No effects were observed in male-derived cells due to IFNβ treatment (**Fig. 4G** and **Supplemental Fig. S6B**). Interestingly, IL10R blockade enhanced the effect of IFNβ treatment on IL13 expression (higher statistical significance) in female cells but had no effect on IL4 (**Supplemental Fig. S6C**). The latter can be due to the fact that the IFNβ treatment-induced ISG signature only partially overlaps with that seen in *Malat1^-/-^*cells (**Supplemental Fig. S6A**), likely resulting in common but also distinct effects on endpoint cytokine expression. Of note the higher ISG expression in female *Malat1^-/-^* cells was not due to differences in levels of type 1 interferons (**Supplemental Fig. S7A;** *Ifna*, *Ifnb*, and *Ifna4* were lowly detectable at mRNA level only on day 2). We did not observe any differences in *Ifnar1* mRNA levels (**Supplemental Fig. S7B**). IFNAR1 protein expression was higher in Day 2 differentiating CD4^+^ T cells compared to naïve CD4^+^ T cells but we did not observe any differences between female and male and *WT* and *Malat1^-/-^* cells (**Supplemental Fig. S7C**) that could account for the observed differences in ISG expression. We only found a modest increase in IFNAR1 expression in male *Malat1^-/-^* naïve cells compared to *WT* controls but this was not followed by any changes in ISG expression (**Fig. 4F and Supplemental Fig. S4**).

Overall, these analyses further supported that *Malat1* is necessary for early Th2 differentiation with the predominant effect being impaired suppression of an ISG signature that is associated with impaired Th2 differentiation.

### Female differentiating Th2 cells are sensitive to suppression of IL2R caused by *Malat1* deficiency

Having shown that *Malat1* deficiency impairs early Th2 differentiation of CD4^+^ T cells, we reasoned that this could affect their readiness to respond to exogenous IL2. Having found decreases in the mRNA levels of the IL2R subunits (**Fig. 3D**), we measured protein levels of IL2Rα and IL2Rγ on day 4, prior to addition of exogenous IL2. We found that IL2Rα was statistically significantly down-regulated only in female *Malat1^-/-^*T cells on day 4 (**Fig. 5A**, **Supplemental Fig. S8A**). IL2Rγ was down-regulated in both female and male *Malat1^-/-^* T cells (**Fig. 5B**, **Supplemental Fig. S8B**). No differences were observed at the mRNA level at this timepoint (**Supplemental Fig. S8C**). Similarly, IL2Rα was significantly down-regulated on day 4 in female cells treated with *Malat1*-targeting GapmeRs (**Fig. 5C**) and this was also the case for IL2Rγ (**Fig. 5D**). In male cells, there was a non-significant trend towards down-regulation for IL2Rγ (**Fig. 5D**). To explore the link between the early *Malat1-*associated ISG signature and IL2R subunit expression on day 4, we treated *WT* cells with IFNβ. This resulted in a statically significant suppression of IL2Rγ expression and a non-significant decrease in IL2Rα, phenocopying in part the effect of *Malat1* loss or inhibition. IFNβ did not affect IL2R expression in male-derived cells (**Fig. E – F**).

**Figure 5.**
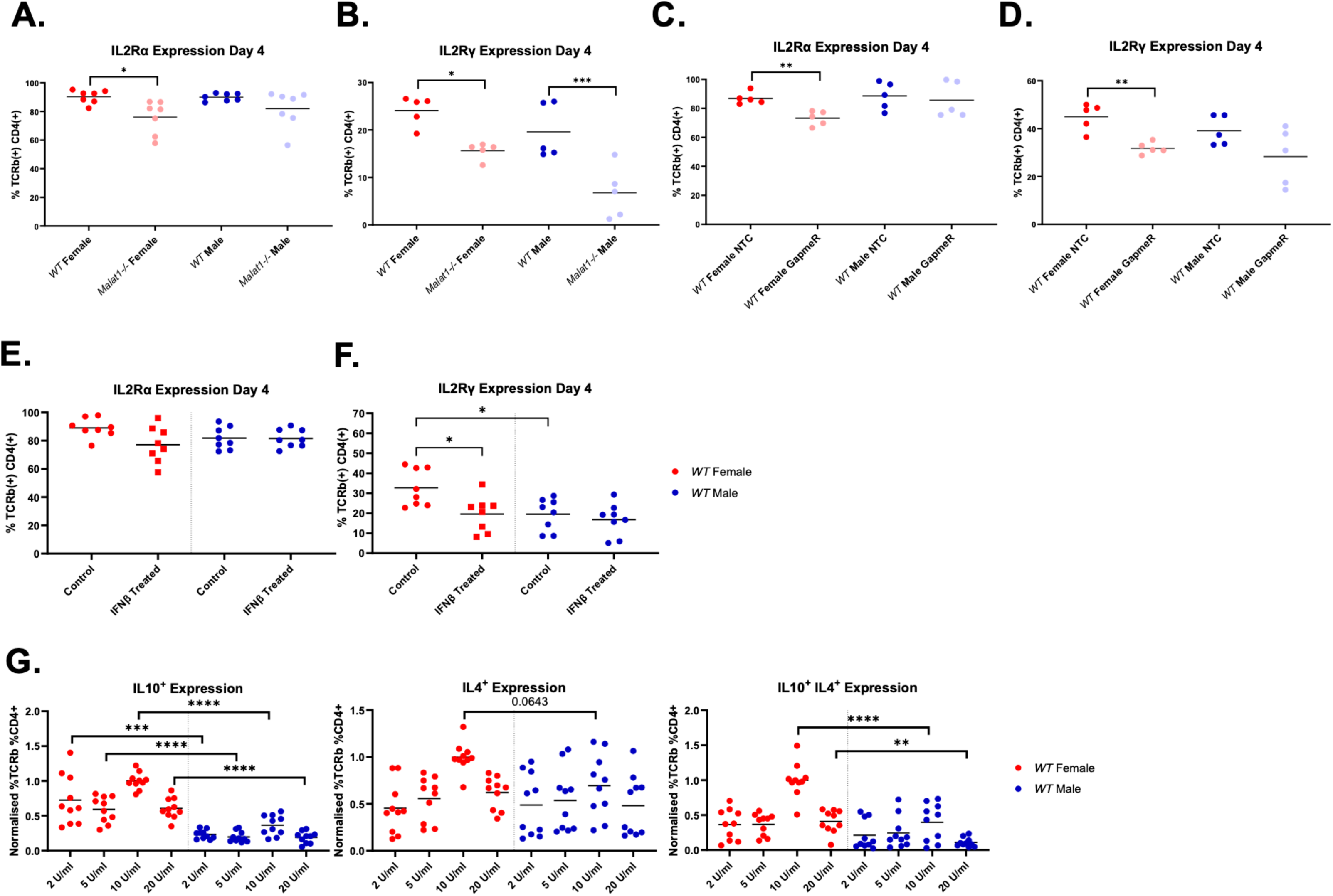
*Malat1* loss decreases expression of the IL2 receptor and impairs the response to exogenous levels of IL2 in cells from female mice. **A.** Percentage of IL2Rα^+^ live TCRb^+^ CD4^+^ *WT* or *Malat1^-/-^* cells derived from female or male mice at day 4 of differentiation (levels determined by surface staining, N=6). **B.** Percentage of IL2Rγ^+^ live TCRb^+^ CD4^+^ *WT* or *Malat1^-/-^* cells derived from female or male mice at day 4 of differentiation (levels determined by surface staining, N=5). **C.** Percentage of IL2Rα^+^ live TCRb^+^ CD4^+^ *WT* cells derived from female and male mice at day 4 of differentiation that were treated with either non-targeting control (NTC) or *Malat1* targeting GapmeRs. Levels determined by surface staining (N=5). **D.** As in **C**, but for IL2Rγ^+^ live TCRb^+^ CD4^+^ cells. **E.** Percentage of IL2Rα^+^ live TCRb^+^ CD4^+^ *WT* cells derived from female and male mice at day 4 of differentiation, treated with 0 (control) or 5000 U/ml IFNβ. Levels determined by surface staining (N=8). **F.** As in **E**, but for IL2Rγ^+^. **G.** Normalised percentage of IL10^+^, IL4^+^ or IL10^+^ IL4^+^ live TCRb^+^ CD4^+^ *WT* or *Malat1^-/-^ in vitro* differentiated Th2 cells derived from female and male mice at day 6, with resuspension at day 4 in the indicated concentrations of IL2. Levels determined by intracellular staining, and normalised to average levels in *WT* female-derived cells treated with 10U/ml IL2 from each experiment (N=10).

The observed down-regulation in the IL2R subunits at this time-point in female *Malat1^-/-^* cells can explain the impaired induction of cytokine expression between day 4 and day 6 (**Fig. 1C – F**). This also indicates that starting from the same baseline at day 4 with regards to IL4 and IL10 expression, *WT* male cells were less responsive to exogenous IL2 compared to female cells. We further tested the dependence of female and male *WT* cells on IL2 stimulation by titrating recombinant IL2. We found that although male cells expand as well or, at the lowest IL2 concentration, better than female cells (**Supplemental Fig. S8D**), they demonstrated lower IL10 expression than female cells at all concentrations (**Fig. 5G**). The observed differential regulation of IL2R receptor and dependence on IL2 for cytokine expression could, at least in part, explain the more profound effect on female Th2 differentiation due to *Malat1* deficiency.

### *Malat1* loss impairs Th2 cell cytokine expression only in female mice in an experimental model of type 2 inflammation

Having demonstrated female-specific effects on Th2 cytokine expression and early differentiation due to *Malat1* deficiency in a cell-intrinsic context *in vitro*, we tested the effect of *Malat1* loss in an experimental model of type 2 inflammation *in vivo*. Intravenous *S. mansoni* egg challenge in mice previously primed with schistosome eggs (**Fig. 6A**) leads to the formation of type 2 inflammatory granulomas in the lungs and mimics helminth-induced lung inflammation (26, 27) leading to cellular infiltration in the lungs, including Th2 cells, and eosinophilia. Indeed, *S. mansoni* egg injection resulted in increased in lung cell numbers, although we note that this only reached statistical significance in *WT* female mice and *Malat1^- -^* male mice (**Supplemental Fig. S9** and **S10A**). Similar splenic cell numbers were observed under all conditions (**Supplemental Fig. S10B**) and similar lung eosinophil numbers and activation, measured by RELMα expression (**Supplemental Fig. S10C – D**). Notably, both lung and splenic CD4^+^ T cells in egg-injected female *Malat1^-/-^* mice expressed lower levels of IL10 than *WT* mice (**Fig. 6B – D**). IL4^+^/IL10^+^ cells were also decreased in the lungs (percentage) and spleens (percentage and numbers) of egg-injected *Malat1^-/-^* mice (**Fig. 6E – F**). This was also the case for triple positive IL4^+^/IL13^+^/IL10^+^ Th2 cells in the lungs of egg-injected mice (**Supplemental Fig. S10E**). None of these differences were observed when comparing male *Malat1^-/-^* mice to *WT* controls. On the contrary the percentage of splenic IL4^+^/IL10^+^ cells increased in egg-injected *Malat1^-/-^* male mice. Under these conditions, we did not observe any differences in total IL4 expression between egg injected *Malat1^-/-^*and *WT* female mice, although we noted an increase in IL4-expressing splenic CD4^+^ T cells in male *Malat1^-/-^* mice compared to *WT* (**Supplemental Fig. S10F – G**). IFNγ^+^ CD4^+^ T cells are also induced as a result of *S. mansoni* egg injection, and IL10 levels were also reduced in female-specific manner within this population in the lungs of inflamed mice (**Supplemental Fig. S10H – I**). With regards to activation status of CD4^+^ T cells, we observed a borderline non-significant decrease in activated (CD44^high^/CD64L^low^) lung CD4^+^ T cells in female *Malat1^-/-^* mice compared to *WT* and no differences in the spleen (**Supplemental Fig. S10J – K**). No differences were observed in percentage and activation status of myeloid cell populations in egg-injected mice (**Supplemental Fig. S10L – M**). Overall, these results indicated that, as observed in our *in vitro* Th2 differentiation experiments, *Malat1* deficiency resulted in a female-specific impairment of Th2 cytokine expression, predominantly demonstrated by reduced IL10 expression in Th2 cells.

**Figure 6.**
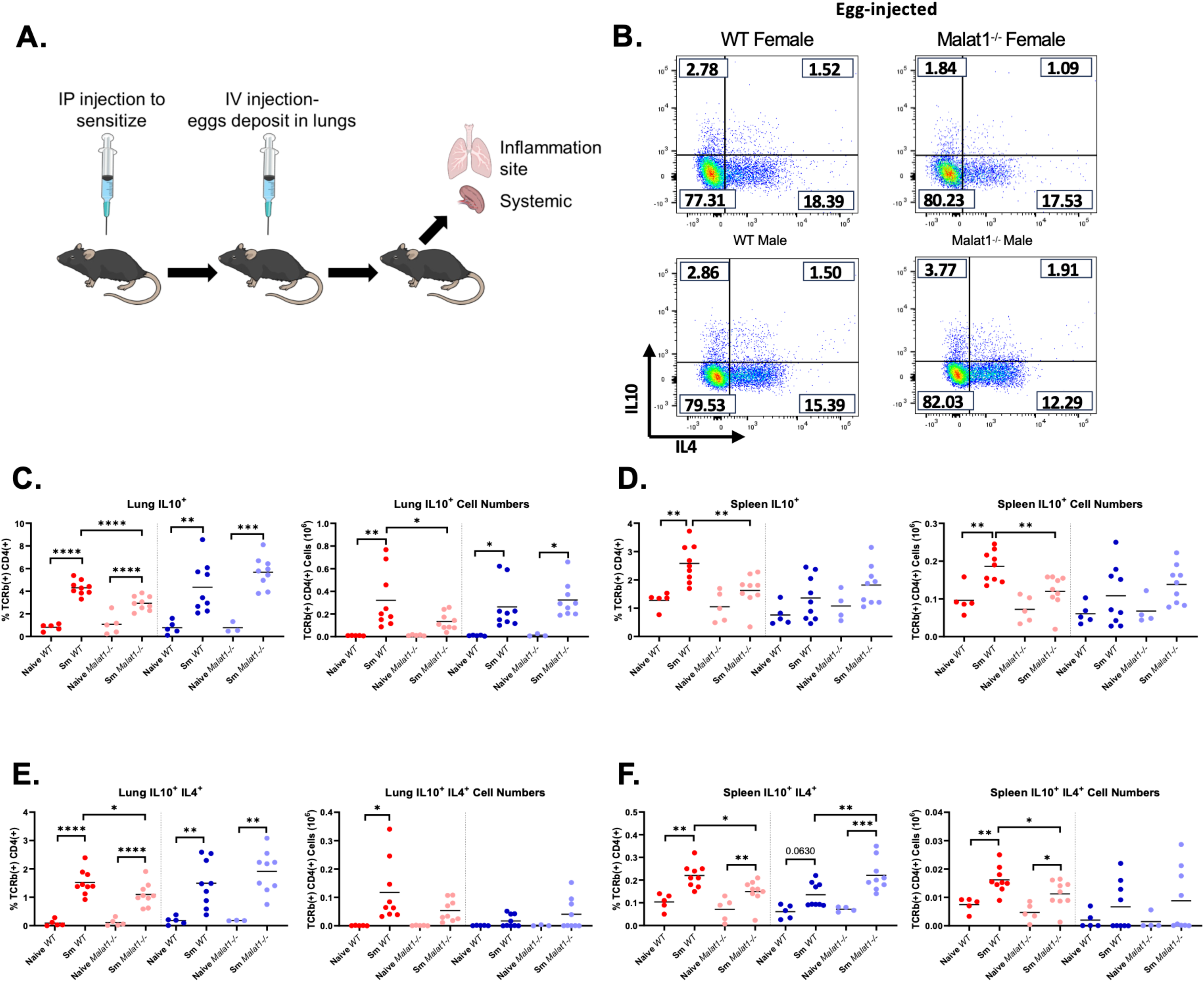
*Malat1* loss causes a female-specific decrease in Th2 cell-derived IL10 levels *in vivo*. **A.** Schematic of *S. mansoni* egg injection experiment. 5,000 dead *S. mansoni* eggs were injected interperitoneally (IP) into mice, then after a week the injection was repeated intravenously (IV) via the tail vein. After another week, lungs and spleens were harvested and processed for FACS. **B.** Representative FACS plots of IL10 and IL4 expression in lung CD4^+^ TCRb^+^ cells from *S. mansoni* egg injected *WT* or *Malat1^-/-^* female or male mice. **C.** Percentage and cell numbers of lung IL10^+^ TCRb^+^ CD4^+^ cells derived from naïve or *S. mansoni* egg injected (*Sm*) *WT* or *Malat1^-/-^,* female and male mice. Levels determined by intracellular staining (N=3 for *Malat1^-/-^*male naïves, N=5 for all other naïves, N=9 for egg injected mice). **D.** As in **C**, but for splenic IL10^+^ TCRb^+^ CD4^+^ cells (N=4 for *Malat1^-/-^* male naïves, N=5 for all other naïves, N=9 for egg injected mice). **E.** Percentage and cell numbers of lung IL10^+^ IL4^+^ TCRb^+^ CD4^+^ cells derived from naïve or *S. mansoni* egg injected (*Sm*) *WT* or *Malat1^-/-^,* female and male mice. Levels determined by intracellular staining (N=3 for *Malat1^-/-^*male naïves, N=5 for all other naïves, N=9 for egg injected mice). **F.** As in **E**, but for splenic IL10^+^ TCRb^+^ CD4^+^ cells (N=4 for *Malat1^-/-^* male naïves, N=5 for all other naïves, N=9 for egg injected mice).

## Discussion

Understanding cell intrinsic mediators of sexual dimorphism in lymphocytes is critical to addressing differences in incidence and severity of immunopathologies between females and males (1–4). We demonstrate that *Malat1*, one of the most highly abundant transcripts in mammalian cells, exerts a sex-specific function in Th2 differentiation, affecting early differentiation and end-point cytokine expression in female cells, predominantly IL10 (**Fig. 7**). *Malat1* deficiency has also some effects in male-derived cells, for example a modest suppression of early GATA3 expression and a suppression in IL2Rγ expression, yet these do not translate in changes in endpoint cytokine expression or gene expression programme. Furthermore, the female-specific effect due to *Malat1* deficiency is observed in the *S. mansoni* egg injection model, as biological sex is a determinant of immune responses in the context of human schistosomiasis (30). We also note that, although *Malat1* loss is associated with suppression of IL10 both *in vitro* and *in vivo*, other effects of *Malat1* deficiency, for example a modest suppression of IL4, are only observed *in vitro*. This can be due to compensatory mechanisms and multiple effects contributing to cytokine expression *in vivo*. For example, through IL10R blockade we found *in vitro* that IL4 and IL13 levels in *Malat1^-/-^* mice are likely to reflect both the loss of *Malat1* and reduced IL10 levels from Th2 cells. This is in agreement with reports demonstrating that IL10 suppresses activation of Th2 cells *in vivo* (31). We speculate that *Malat1* might play a critical role in Th2 differentiation in type 2 immunopathologies, especially those characterised by impaired IL10 signalling (32). In addition, IL10 has been shown to be expressed later than other effector cytokines (33) and from heterogeneous populations of CD4^+^ T cells (34). As such, discrepancies between *in vitro* and *in vivo* cytokine expression can be due to the time-point assessed and kinetics of Th2 cell emergence in the *S. mansoni* egg injection model. Nevertheless, the observed impairment of cytokine expression during *in vitro* Th2 differentiation and in the *S. mansoni* egg injection model strongly support a cell-intrinsic function of *Malat1* in controlling Th2 differentiation in a sex-specific manner.

**Figure 7.**
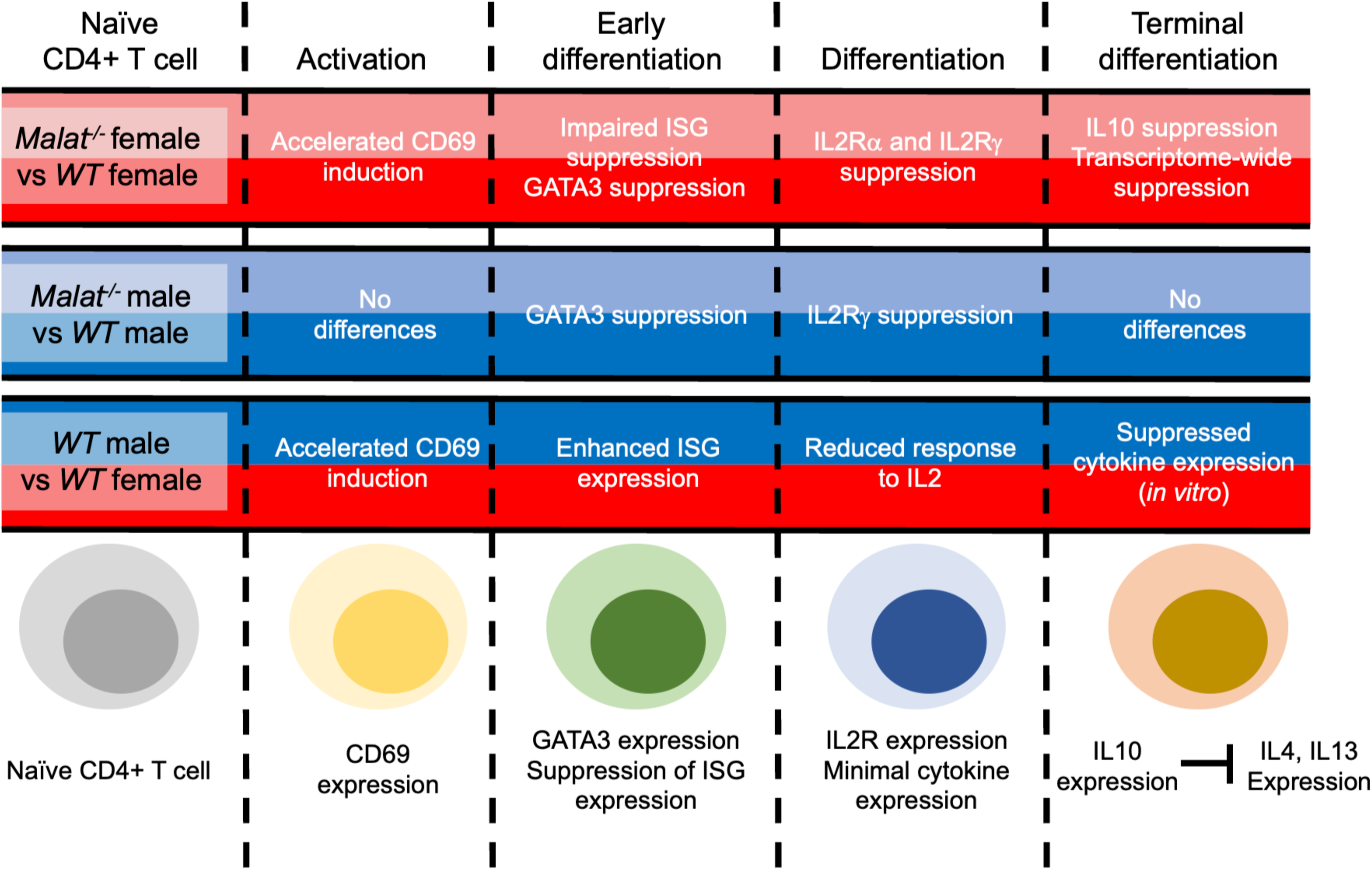
**Schematic summary of the effect of *Malat1* loss on Th2 cell differentiation.** Top row shows the effects observed when comparing *Malat1^-/-^* female-derived cells to *WT*. Second row when comparing male-derived *Malat1^-/-^* and *WT* cells. Third row when comparing male to female-derived *WT* cells. Findings based on the *in vitro* process shown in Fig. 1A.

Despite the limitation of not capturing the complexities of *in vivo* Th2 cell differentiation, *in vitro* differentiation of naïve CD4^+^ T cells to Th2 cells allowed us to demonstrate that *Malat1* loss results in a sex-specific impairment of differentiation in a cell-intrinsic manner. Mechanistically, down-regulation of IL2R upon *Malat1* loss, prior to exposure to exogenous IL2, results in a transcriptome-wide suppression of the Th2 differentiation programme. Although down- regulation of IL2Rγ is observed in both male and female cells, IL2Rα, which is necessary for formation of the high affinity IL2R (35), is only suppressed in female cells. *In vitro*, male Th2 differentiating cells are less responsive to IL2 with regards to cytokine expression, providing an additional potential explanation of the sex-specificity of the observed effect. Evidence for sexual dimorphism in response to IL2 in other immune cell types has been already reported. Female group 2 Innate Lymphoid Cells demonstrate increased proliferation in response to IL2 when compared to male ILC2 cells (36), and female NK cells produce higher levels of IFNγ than males in response to increased levels of IL2 (37).

We postulated that the observed downregulation of IL2Rα and IL2Rγ on day 4 is preceded by impaired early differentiation (days 2-3) of *Malat1^-/-^* cells, an effect which is more pronounced in female cells. To probe how early *Malat1* deficiency impairs Th2 cytokine expression we used long-read RNA sequencing as a superior method in identifying effects on gene expression and, particularly, splicing. *Malat1* deficiency had minimal effects on DTU at day 2 post-differentiation induction. We did not analyse our endpoint (day 6) short read RNA sequencing experiments for DTU because (1) our data showed that it was the effect of *Malat1* on early Th2 differentiation that was driving the phenotype of *Malat1^-/-^* Th2 cells (**Figs. 2 and 3**) and (2) any effects would most likely be a result of impaired response to IL2 (**Fig. 5**) rather than a direct effect of *Malat1*. Indeed, rather than changes in splicing, we found that early differentiation of female *Malat1^-/-^* cells is characterised by impaired suppression of an ISG cluster. It is notable that Th cells demonstrating ISG (including expression of *Ifit3*, *Irf7*, *Stat1*, but not *Ifng*) have been described in type 2 inflammation both in humans (38) and in mouse models (39). In all cases, these Th cells represent a distinct population to Th2 cells indicating that this signature needs to be suppressed in Th2 cells, which in agreement with our findings in *WT* cells. Interestingly a recent publication identified an interferon-experienced population of naïve CD4^+^ T cells that show impaired response to TCR stimulation, as seen here for *Malat1^-/-^* cells (40). Another report demonstrated that these cells exist constitutively including in germ free mice (41). The 50-gene signature that defines interferon-experienced naïve CD4^+^ T cells (40) displays substantial overlap with the *Malat1-*regulated gene cluster (at day 2) identified here (26/50 genes, including *Irf7* and *Stat1*). Taken together with the observed effect of IFNβ on Th2 differentiation here (partially phenocopying of *Malat1* deficiency), this suggests that suppression of this ISG signature in naïve CD4^+^ T cells is necessary for optimal Th2 differentiation and regulated by *Malat1* in female-derived cells. Of note, sexual dimorphism in IFN responses and ISG expression have been reported (42, 43). It will be interesting to further investigate how *Malat1* promotes specifically down-regulation of ISGs in female cells during early CD4^+^ T cell differentiation. It has been shown that *Malat1* can promote gene set-specific Polycomb Repressor Complex-mediated epigenetic silencing in CD8^+^ T cells (24). It could be explored whether this is also the case for ISGs in naive CD4^+^ T cells.

Our results suggest that the impairment of early Th2 differentiation in female *Malat1^-/-^* cells is associated with a stronger but shorter-lived initial activation (0-24h) as measured by CD69 expression. Indeed, strong TCR stimulation in the presence of CD28 co-stimulation has been previously linked to impaired Th2 differentiation (44). During the first 24h of activation, male *WT* cells demonstrate CD69 expression kinetics similar to female *Malat1^-/-^* cells, while deletion of *Malat1* does not cause any further increase in CD69 expression in male cells. Overall, we propose that *Malat1* downregulation upon activation of naïve CD4^+^ T cells facilitates activation, potentially through release of associated proteins that contribute co-transcriptional processing. However, in female cells, complete loss of *Malat1* alters activation kinetics, which in combination with failure to suppress expression of ISGs, leads to impaired Th2 differentiation and subsequent suppression of IL2R and IL2-mediated cytokine expression. Male cells are not sensitive to *Malat1* loss, at least partly, due to their higher early activation threshold, higher ISG expression during early differentiation, and, lower sensitivity to exogenous IL2 during the later stages of differentiation.

We cannot exclude additional female-specific mechanisms being affected by *Malat1*, for example with regards to regulation of X Chromosome-linked genes and the role of X chromosome inactivation during Th2 differentiation (10). Furthermore, it will be essential for our results to be validated in other models of type 2 immunity and in human CD4^+^ T cells. Despite these limitations, our study reveals that female Th2 cells show a specific dependence on *Malat1*, a highly expressed lincRNA not located on a sex chromosome and displaying similar expression in male and female CD4^+^ T cells. This can have far reaching implications for our understanding of immune sexual dimorphism and provide novel routes for sex-specific manipulation of adaptive immunity.

## Author contributions

MG and KAW contributed to experimental design, data generation, data analysis, figure design, and manuscript writing. SvD and KRJ generated and analysed Illumina RNAseq experiments. IK, DC, and SAT contributed to study supervision and data interpretation. JPH contributed to study design, data generation, and data interpretation. DL conceived and coordinated the study, contributed to study design, data interpretation, and study supervision, and wrote the manuscript. All authors contributed to manuscript editing and approved the final manuscript. MG and KAW share the first position in the author list and are listed alphabetically.

## Acknowledgements

This work was funded by the Hull York Medical School (PhD studentship to MG and additional funding to DL), the Biotechnology and Biological Sciences Research Council (BBSRC) White Rose doctoral training partnership (to KAW, reference BB/J014443/1), and the Medical Research Council (to JPH, reference MR/W018578/1). We thank Sally James and Fabiano Pais at the Genomics Lab and Data Science Hub, respectively, in the University of York Bioscience Technology Facility for support with long-read sequencing data. We thank Joanna Greenman for technical support and Allison Green for comments on the manuscript.

## Data availability

Raw data are available upon request. RNAseq data have been deposited in the National Center for Biotechnology (NCBI) Gene Expression Omnibus (GEO) database. Accession numbers: GSE279185 for Illumina RNAseq and GSE278413 for Oxford Nanopore Technologies RNAseq data.

