## Supplemental Figures for "*Malat1* regulates female Th2 cell cytokine expression through controlling early differentiation and response to IL2"

**Supplemental Figures S1-S10**

### *Malat1* regulates female Th2 cell cytokine expression through controlling early differentiation and response to IL2

Mags Gwynne[1, 2, *], Katie A. West[1, 2, 3, *], Stijn van Dongen[4, 5], Dawn Coverley[1, 3], Teichmann[6, 7], S.A., Kylie R. James[8], James P. Hewitson[1, 3], Dimitris Lagos[1, 2].

**1.** Hull York Medical School, University of York, United Kingdom. **2.** York Biomedical Research Institute, University of York, United Kingdom. **3.** Department of Biology, University of York, United Kingdom. **4.** Wellcome Sanger Institute, Wellcome Genome Campus, Hinxton, United Kingdom. **5.** Current address: ExpressionEdits Ltd, Haverhill, United Kingdom. **6.** Wellcome Sanger Institute, Hinxton, United Kingdom. **7.** Current address: Cambridge Stem Cell Institute, University of Cambridge, Cambridge, United Kingdom. **8.** Translational Genomics Program, Garvan Institute of Medical Research, The Kinghorn Cancer Centre, Darlinghurst, New South Wales, Australia.

* These authors contributed equally

#### Supplemental Figure S1

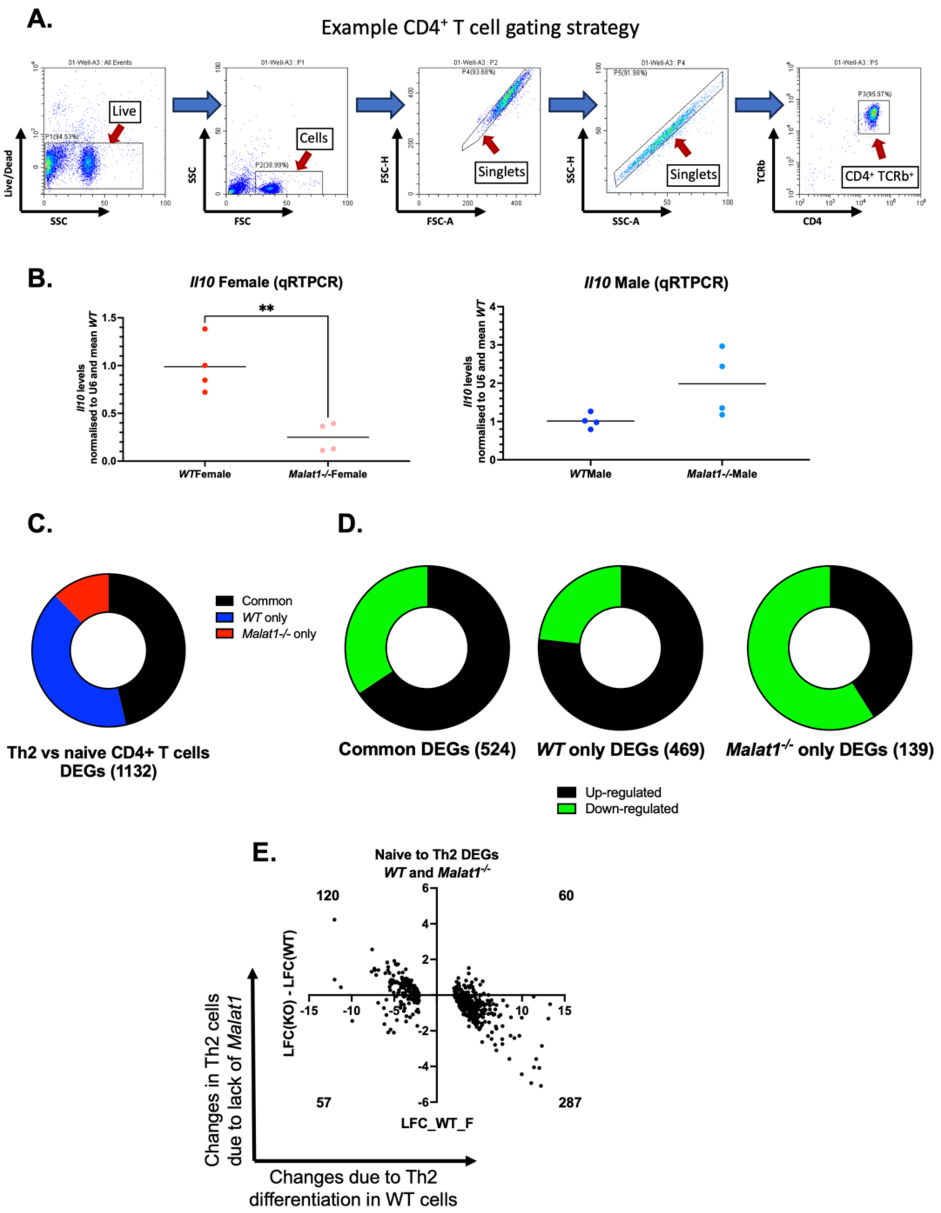


**Supplemental Figure 1. *Malat1* loss impairs *in vitro* Th2 differentiation only in female cells.**

**A.** Example gating strategy for naive CD4+ TCRb+ cells from in vitro culture. **B.** Levels of *Il10* mRNA in the samples used for bulk RNA sequencing in Figure 1. Levels determined by qRTPCR and normalised to *U6* and *WT* average. **C.** Significantly DEGs between naïve and *in vitro* differentiated Th2 cells for both *WT* and *Malat1*$^{-/-}$ cells (black), only *WT* (blue), and only

*Malat1*$^{-/-}$ cells. **D.** Number of up-regulated and down-regulated DEGs between naïve CD4$^{+}$ T and Th2 cells observed for both *WT* or *Malat1*$^{-/-}$ cells or only in one of the two genotypes. **E.** $Log_2$ fold change (LFC) between *WT* CD4$^{+}$ naïve and Th2 cells against the difference in LFC (Th2 – naïve CD4$^{+}$ T cells) between *Malat1*$^{-/-}$ (KO) and *WT* conditions for genes that are statistically significantly deregulated in both *WT* and *Malat1*$^{-/-}$ conditions. The number of genes in each quadrant is also shown.

### Supplemental Figure S2

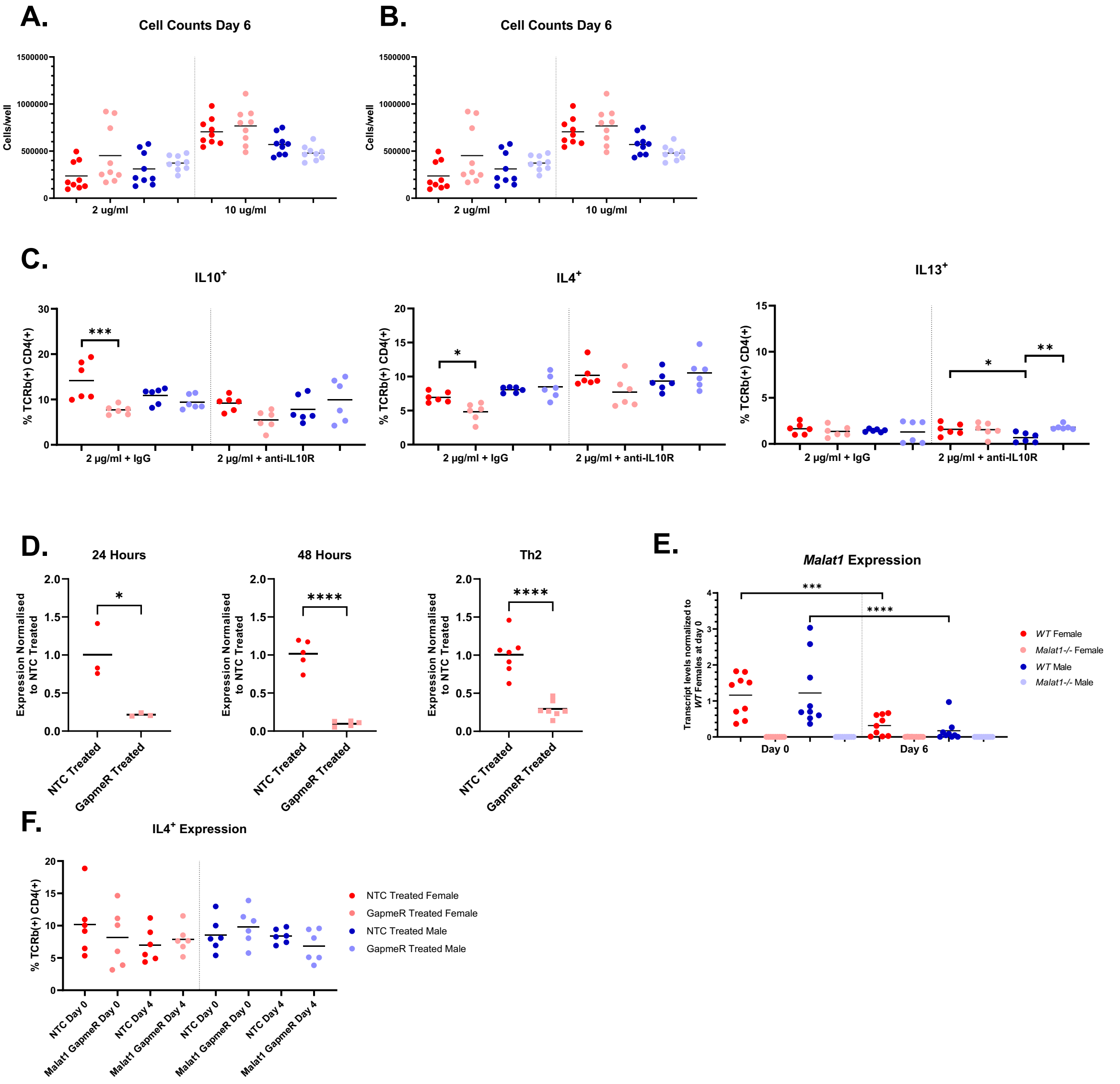

**Supplemental Figure 2. The effect of *Malat1* depletion on IL10 expression is independent of activation strength.**

**A.** Cell counts of live $TCRb^+$ $CD4^+$ *WT* or *Malat1*$^{-/-}$ *in vitro* differentiated Th2 cells at day 4 of differentiation (prior to resuspension in IL2) derived from female and male mice, with standard (10 μg/ml) and 5 times lower (2 μg/ml) levels of anti-CD3 (N=9). **B.** Cell counts of live $TCRb^+$ $CD4^+$ *WT* or *Malat1*$^{-/-}$ *in vitro* differentiated Th2 cells at day 6 of differentiation derived from female and male mice, with strongly activating (10 μg/ml) and weakly activating (2 μg/ml) levels of anti-CD3 (n=9). **C.** Percentage of $IL10^+$, $IL4^+$ and $IL13^+$ live $TCRb^+$ $CD4^+$ *WT* or *Malat1*$^{-/-}$ *in vitro* differentiated Th2 cells derived from female and male mice with 2 μg/ml levels of anti-CD3 antibody, and treatment with IgG or anti-IL10R antibody. Levels determined by intracellular cytokine staining (n=6). **D.** *Malat1* levels at 1 day, 2 days and in differentiated Th2 cells (6 days), following treatment with NTC or *Malat1* targeting GapmeRs on day 0. Levels determined by qRTPCR, and normalised to U6 and average levels of NTC treated GapmeRs. **E.** *Malat1* transcript levels in *in vitro* differentiated Th2 cells at day 0 (naive) and at day 6, determined by qRTPCR. Levels normalised to U6 and average levels of *WT* females at day 0. **F.** Percentage of $IL4^+$ live $TCRb^+$ $CD4^+$ *in vitro* differentiated *WT* Th2 cells derived from female and male mice, treated with either non-targeting control (NTC) gapmers or *Malat1* targeting GapmeRs. Levels determined by intracellular cytokine staining (n=6).

### Supplemental Figure S3

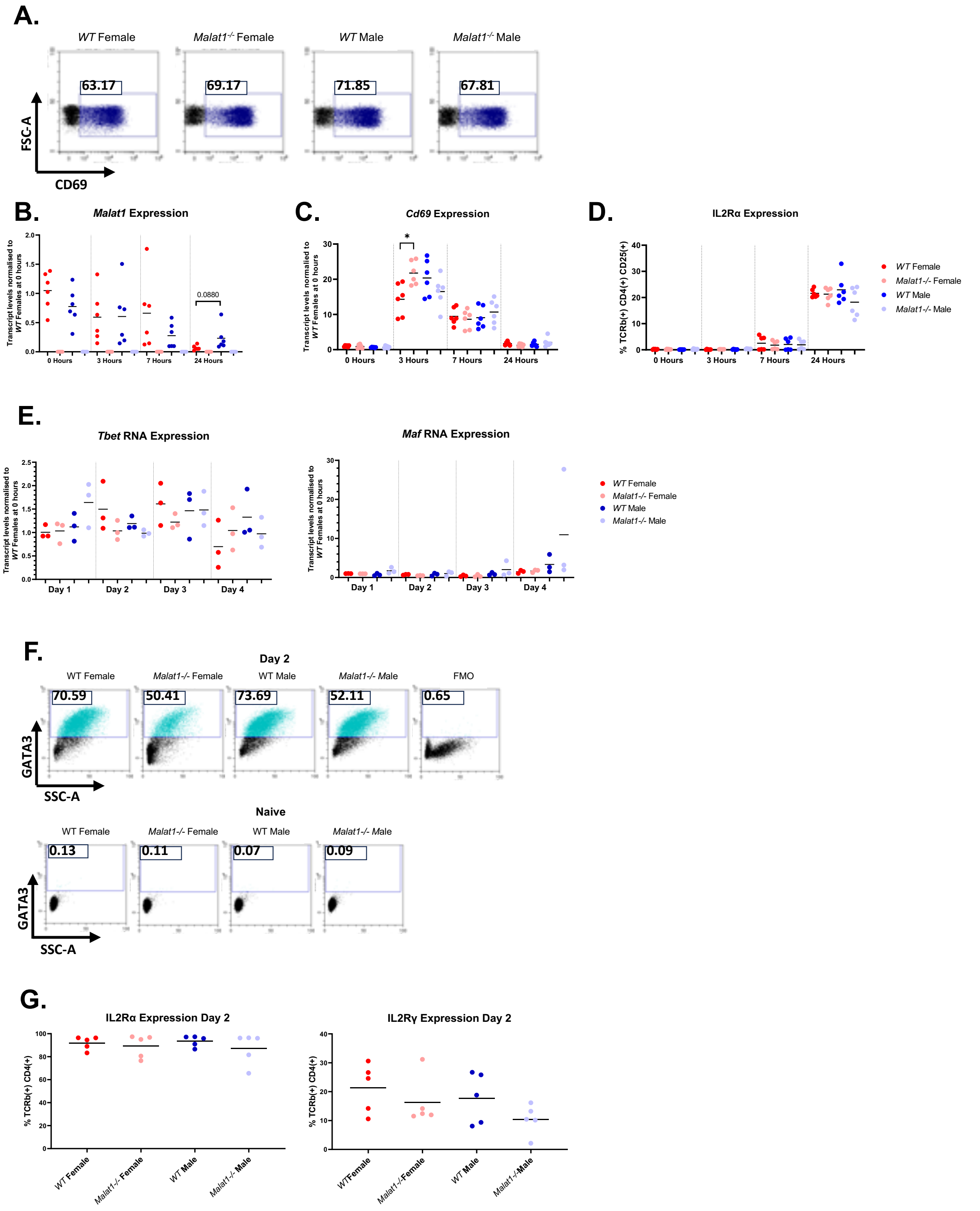


**Supplemental Figure 3. *Malat1* loss alters naïve CD4$^{+}$ T cell activation kinetics and impairs early Th2 differentiation. A.** Representative dot plots displaying CD69 expression in *WT* and *Malat1-/-* mice of both sexes at 3 hours post-activation. **B.** *Malat1* transcript levels in *in vitro* differentiated Th2 cells at 0, 3, 7 and 24 hours post-initial activation, determined by qRTPCR. Levels normalised to U6 and average levels of *WT* females at 0 hours. **C.** *Cd69* transcript levels in *in vitro* differentiated Th2 cells at 0, 3, 7 and 24 hours post-initial activation, determined by qRTPCR. Levels normalised to U6 and average levels of *WT* females at 0 hours. **D.** Percentage of IL2Rα$^{+}$ live TCRb$^{+}$ CD4$^{+}$ *WT* or *Malat1*$^{-/-}$ *in vitro* differentiated Th2 cells derived from female or male mice at 0, 3, 7 and 24 hours post-initial activation. Levels determined by surface staining (n=6). **E.** *Tbet* and *Maf* transcript levels in *in vitro* differentiated Th2 cells at days 1, 2, 3 and 4 determined by qRTPCR. Levels normalised to U6 and average levels of *WT* females at day 1. **F.** Representative dot plots of GATA3 staining in *WT* and *Malat1*$^{-/-}$ in naïve CD4$^{+}$ and differentiating Th2 cells (day 2) (FMO: fluorescence minus one). **G.** Percentage of IL2Rα$^{+}$ or IL2Rg live TCRb$^{+}$ CD4$^{+}$ WT or *Malat1*$^{-/-}$ *in vitro* differentiated Th2 cells derived from female or male mice at 0, 3, 7 and 24 hours post-initial activation. Levels determined by surface staining (n=6).

### Supplemental Figure S4

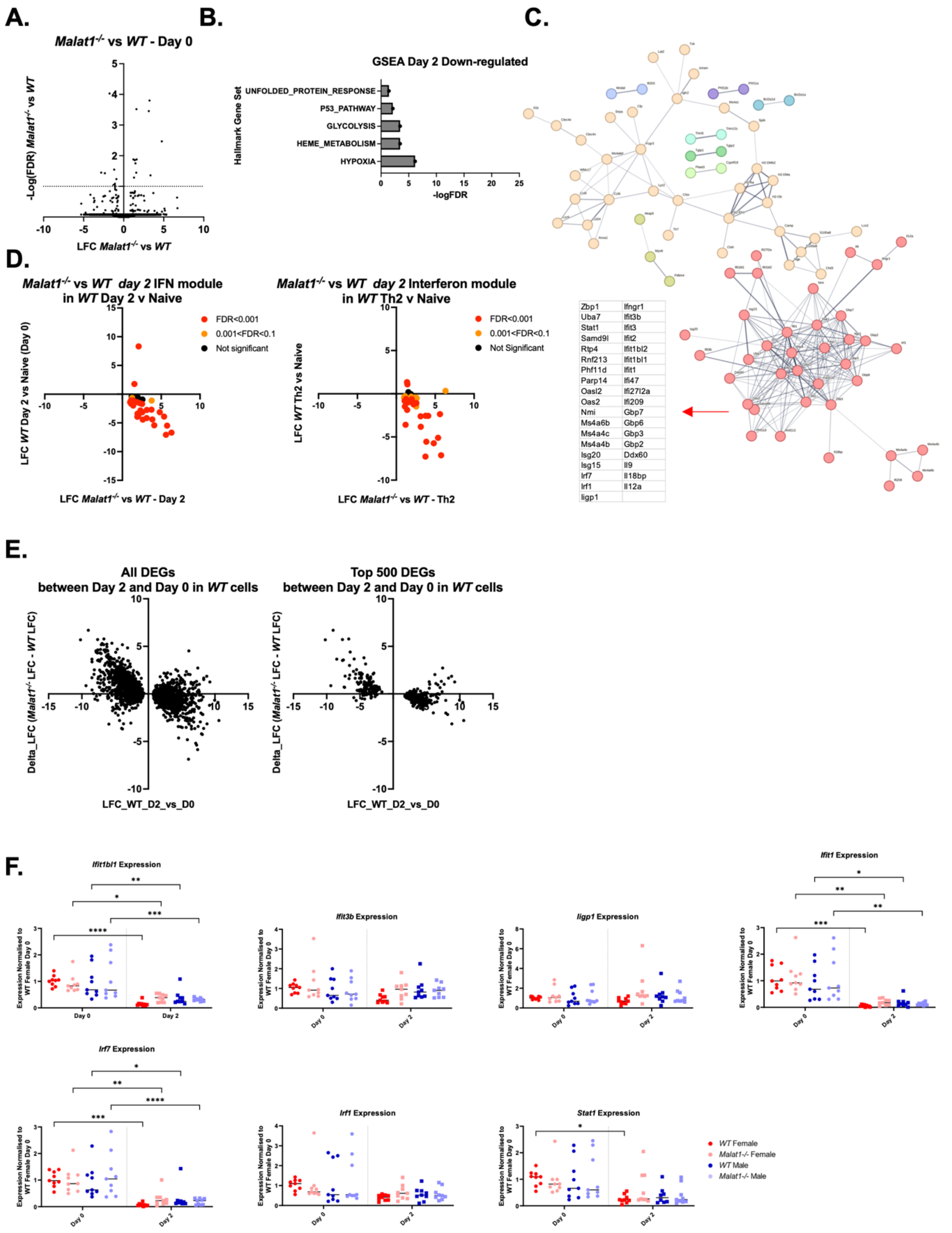


**Supplemental Figure 4. Impaired suppression of an interferon gene cluster during early differentiation of female *Malat1*$^{-/-}$ Th2 cells.**

**A.** Volcano plot displaying LFC in gene expression between *WT* and *Malat1*$^{-/-}$ female naïve CD4$^+$ T cells (Day 0), determined by Nanopore long-read RNAseq, (N=4 per group) against -logFDR at day 2 of *in vitro* differentiation. **B.** Gene set enrichment analysis (GSEA) hallmark gene set terms enriched within significantly down-regulated genes in *Malat1*$^{-/-}$ cells at day 2 of *in vitro* differentiation. **C.** STRING network of significantly up-regulated genes in *Malat1*$^{-/-}$ cells at day 2 of *in vitro* differentiation. Clusters were identified by k-means clustering (k=9). Insert table shows the genes that form part of the interferon gene cluster. **D.** As in main **Fig. 4D** and **E**, but for genes comprising the interferon gene module shown in panel **C** of this figure. Colours indicate level of significance in differential expression when comparing day 2 vs naïve CD4$^+$ T cells (left) or Th2 vs naïve CD4$^+$ T cells (right). **E.** LFC in gene expression between *WT* female naïve and *in vitro* day 2 differentiated *WT* cells against LFC in gene expression between *WT* and *Malat1*$^{-/-}$ cells at the same time point. Data shown for the top 500 and all DEGs between day 2 and naïve CD4$^+$ T *WT* cells. **F.** Levels of interferon induced genes (*Ifit1bl1*, *Ifit3b*, *Iigp1*, *Ifit1*) and transcription factors (*Stat1, Irf1, Irf7*) in *WT* and *Malat1*$^{-/-}$ naïve (Day 0) or *in vitro* differentiated Day 2 female or male cells. mRNA levels determined by qRTPCR (N=9) and normalised to U6 and average levels of naive *WT* cells from female mice. Statistical comparisons between time points shown. Note that data for Day 2 are also shown in **Fig. 4F**.

### Supplemental Figure S5

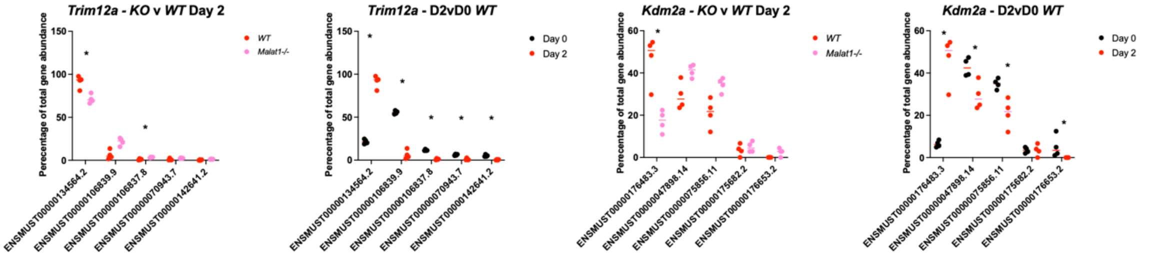


| Tx_ID | Transcript_size | Protein_size | Biotype |
|---|---|---|---|
| ENSMUST00000134564.2 | 3730 | No protein | Retained intron |
| ENSMUST00000106839.9 | 1486 | 297aa | Protein coding |
| ENSMUST00000106837.8 | 1389 | 265aa | Protein coding |
| ENSMUST00000070943.7 | 1531 | 284aa | Protein coding |
| ENSMUST00000142641.2 | 562 | No protein | Protein coding CDS not defined |

| Tx_ID | Transcript_size | Protein_size | Biotype |
|---|---|---|---|
| ENSMUST00000176483.3 | 1128 | 35aa | Protein coding |
| ENSMUST00000047898.14 | 8578 | 1161aa | Protein coding |
| ENSMUST00000075856.11 | 7242 | 1161aa | Protein coding |
| ENSMUST00000175682.2 | 2638 | No protein | Retained intron |
| ENSMUST00000176653.2 | 1076 | 292aa | Protein coding |

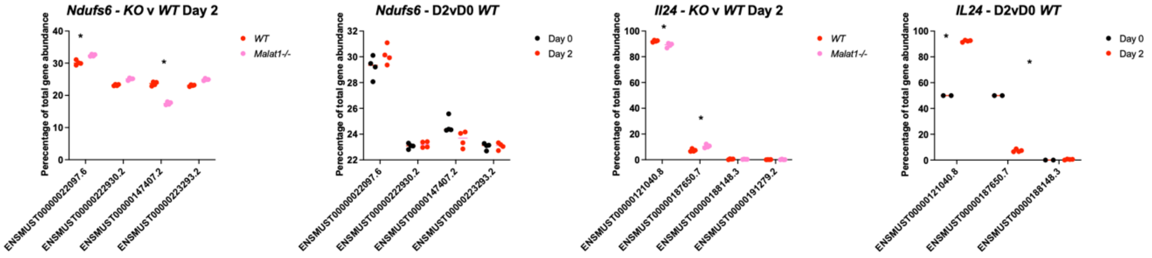


| Tx_ID | Transcript_size | Protein_size | Biotype |
|---|---|---|---|
| ENSMUST00000022097.6 | 568 | 116aa | Protein coding |
| ENSMUST00000222930.2 | 512 | 124aa | Protein coding |
| ENSMUST00000147407.2 | 534 | 48aa | Nonsense mediated decay |
| ENSMUST00000223293.2 | 757 | 98aa | Protein coding |

| Tx_ID | Transcript_size | Protein_size | Biotype |
|---|---|---|---|
| ENSMUST00000121040.8 | 1159 | 181aa | Protein coding |
| ENSMUST00000187650.7 | 1182 | 181aa | Protein coding |
| ENSMUST00000188148.3 | 393 | 77aa | Protein coding |
| ENSMUST00000191279.2 | 634 | 119aa | Nonsense mediated decay |

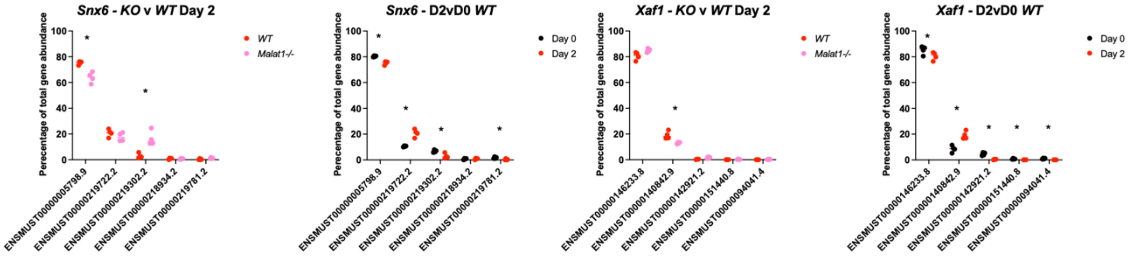


| Tx_ID | Transcript_size | Protein_size | Biotype |
|---|---|---|---|
| ENSMUST00000005798.9 | 1928 | 406aa | Protein coding |
| ENSMUST00000219722.2 | 286 | No protein | Retained intron |
| ENSMUST00000219302.2 | 732 | No protein | Retained intron |
| ENSMUST00000218934.2 | 1441 | 290aa | Protein coding |
| ENSMUST00000219781.2 | 436 | 90aa | Nonsense mediated decay |

| Tx_ID | Transcript_size | Protein_size | Biotype |
|---|---|---|---|
| ENSMUST00000146233.8 | 2597 | 273aa | Protein coding |
| ENSMUST00000140842.9 | 651 | 178aa | Protein coding |
| ENSMUST00000142921.2 | 2582 | No protein | Retained intron |
| ENSMUST00000151440.8 | 672 | 207aa | Protein coding |
| ENSMUST00000094041.4 | 415 | 75aa | Nonsense mediated decay |

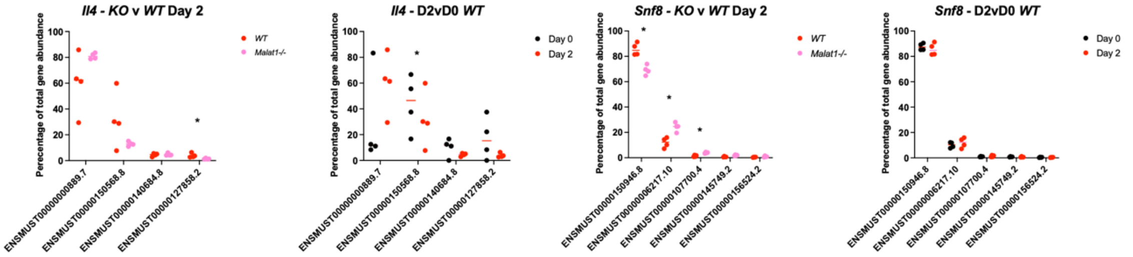


| Tx_ID | Transcript_size | Protein_size | Biotype |
|---|---|---|---|
| ENSMUST00000000889.7 | 590 | 140aa | Protein coding |
| ENSMUST00000150568.8 | 605 | 45aa | Protein coding |
| ENSMUST00000140684.8 | 530 | 45aa | Protein coding |
| ENSMUST00000127858.2 | 541 | No protein | Protein coding CDS not defined |

| Tx_ID | Transcript_size | Protein_size | Biotype |
|---|---|---|---|
| ENSMUST00000150946.8 | 484 | No protein | Protein coding CDS not defined |
| ENSMUST00000006217.10 | 1004 | 258aa | Protein coding |
| ENSMUST00000107700.4 | 849 | 233aa | Protein coding |
| ENSMUST00000145749.2 | 1764 | No protein | Protein coding CDS not defined |
| ENSMUST00000156524.2 | 665 | No protein | Protein coding CDS not defined |

**Supplemental Figure 5. *Malat1* loss has modest effects on differential transcript usage.** Transcript usage as a percentage of total gene abundance for indicated genes that demonstrate statistically significant DTU. Levels shown between *WT* and *Malat1*$^{-/-}$ females at day 2 of *in vitro* differentiation (left graphs) or between naïve (day 0) and day 2 in *WT* cells (right graphs). Levels determined by long-read RNAseq (N=4 per condition). Tables indicate the product of each isoform. Stars indicate isoforms demonstrating statistically significant differences in abundance as determined by Dexseq.

### Supplemental Figure S6

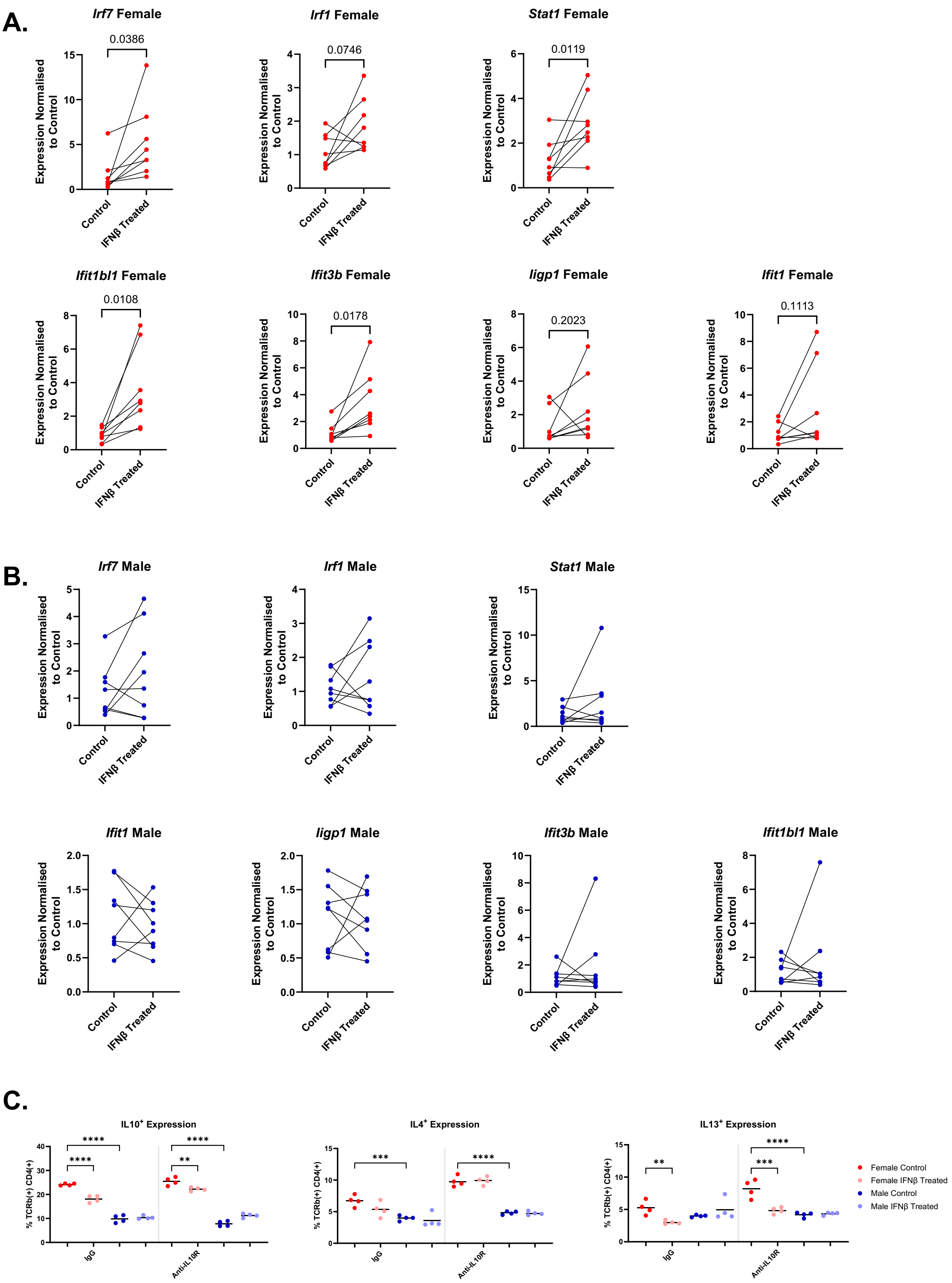


**Supplemental Figure 6. Type-I IFN treatment causes an increase in female IFN-stimulated genes at during early activation.**

**A.** Levels of interferon induced transcription factors (*Stat1, Irf1, Irf7*) and genes (*Ifit1bl1*, *Ifit3b*, *Iigp1*, *Ifit1*) in *WT* female Th2 cells at day 2 of differentiation, with treatment of 0 or 5000 U/ml IFNβ. mRNA levels determined by qRTPCR (N=6) and normalised to *U6* and average levels for the 0 U/ml treatment. **B.** As in **A**, but for male Th2 cells. **C.** Percentage of IL10$^{+}$, IL4$^{+}$, and IL13$^{+}$ live TCRb$^{+}$ CD4$^{+}$ *in vitro* differentiated Th2 cells derived from WT or *Malat1*$^{-/-}$ female and male mice at day 6, treated with or without 5000 U/ml IFNβ, and with either IgG or anti-IL10R. Levels determined by intracellular cytokine staining (N=4 for all conditions).

#### Supplemental Figure S7

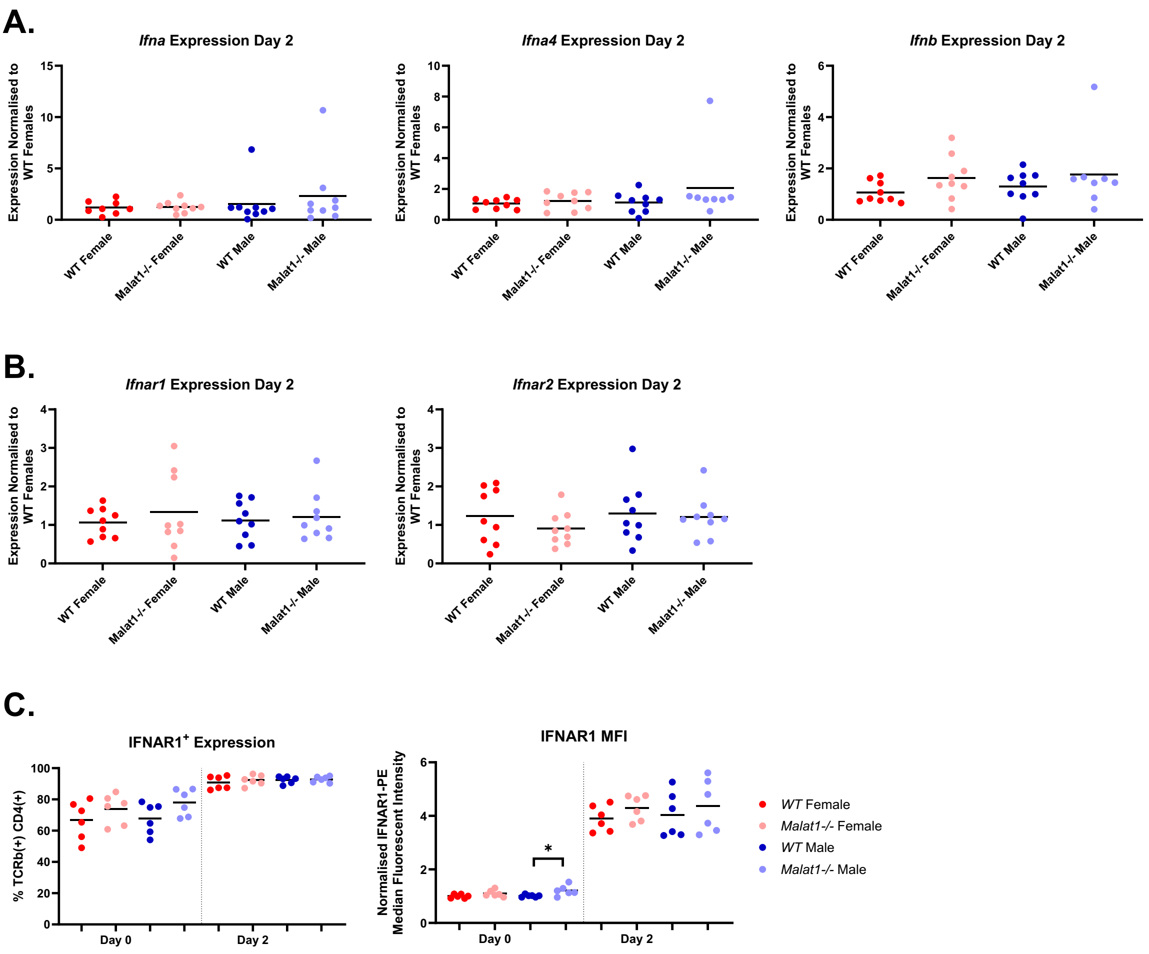


**Supplemental Figure 7. *Malat1* loss does not affect type I IFN and IFNAR1 expression.**

**A.** *Ifna*, *Ifna4* and *Ifnb* transcript expression at day 2 of Th2 differentiation (n=9), determined by qRTPCR. Levels normalised to U6 and average levels of *WT* females. **B.** *Ifnar*1 and *Ifnar2* transcript expression at day 2 of Th2 differentiation (n=9), determined by qRTPCR. Levels normalised to U6 and average levels of *WT* females. **C.** Percentage and normalised median fluorescence intensity of IFNAR1$^{+}$ live TCRb$^{+}$ CD4$^{+}$ *WT* or *Malat1*$^{-/-}$ cells derived from female or male mice at day 0 and day 2 of differentiation (levels determined by surface staining, n=6).

#### Supplemental Figure S8

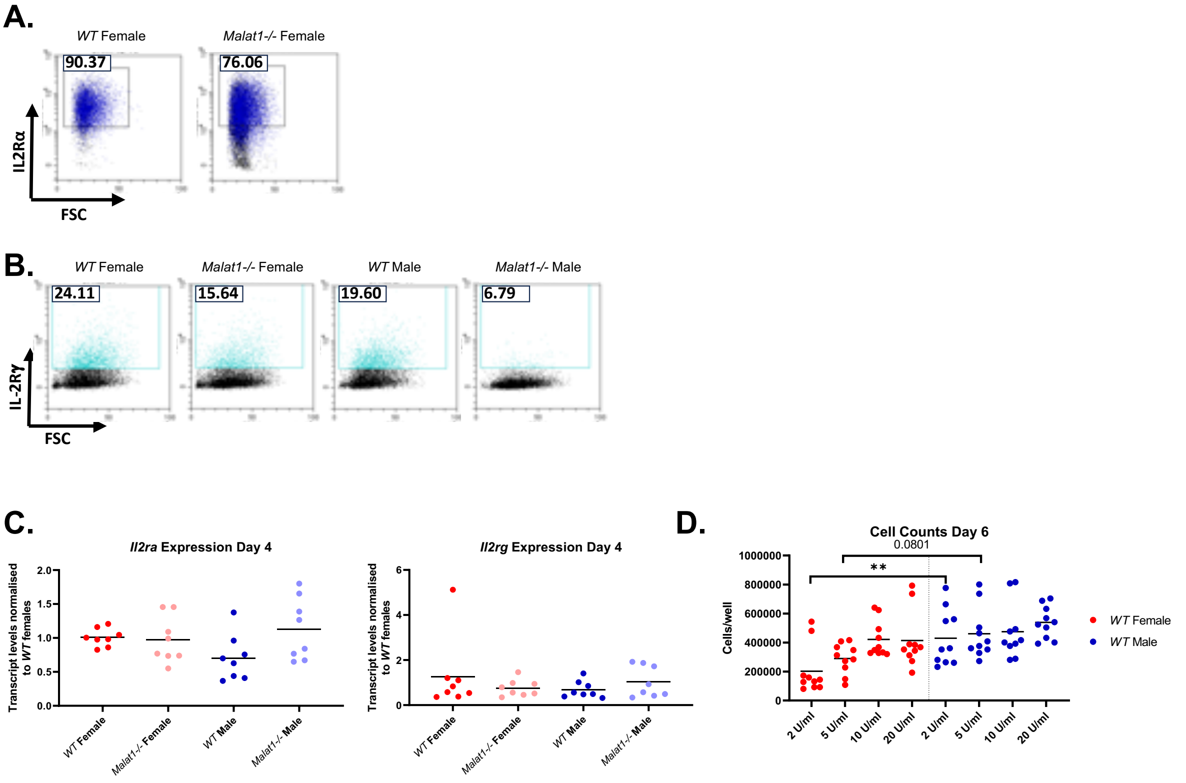


**Supplemental Figure 8. *Malat1* loss does not affect *Il2ra* and *Il2rg* mRNA levels on day 4 of differentiation.**

**A.** Representative dot plots displaying IL2Rα expression in *WT* and *Malat1-/-* female mice at day 4 of Th2 *in vitro* differentiation. **B.** Representative dot plots displaying IL2Rγ expression in *WT* and *Malat1-/-* female mice at day 4 of Th2 *in vitro* differentiation. **C.** *Il2ra* and *Il2rg* transcript levels at day 4 of differentiation in *WT* or *Malat1*$^{-/-}$ CD4$^{+}$ T cells derived from male or female mice, determined by qRTPCR. Levels normalized to U6 and average levels of *WT* females. **D.** Cell numbers of live TCRb$^{+}$ CD4$^{+}$ *WT* or *Malat1*$^{-/-}$ *in vitro* differentiated Th2 cells derived from female and male mice at day 6, with resuspension at day 4 in the indicated concentrations of IL2.

#### Supplemental Figure S9

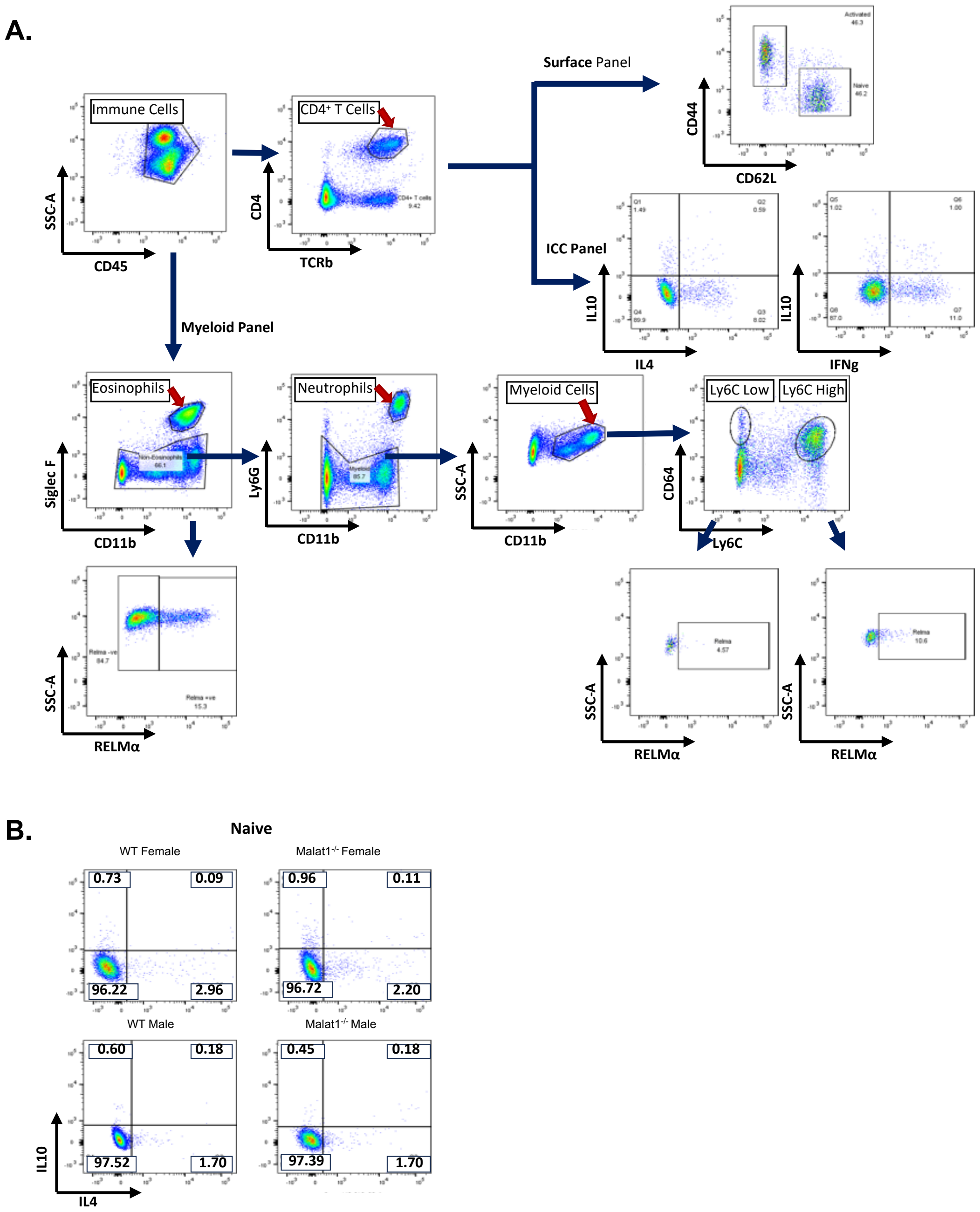


**Supplemental Figure 9. Gating strategy for the *in vivo S. mansoni* egg injection experiment.**

**A.** Gating strategy for the three panels- $CD4^+$ T cell surface panel, $CD4^+$ T cell intracellular cytokine panel, and the myeloid cell panel.

**B.** Representative gating for IL4 and IL10 expression in naïve $CD4^+$ T cells.

### Supplemental Figure S10

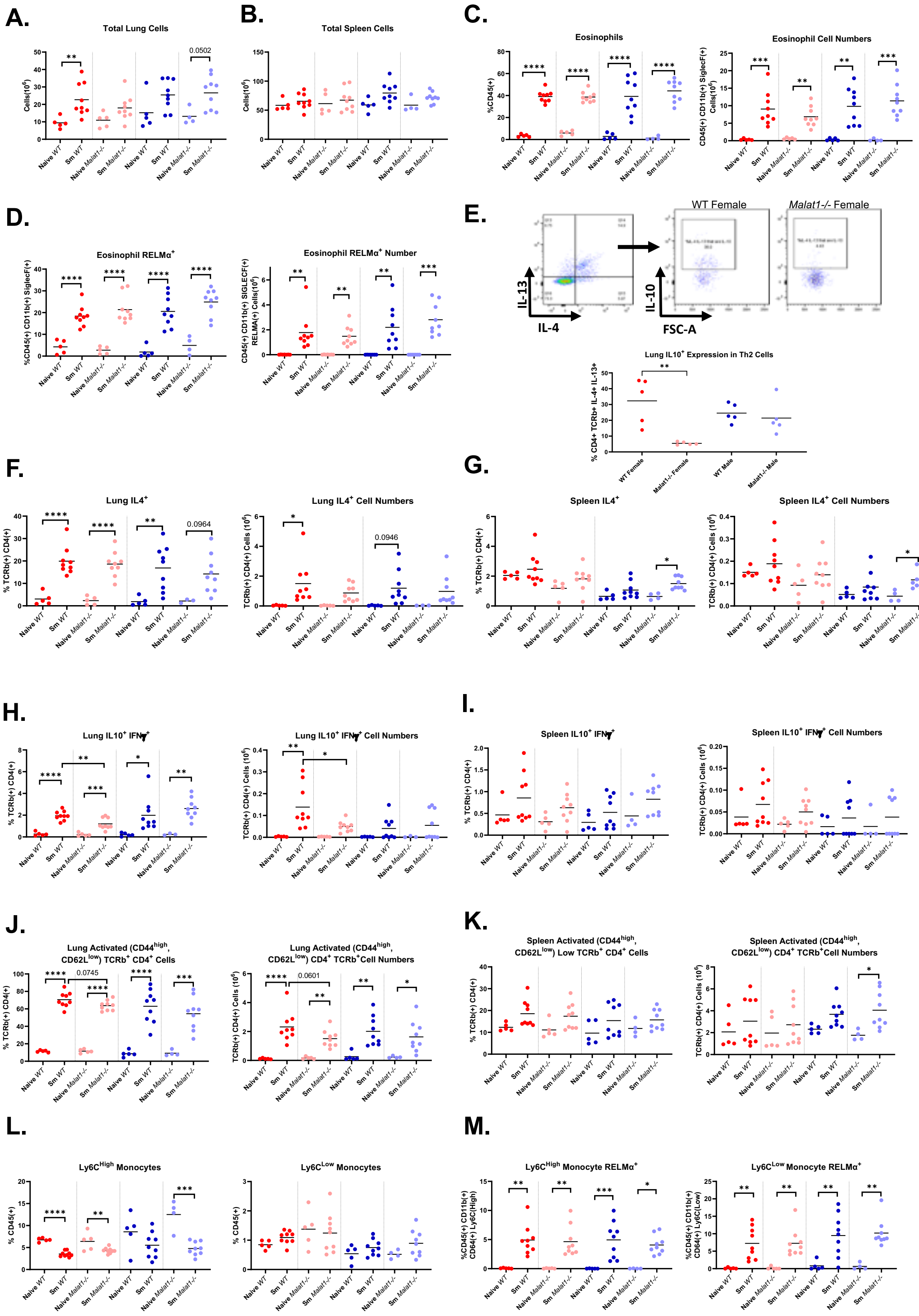

**Supplemental Figure 10. *Malat1* loss causes a female-specific decrease in Th2 cell-derived IL10 levels *in vivo.* A.** Total cell numbers in lungs of *S. mansoni* egg injected (*Sm*) *WT* or *Malat1*$^{-/-}$, female or male mice (N=4 for *Malat1*$^{-/-}$ male naïves, N=5 for all other naïves, N=9 for egg injected mice n=5 for naïve, n=9 for egg injected mice).

**B.** As in **A**, but for splenic cell numbers. **C.** Percentage and cell numbers of eosinophils (CD45$^{+}$, CD11b$^{+}$, SiglecF$^{+}$) in lungs of naïve or *S. mansoni* egg injected, *WT* or *Malat1*$^{-/-}$, female or male mice. Levels determined by surface staining (N=4 for *Malat1*$^{-/-}$ male naïves, N=5 for all other naïves, N=9 for egg injected mice). **D.** Percentage and cell numbers of RELMα expressing lung eosinophils (CD45$^{+}$, CD11b$^{+}$, SiglecF$^{+}$) derived from naïve or *S. mansoni* egg injected, *WT* or *Malat1*$^{-/-}$, female or male mice. Levels determined by surface staining (N=4 for *Malat1*$^{-/-}$ male naïves, N=5 for all other naïves, N=9 for egg injected mice). **E.** Percentages of IL10$^{+}$ IL4$^{+}$ IL13$^{+}$ TCRb$^{+}$ CD4$^{+}$ cells in lungs of *S. mansoni* egg injected, *WT* or *Malat1*$^{-/-}$, female or male mice. Levels determined by surface staining (N=5 for all conditions). Also shown, representative gating strategy displaying IL4 and IL13 staining CD4$^{+}$ TCRb$^{+}$ cells in lungs of *S. mansoni* egg injected mice, and IL10 staining within the IL4$^{+}$ IL13$^{+}$ population for *WT* and *Malat1*$^{-/-}$ females. **F.** Percentage and cell numbers of lung IL4$^{+}$ TCRb$^{+}$ CD4$^{+}$ cells derived from naïve or *S. mansoni* egg injected (*Sm*) *WT* or *Malat1*$^{-/-}$, female and male mice. Levels determined by intracellular staining (N=3 for *Malat1*$^{-/-}$ male naïves, N=5 for all other naïves, N=9 for egg injected mice). **G.** As in **E**, but for splenic IL10$^{+}$ TCRb$^{+}$ CD4$^{+}$ cells (N=4 for *Malat1*$^{-/-}$ male naïves, N=5 for all other naïves, N=9 for egg injected mice). **H.** Percentage and cell numbers of lung IFN$\gamma^{+}$/IL10$^{+}$ TCRb$^{+}$ CD4$^{+}$ cells derived from naïve or *S. mansoni* egg injected (*Sm*) *WT* or *Malat1*$^{-/-}$, female and male mice. Levels determined by intracellular staining (N=3 for *Malat1*$^{-/-}$ male naïves, N=5 for all other naïves, N=9 for egg injected mice). **I.** As in **H**, but for splenic IFN$\gamma^{+}$/IL10$^{+}$ TCRb$^{+}$ CD4$^{+}$ cells (N=4 for *Malat1*$^{-/-}$ male naïves, N=5 for all other naïves, N=9 for egg injected mice). **J.** Percentage and cell numbers of activated (CD44$^{high}$ CD62L$^{low}$) TCRb$^{+}$ CD4$^{+}$ cells in lungs of naïve or *S. mansoni* egg injected, *WT* or *Malat1*$^{-/-}$, female or male mice. Levels determined by surface staining (N=4 for

*Malat1-/-* male naïves, N=5 for all other naïves, N=9 for egg injected mice). **K.** As in **J**, but for splenic CD4$^+$ T cells. **L.** Percentages of Ly6C$^{high}$ and Ly6C$^{low}$ monocytes (CD45$^+$, CD11b$^+$, CD64$^+$) in lungs of naïve or *S. mansoni* egg injected, *WT* or *Malat1*$^{-/-}$*,* female or male mice. Levels determined by surface staining (N=4 for *Malat1*$^{-/-}$ male naïves, N=5 for all other naïves, N=9 for egg injected mice). **M.** Percentages of RELMα expression in Ly6C$^{high}$ and Ly6C$^{low}$ monocytes (CD45$^+$, CD11b$^+$, CD64$^+$) in lungs of naïve or *S. mansoni* egg injected, *WT* or *Malat1*$^{-/-}$*,* female or male mice. Levels determined by surface staining (N=4 for *Malat1*$^{-/-}$ male naïves, N=5 for all other naïves, N=9 for egg injected mice).
