## Supplemental Tables for "*Malat1* regulates female Th2 cell cytokine expression through controlling early differentiation and response to IL2"

**List of Supplemental Tables**

| | |
|---|---|
| **Table S1** | **List of primers for qRTPCR** |
| **Table S2** | **Differentially expressed genes at FDR<0.2 between WT and Malat1-/- cells naïve CD4+ T cells and Th2 cells from male and female mice** |
| **Table S3** | **Differentially expressed genes between Th2 cells and naïve CD4+ T cells at FDR<0.001 and AbsLFC>2, both for WT and Malat1-/- cells from female mice** |
| **Table S4** | **Differentially expressed genes between Th2 cells and naïve CD4+ T cells at FDR<0.001 and AbsLFC>2, reaching significance only for cells from WT mice (female)** |
| **Table S5** | **Differentially expressed genes between Th2 cells and naïve CD4+ T cells at FDR<0.001 and AbsLFC>2, reaching significance only for cells from Malat1-/- mice (female)** |
| **Table S6** | **Differentially expressed genes between *WT* and *Malat1-/-* naïve CD4+ T cells from long-read sequencing (female)** |
| **Table S6** | **Differentially expressed genes between *WT* and *Malat1-/-* CD4+ T cells on day 2 of differentiation from long-read sequencing (female)** |
| **Table S8** | **Gene clusters identified by K-means clustering amonsgt up-regulated DEGs in day 2 *Malat1-/-* cells (related to Fig. S4C)** |
| **Table S9** | **Genes demonstrating DTU between WT and Malat1-/- cells on day 2 of CD4+ T cell differentiation** |

### TABLE S1

| Transcript | Forward Primer | Reverse Primer |
|---|---|---|
| *Malat1* | TGCAGTGTGCCAATGTTTCG | GGCCAGCTGCAAACATTCAA |
| *Il2ra* | GCATAGACTGTGTTGGCTTCTGC | GCGTTGCTTAGGAAACTCCTGG |
| *Il2rg* | GGAGCAACAGAGATCGAAGCTG | CCACAGATTGGGTTATAGCGGC |
| *Il2* | CCAATTCGATGATGAGTCAGC | CTTATGTGTTGTAAGCAGGAGG |
| *Cd69* | CCCTTGGGCTGTGTTAATA | AACTTCTCGTACAAGCCTG |
| *U6* | CGCTTCGGCAGCACATATAG | TTCACGAATTTGGCTGCTAT |
| *IFNAR1* | TGTGCTTCCCACCACTCAAG | AGGCGCGTGCTTTACTTCTA |
| *IFNAR2* | TGGGTATCCAGATGAACCTTG | GCCCTCCAACCACTTATCTG |
| *Ifit1bl1* | GAGATGGACTGTGAGGAAGGCT | ATCCAGGCGATAGGCTACGACT |
| *Ifit1* | GCAGAGAGTCAAGGCAGGTT | TTGTGCATCCCCAATGGGTT |
| *Ifit3b* | GCTCAGGCTTACGTTGACAAGG | CTTTAGGCGTGTCCATCCTTCC |
| *Iigp1* | GAGAACCAAGGTGGACTCTGAC | GAGAGCAGGAAGATTGGTGGCT |
| *Irf1* | CCCACAGAAGAGCATAGCAC | AGCAGTTCTTTGGGAATAGG |
| *Irf7* | GCATTTCGGTCGTAGGGATCTGGATGAAGA | CGTACACCTTATGCGGATCAACTGGA |
| *Stat1* | TACGGAAAAGCAAGCGTAATCT | TGCACATGACTTGATCCTTCAC |
| *IFNa(Ifna1+Ifna13)* | TGCCCAGCAGATCAAGAAGG | TCAGGGGAAATTCCTGCACC |
| *Ifnb* | GTACAACAGCTACGCCTGGA | GAGTCCGCCTCTGATGCTTA |
| *Ifna4* | CAATGATTGAACCCACATCCCCA | CCTTGCTTTCCAATTCTCTTTTCA |
| *Gata3* | Mm_Gata3_1_SG QuantiTect Primer Assay | |
| *Tbet* | Mm_Tbx21_1_SG Quantitect Primer Assay | |
| *Maf* | Mm_Maf_1_SG Quantitect Primer Assay | |

**TABLE S2**

| | Gene_ID | Ensml_ID | logFC | logCPM | F | PValue | FDR |
|---|---|---|---|---|---|---|---|
| **Female Naïve** | | | | | | | |
| **N=21 DEGs** | Cndp2 | ENSMUSG00000024644 | 124003199 | 564740744 | 399499497 | 394E-08 | 000045412 |
| | Gm5898 | ENSMUSG00000081788 | 329186833 | 197932603 | 235343049 | 313E-07 | 000180685 |
| | Fads1 | ENSMUSG00000010663 | -11989521 | 47213686 | 191526689 | 697E-07 | 000200966 |
| | Malat1 | ENSMUSG00000092341 | 112610618 | 893490283 | 20177943 | 570E-07 | 000200966 |
| | Gm10036 | ENSMUSG00000058064 | 416204637 | 309183072 | 178029192 | 924E-07 | 000213202 |
| | Gm28438 | ENSMUSG00000101939 | -43033586 | 049057825 | 156005751 | 154E-06 | 000295265 |
| | Gpatch11 | ENSMUSG00000050668 | 074292583 | 530790796 | 132450294 | 287E-06 | 000413931 |
| | NA | ENSMUSG00000095041 | 059531606 | 730819217 | 134902432 | 268E-06 | 000413931 |
| | Ncmap | ENSMUSG00000043924 | 17109089 | 372461192 | 994619818 | 847E-06 | 001085085 |
| | Tshz3 | ENSMUSG00000021217 | -18363991 | 146976014 | 766632201 | 222E-05 | 002564037 |
| | Sytl1 | ENSMUSG00000028860 | 087671097 | 502024473 | 696440497 | 316E-05 | 00331083 |
| | Cep85 | ENSMUSG00000037443 | -10779001 | 588507612 | 621189066 | 478E-05 | 004497193 |
| | Gm11223 | ENSMUSG00000046341 | -42018685 | 110732326 | 598508092 | 546E-05 | 004497193 |
| | Plaat3 | ENSMUSG00000060675 | -05812931 | 547201703 | 610203246 | 509E-05 | 004497193 |
| | Camk2b | ENSMUSG00000057897 | -11949753 | 468187477 | 532848579 | 826E-05 | 006349555 |
| | Spry2 | ENSMUSG00000022114 | -07151442 | 350693821 | 513499555 | 941E-05 | 00678323 |
| | Ubxn11 | ENSMUSG00000012126 | 062758058 | 592566704 | 483225682 | 000011646 | 007902086 |
| | Gm10110 | ENSMUSG00000062093 | 150163921 | 083517913 | 466130837 | 000013205 | 008462357 |
| | Wdtc1 | ENSMUSG00000037622 | 040784966 | 618604563 | 429665604 | 000017507 | 010628688 |
| | Ighv5-16 | ENSMUSG00000094194 | -40105197 | 048492404 | 404070343 | 000021609 | 012463154 |
| | Carns1 | ENSMUSG00000075289 | 076361762 | 58838525 | 376137618 | 000027557 | 015136553 |
| | Nipal3 | ENSMUSG00000028803 | -03841585 | 637466436 | 370615681 | 000028966 | 015187236 |

| | Gene_ID | Ensmbl | logFCWTvKO_ | logCPMWTvK | FWTvKO_fem | PValueWTvK | FDRWTvKO_female |
|---|---|---|---|---|---|---|---|
| **Female Th2** | | | | | | | |
| **N=8 DEGs** | Malat1 | ENSMUSG00000092341 | 112967177 | 845184692 | 210082293 | 455E-07 | 000530819 |
| | Gm28438 | ENSMUSG00000101939 | -70364878 | 110104827 | 143218534 | 201E-06 | 001172574 |
| | Cndp2 | ENSMUSG00000024644 | 10565631 | 584273855 | 935472611 | 101E-05 | 003946454 |
| | Prnp | ENSMUSG00000079037 | 166602665 | 539207726 | 764989416 | 215E-05 | 006259273 |
| | Espn | ENSMUSG00000028943 | 06531511 | 610687836 | 656868943 | 375E-05 | 008756032 |
| | Gm11223 | ENSMUSG00000046341 | -36081363 | 29841745 | 485907783 | 000011047 | 014326908 |
| | Gpatch11 | ENSMUSG00000050668 | 090052631 | 529565472 | 489153736 | 000010791 | 014326908 |
| | Tshz3 | ENSMUSG00000021217 | -15193265 | 120279004 | 493252626 | 000010478 | 014326908 |
| | Zdhhc24 | ENSMUSG00000006463 | -09276665 | 47450222 | 505795475 | 959E-05 | 014326908 |

| | Gene_ID | Ensml_ID | logFC | logCPM | F | PValue | FDR |
|---|---|---|---|---|---|---|---|
| **Male Naïve** | | | | | | | |
| **N=2 DEGs** | Malat1 | ENSMUSG00000092341 | 113506263 | 856192378 | 200155787 | 225E-06 | 002684758 |
| | Cndp2 | ENSMUSG00000024644 | 120957576 | 572925269 | 128425886 | 991E-06 | 00415496 |
| | Gm28438 | ENSMUSG00000101939 | -50230092 | 117710548 | 126425749 | 104E-05 | 00415496 |

| | Gene_ID | Ensml_ID | logFC | logCPM | F | PValue | FDR |
|---|---|---|---|---|---|---|---|
| **Male Th2** | | | | | | | |
| **N=28 DEGs** | Malat1 | ENSMUSG00000092341 | 126994369 | 854811081 | 224687744 | 909E-12 | 109E-07 |
| | Gpatch11 | ENSMUSG00000050668 | 094489291 | 533515253 | 519525564 | 492E-09 | 295E-05 |
| | Prnp | ENSMUSG00000079037 | 145587965 | 546235325 | 268687918 | 805E-08 | 000032191 |
| | Gm11223 | ENSMUSG00000046341 | -32893304 | 268127817 | 20601388 | 245E-07 | 0000588 |
| | Gm28438 | ENSMUSG00000101939 | -57600318 | 096501264 | 216291297 | 200E-07 | 0000588 |
| | Camk2b | ENSMUSG00000057897 | -13488713 | 427396807 | 189832652 | 345E-07 | 000068899 |
| | 4930453N24 | ENSMUSG00000059920 | -06078388 | 630154155 | 170910376 | 533E-07 | 000091319 |
| | Fads1 | ENSMUSG00000010663 | -06629958 | 564961436 | 122383535 | 210E-06 | 000315118 |
| | Mmrn1 | ENSMUSG00000054641 | -11416206 | 124107132 | 262269879 | 585E-06 | 000779834 |
| | Cndp2 | ENSMUSG00000024644 | 105934316 | 609347641 | 92255987 | 657E-06 | 000788286 |
| | Zdhhc24 | ENSMUSG00000006463 | -07806963 | 449045174 | 898474074 | 731E-06 | 000796398 |
| | Mid1-ps1 | ENSMUSG00000095134 | -38671242 | 121561799 | 798562161 | 117E-05 | 001165224 |
| | Eps8l1 | ENSMUSG00000006154 | -12467447 | 431458894 | 772571982 | 133E-05 | 001225393 |
| | Gm43305 | ENSMUSG00000105703 | -14700852 | 702270346 | 721185138 | 174E-05 | 001490526 |
| | Carns1 | ENSMUSG00000075289 | 066557635 | 458221572 | 593386121 | 370E-05 | 002959927 |
| | Gm10505 | ENSMUSG00000073430 | -090448 | 347802242 | 569830696 | 432E-05 | 003239473 |
| | Nthl1 | ENSMUSG00000041429 | -08066626 | 142896856 | 536656441 | 543E-05 | 003829072 |
| | Tkfc | ENSMUSG00000034371 | 029457902 | 480232362 | 471926844 | 879E-05 | 005550095 |
| | Ap5b1 | ENSMUSG00000049562 | -05200927 | 42226486 | 478237374 | 837E-05 | 005550095 |
| | Erdr1 | ENSMUSG00000095562 | 219312559 | 250142018 | 441214127 | 000011282 | 006764786 |
| | Lrfn4 | ENSMUSG00000045045 | 114470887 | 120744388 | 409433233 | 000014838 | 008434841 |
| | Vps37c | ENSMUSG00000048832 | 034148126 | 446357203 | 404738484 | 000015474 | 008434841 |

| | | | | | | |
|---|---|---|---|---|---|---|
| Mmp13 | ENSMUSG00000050578 | -09462041 | 247241411 | 394716737 | 000016949 | 008836877 |
| Patl1 | ENSMUSG00000046139 | -01786281 | 629048601 | 351176124 | 000025778 | 012880557 |
| Slc25a45 | ENSMUSG00000024818 | -02644545 | 652863904 | 341743873 | 000028389 | 013596399 |
| Rad9b | ENSMUSG00000038569 | 076147423 | 322955105 | 338119272 | 000029479 | 013596399 |
| Shc1 | ENSMUSG00000042626 | -0179956 | 686470335 | 333539267 | 00003093 | 013737363 |
| Eef1d | ENSMUSG00000055762 | 018636907 | 727628474 | 307359656 | 000041148 | 017576145 |
| Tmem181b-p | ENSMUSG00000096780 | -71798582 | 332369029 | 304494918 | 000042504 | 017576145 |

**TABLE S3**

| Gene | ENSMBL | logFC_WT_F | logCPM_WT_F | F_WT_F | PValue_WT_F | FDR_WT_F | logFC_KO_F | logCPM_KO_F | F_KO_F | PValue_KO_F | FDR_KO_F | delta_LFC(KO-WT) | higher in KOs |
|---|---|---|---|---|---|---|---|---|---|---|---|---|---|
| Apol9b | ENSMUSG00000068246 | -1201369698 | 1066392531 | 3612473283 | 502E-07 | 114E-05 | -7782926674 | 1249196006 | 1598954204 | 684E-07 | 353E-05 | 4230770306 | 1 |
| Hspa1a | ENSMUSG00000091971 | -1200608908 | 6006947072 | 7838560828 | 129E-05 | 960E-05 | -1113482567 | 6822935704 | 1671302747 | 569E-07 | 338E-05 | 087126341 | 1 |
| Hspa1b | ENSMUSG00000090877 | -1126391326 | 8152099177 | 1285583491 | 179E-06 | 259E-05 | -1082527641 | 9038986971 | 2037387493 | 249E-07 | 223E-05 | 043863685 | 1 |
| Car1 | ENSMUSG00000027556 | -9929147567 | 2997533637 | 3684860475 | 225E-08 | 183E-06 | -1137869016 | 4796798499 | 1850175751 | 373E-07 | 273E-05 | -1449542593 | 0 |
| Slc16a5 | ENSMUSG00000045775 | -7663128082 | 4832452358 | 266709047 | 477E-12 | 278E-08 | -6354010755 | 4564448515 | 6878456903 | 142E-09 | 229E-06 | 1309117327 | 1 |
| Gm11695 | ENSMUSG00000085651 | -759674646 | 141117526 | 1823069437 | 426E-07 | 103E-05 | -5035938441 | 12653286 | 2295412447 | 151E-07 | 168E-05 | 2560808019 | 1 |
| Ncr1 | ENSMUSG00000062524 | -7333365709 | 1974117855 | 14339847 | 114E-06 | 190E-05 | -5845420817 | 3083921235 | 1701078941 | 529E-07 | 334E-05 | 1487944892 | 1 |
| Oasl2 | ENSMUSG00000029561 | -7269764492 | 3507699108 | 2306925002 | 160E-07 | 564E-06 | -5870392258 | 3828095291 | 354324526 | 242E-08 | 683E-06 | 1399372234 | 1 |
| Oas2 | ENSMUSG00000032690 | -7126755992 | 5372718626 | 8003396369 | 832E-10 | 347E-07 | -5999864247 | 5251300305 | 3082776473 | 436E-08 | 867E-06 | 1126891745 | 1 |
| S1pr5 | ENSMUSG00000045087 | -6882209642 | 1178767859 | 9113634602 | 713E-06 | 636E-05 | -7316303186 | 2055729837 | 2643851484 | 835E-08 | 130E-05 | -0434093544 | 0 |
| Dtx1 | ENSMUSG00000029603 | -6863481384 | 7792754765 | 1708227295 | 324E-11 | 126E-07 | -5460047036 | 7588275488 | 1761639987 | 457E-07 | 315E-05 | 1403434348 | 1 |
| Fosb | ENSMUSG00000003545 | -6223761041 | 7526143189 | 5504197665 | 505E-05 | 0000270438 | -6951135712 | 7223633193 | 3029103782 | 470E-08 | 890E-06 | -0727374671 | 0 |
| Sardh | ENSMUSG00000009614 | -6162530565 | 0442493179 | 1153493943 | 277E-06 | 339E-05 | -7793832288 | 0120766051 | 1351077773 | 137E-06 | 513E-05 | -1631301723 | 0 |
| Trpm1 | ENSMUSG00000030523 | -6044514704 | 1331691235 | 1716203602 | 547E-07 | 119E-05 | -4817375126 | 1595709101 | 3956975367 | 151E-08 | 595E-06 | 1227139578 | 1 |
| Fos | ENSMUSG00000021250 | -5924551293 | 8726610491 | 5044687067 | 701E-05 | 0000349401 | -7041207126 | 8547466236 | 1898681088 | 335E-07 | 252E-05 | -1116655833 | 0 |
| Ces2g | ENSMUSG00000031877 | -5918107247 | 1164915869 | 7155565451 | 185E-05 | 0000124807 | -6176107102 | 3251189089 | 5638934318 | 443E-05 | 0000386222 | -0257999855 | 0 |
| Itga9 | ENSMUSG00000039115 | -5823378343 | 1185498933 | 3057760887 | 493E-08 | 282E-06 | -3990356826 | 1417195615 | 1144327955 | 270E-06 | 720E-05 | 1833021517 | 1 |
| Ifit3b | ENSMUSG00000062488 | -5735773208 | 1412288082 | 9528561101 | 597E-06 | 560E-05 | -4348401883 | 1553427131 | 1196486738 | 225E-06 | 652E-05 | 1387371325 | 1 |
| Hspa12a | ENSMUSG00000025092 | -5710446876 | 180462443 | 9083851334 | 722E-06 | 640E-05 | -4133587989 | 168445482 | 3450885252 | 270E-08 | 728E-06 | 1576858887 | 1 |
| Igha | ENSMUSG00000095079 | -5703692907 | 395934915 | 1347477601 | 148E-06 | 229E-05 | -7793618522 | 7446763626 | 4157376745 | 0000138571 | 0000817875 | -2089925615 | 0 |
| Rtp4 | ENSMUSG00000033355 | -555659634 | 4059468508 | 1756434868 | 497E-07 | 114E-05 | -4320865432 | 4153170814 | 9655511458 | 537E-06 | 0000104804 | 1235730908 | 1 |
| Atp1b1 | ENSMUSG00000026576 | -5547849828 | 5953803631 | 3742216561 | 210E-08 | 178E-06 | -5093241266 | 603403456 | 5174340981 | 480E-09 | 364E-06 | 0454608562 | 1 |
| Xkrx | ENSMUSG00000031258 | -5497390541 | 3700558553 | 5741678293 | 343E-09 | 714E-07 | -474001119 | 370788998 | 3872935582 | 165E-08 | 595E-06 | 0757379351 | 1 |
| Jun | ENSMUSG00000052684 | -5478308223 | 8927190233 | 203709137 | 269E-07 | 777E-06 | -5201009808 | 863636039 | 1432606826 | 108E-06 | 447E-05 | 0277298415 | 1 |
| Adam6b | ENSMUSG00000051804 | -544472888 | 0439470615 | 4621411152 | 971E-05 | 0000445207 | -4781597033 | 0515960521 | 9858523075 | 493E-06 | 0000101529 | 0663131847 | 1 |
| Ermap | ENSMUSG00000028644 | -5433759148 | 1897188532 | 1327449545 | 157E-06 | 238E-05 | -736592162 | 4060864122 | 6499286843 | 256E-05 | 000027262 | -1932162472 | 0 |
| Gm15915 | ENSMUSG00000085723 | -5359156914 | 1249217417 | 1205065999 | 232E-06 | 304E-05 | -4500706964 | 2389741767 | 1430709795 | 108E-06 | 447E-05 | 085844995 | 1 |
| Oasl1 | ENSMUSG00000041827 | -5320987671 | 298746674 | 2680642027 | 856E-08 | 382E-06 | -4702805174 | 2967870864 | 321188482 | 366E-08 | 823E-06 | 0618182497 | 1 |
| Nr4a1 | ENSMUSG00000023034 | -5149196264 | 7942844812 | 1181029686 | 252E-06 | 318E-05 | -4307323788 | 7552753057 | 7746370412 | 129E-05 | 0000181355 | 0841872476 | 1 |
| Ifit3 | ENSMUSG00000074896 | -5099508628 | 3754712216 | 4539512629 | 929E-09 | 113E-06 | -4391555868 | 3834322604 | 1120069132 | 295E-06 | 755E-05 | 070795276 | 1 |
| Dock4 | ENSMUSG00000035954 | -5087682337 | 2169099956 | 3528142026 | 270E-08 | 200E-06 | -3988067789 | 2166068981 | 1071316835 | 353E-06 | 831E-05 | 1099614548 | 1 |
| Klf4 | ENSMUSG00000003032 | -5067687793 | 5080597815 | 1598214829 | 734E-07 | 142E-05 | -4918429777 | 4798644403 | 3639660128 | 216E-08 | 673E-06 | 0149258016 | 1 |
| Aqp1 | ENSMUSG00000004655 | -5038288061 | 2033086504 | 1049995479 | 405E-06 | 436E-05 | -714109557 | 4467805231 | 7301354983 | 163E-05 | 0000209926 | -2102807509 | 0 |
| Chdh | ENSMUSG00000015970 | -4999764105 | 19298933 | 9034251482 | 738E-06 | 648E-05 | -3085544482 | 1911173965 | 1079207342 | 342E-06 | 818E-05 | 1914219623 | 1 |
| Mx2 | ENSMUSG00000023341 | -4857131847 | 2247265635 | 9235634391 | 676E-06 | 612E-05 | -4157246938 | 2094853358 | 2394415912 | 127E-07 | 160E-05 | 0699884909 | 1 |
| 1810006J02Rik | ENSMUSG00000100182 | -4832368785 | 1652507033 | 6552354663 | 260E-05 | 0000160627 | -3089264362 | 1610148152 | 1440451876 | 105E-06 | 444E-05 | 1743104423 | 1 |
| Ifi44 | ENSMUSG00000028037 | -4742156898 | 1221146342 | 1168889147 | 263E-06 | 326E-05 | -4802756925 | 0851980145 | 5378100656 | 530E-05 | 0000432067 | -0060600027 | 0 |
| Rgmb | ENSMUSG00000048027 | -473762529 | 3484546015 | 8316807897 | 706E-10 | 347E-07 | -3989427167 | 3553539717 | 3791703252 | 181E-08 | 616E-06 | 0748198123 | 1 |
| Grm6 | ENSMUSG00000000617 | -4714563162 | 1684197149 | 8563744786 | 912E-06 | 744E-05 | -3015057795 | 1541463578 | 7384709821 | 156E-05 | 0000203368 | 1699505367 | 1 |
| Cd163 | ENSMUSG00000008845 | -4693310959 | 0829226374 | 1083600308 | 357E-06 | 401E-05 | -6095937563 | 1924813541 | 1530125099 | 820E-07 | 395E-05 | -1402626604 | 0 |
| Arhgap20 | ENSMUSG00000053199 | -455230002 | 1241479129 | 5203412028 | 624E-05 | 0000317475 | -4451122861 | 1444483669 | 1199310591 | 223E-06 | 652E-05 | 0101177159 | 1 |
| Oas1a | ENSMUSG00000052776 | -4538392303 | 2855458423 | 2880419847 | 633E-08 | 326E-06 | -2803067088 | 2736174659 | 6053135022 | 338E-05 | 0000324683 | 1735325215 | 1 |
| Nsg2 | ENSMUSG00000020297 | -4495848447 | 7415934719 | 562354989 | 374E-09 | 727E-07 | -3818094484 | 7299163481 | 994548831 | 476E-06 | 100E-04 | 0677753963 | 1 |
| Ntn1 | ENSMUSG00000020902 | -4458002867 | 1025449527 | 2507852564 | 113E-07 | 450E-06 | -4192427476 | 1132828129 | 4394361643 | 966E-09 | 509E-06 | 0265575391 | 1 |
| Dapl1 | ENSMUSG00000026989 | -4456593647 | 490186939 | 2672575025 | 867E-08 | 382E-06 | -3202522327 | 4979821845 | 5836852872 | 287E-09 | 242E-06 | 125407132 | 1 |
| Ppp1r15a | ENSMUSG00000040435 | -4437040138 | 8744584483 | 3169886205 | 423E-08 | 257E-06 | -4259027521 | 8445340447 | 818399276 | 104E-05 | 0000159607 | 0178012617 | 1 |
| Ccr9 | ENSMUSG00000029530 | -4266968489 | 2762968487 | 1209163629 | 229E-06 | 302E-05 | -4982649207 | 3000558131 | 3155561862 | 395E-08 | 856E-06 | -0715680718 | 0 |
| Klra14-ps | ENSMUSG00000072721 | -4258849947 | 0351613388 | 382574074 | 0000193394 | 0000757288 | -4633931961 | 137619274 | 107271044 | 351E-06 | 831E-05 | -0375082014 | 0 |
| Gm18752 | ENSMUSG00000103105 | -4217994953 | 1291331256 | 6173573029 | 326E-05 | 0000192067 | -3640754 | 1354401663 | 7847889372 | 123E-05 | 0000175521 | 0577240953 | 1 |
| Gm11707 | ENSMUSG00000086481 | -4215045755 | 2621522488 | 2274191679 | 170E-07 | 586E-06 | -4764523321 | 252781387 | 3897252589 | 161E-08 | 595E-06 | -0549477566 | 0 |
| Adrb2 | ENSMUSG00000045730 | -4212396139 | 4994904594 | 3112405181 | 457E-08 | 267E-06 | -3796564487 | 4990606841 | 2485571885 | 108E-07 | 152E-05 | 0415831652 | 1 |
| Klra4 | ENSMUSG00000079852 | -4192953453 | 1371002293 | 1341707791 | 150E-06 | 232E-05 | -4592625903 | 2321768177 | 1162289858 | 253E-06 | 705E-05 | -039967245 | 0 |
| Tox | ENSMUSG00000041272 | -4180659096 | 5036327939 | 4684482138 | 813E-09 | 108E-06 | -3658461207 | 4968094442 | 6738659516 | 155E-09 | 229E-06 | 0522197889 | 1 |
| Arc | ENSMUSG00000022602 | -4101095087 | 3987620667 | 1180884126 | 252E-06 | 318E-05 | -3114641012 | 3500420739 | 6815443155 | 213E-05 | 0000246855 | 0986454075 | 1 |
| Nt5e | ENSMUSG00000032420 | -4048894914 | 4405314009 | 1176741127 | 256E-06 | 321E-05 | -3151351988 | 4576657357 | 1697035164 | 534E-07 | 334E-05 | 0897542926 | 1 |
| Rasal1 | ENSMUSG00000029602 | -404328448 | 5012505953 | 8205193587 | 748E-10 | 347E-07 | -3348206483 | 4936535009 | 6886406165 | 141E-09 | 229E-06 | 0695077997 | 1 |
| Lbp | ENSMUSG00000016024 | -4006854087 | 1275471123 | 8207267262 | 108E-05 | 838E-05 | -2739388444 | 138020252 | 1007510556 | 452E-06 | 962E-05 | 1267465643 | 1 |
| Adamts14 | ENSMUSG00000059901 | -4005452472 | 3919956358 | 306263948 | 489E-08 | 281E-06 | -3333837872 | 4486835498 | 3964616529 | 150E-08 | 595E-06 | 06716146 | 1 |
| Rasl11b | ENSMUSG00000049907 | -39976151 | 2742703058 | 2399455626 | 136E-07 | 509E-06 | -3529337063 | 2621075803 | 8969158211 | 721E-06 | 0000127106 | 0468278037 | 1 |
| Apol8 | ENSMUSG00000056656 | -3977014424 | 131011069 | 3910200864 | 0000178784 | 0000716036 | -3864166491 | 0941047009 | 4421147127 | 0000110511 | 0000705832 | 0112847933 | 1 |
| Gm4956 | ENSMUSG00000025936 | -3953434218 | 1049239261 | 1015741379 | 462E-06 | 475E-05 | -3018571122 | 1407353551 | 1520333913 | 842E-07 | 399E-05 | 0934863096 | 1 |
| Dusp1 | ENSMUSG00000024190 | -390426271 | 7543820687 | 834542756 | 101E-05 | 804E-05 | -3986731303 | 7414267888 | 1390050278 | 122E-06 | 488E-05 | -0082468593 | 0 |
| Rgs2 | ENSMUSG00000026360 | -3868581531 | 6511199766 | 1826838947 | 423E-07 | 103E-05 | -4403066005 | 6438492884 | 2110942409 | 215E-07 | 211E-05 | -0534484474 | 0 |
| Bambi | ENSMUSG00000024232 | -3866065354 | 2798313036 | 6574020333 | 256E-05 | 000015927 | -4757673741 | 257754135 | 6518352497 | 254E-05 | 000027194 | -0891608387 | 0 |
| Vangl2 | ENSMUSG00000026556 | -3851011194 | 2327572063 | 5210463355 | 621E-05 | 0000316278 | -344785399 | 2358657126 | 6382080902 | 275E-05 | 0000285531 | 0403157204 | 1 |
| Ampd1 | ENSMUSG00000070385 | -3827164632 | 3746892704 | 4381452924 | 108E-08 | 121E-06 | -3913401736 | 3934023431 | 1228396131 | 202E-06 | 641E-05 | -0086237104 | 0 |
| Ifit1 | ENSMUSG00000034459 | -3826004574 | 4114771973 | 4797733488 | 735E-09 | 105E-06 | -3219463024 | 4160953562 | 1152983439 | 262E-06 | 706E-05 | 060654155 | 1 |
| Klra8 | ENSMUSG00000089727 | -3772856328 | 168509502 | 112755291 | 304E-06 | 360E-05 | -4510659953 | 2834164107 | 6144564398 | 319E-05 | 0000313317 | -0737803625 | 0 |
| Dnajc6 | ENSMUSG00000028528 | -3759292504 | 1879040669 | 1094410474 | 343E-06 | 390E-05 | -3662982716 | 2266729127 | 3058717641 | 451E-08 | 870E-06 | 0096309788 | 1 |
| Ifi213 | ENSMUSG00000073491 | -3744322161 | 7378800761 | 9895798227 | 336E-10 | 268E-07 | -3016580613 | 7432740593 | 5791838392 | 400E-05 | 0000361671 | 0727741548 | 1 |
| Egr1 | ENSMUSG00000038418 | -3695240211 | 7346032877 | 6312793538 | 300E-05 | 0000179947 | -3100884402 | 7004661799 | 1726494866 | 497E-07 | 329E-05 | 0594355809 | 1 |
| AW011738 | ENSMUSG00000078349 | -3689012712 | 3034396948 | 4632631621 | 852E-09 | 108E-06 | -2921639159 | 3063640434 | 725963618 | 167E-05 | 0000213368 | 0767373553 | 1 |
| Il17rb | ENSMUSG00000015966 | -3641033152 | 2113276862 | 1706116914 | 561E-07 | 120E-05 | -2194669801 | 2008701581 | 5346000241 | 543E-05 | 0000437002 | 1446363351 | 1 |
| Ifi208 | ENSMUSG00000066677 | -3613641675 | 6984343191 | 6181321407 | 250E-09 | 603E-07 | -3061458535 | 7076626446 | 6530805751 | 252E-05 | 0000271113 | 055218314 | 1 |
| Atp1b2 | ENSMUSG00000041329 | -3609127627 | 0723677991 | 6602176502 | 252E-05 | 0000157419 | -5089691492 | 2305069483 | 5830070613 | 390E-05 | 0000357081 | -1480563865 | 0 |
| Gm42778 | ENSMUSG00000105112 | -360320704 | 1102526147 | 8055454071 | 116E-05 | 887E-05 | -3615611249 | 1094571329 | 1025218832 | 421E-06 | 925E-05 | -0012404209 | 0 |
| Isg15 | ENSMUSG00000035692 | -3594136657 | 307282065 | 1483982617 | 995E-07 | 172E-05 | -2656870798 | 2974337407 | 1144471266 | 270E-06 | 720E-05 | 0937265859 | 1 |
| Mcoln3 | ENSMUSG00000036853 | -3588906208 | 2242058246 | 1451806984 | 109E-06 | 184E-05 | -3354872639 | 2031313726 | 1382810935 | 125E-06 | 493E-05 | 0234033569 | 1 |
| Slc22a3 | ENSMUSG00000023828 | -356841288 | 1270277716 | 6780474145 | 227E-05 | 0000146531 | -4749489611 | 2412313788 | 1986569087 | 277E-07 | 232E-05 | -1181076731 | 0 |
| Hes1 | ENSMUSG00000022528 | -3487750614 | 059970048 | 3910044851 | 000017881 | 0000716036 | -2815620509 | 1380723355 | 747545416 | 148E-05 | 0000196801 | 0672130105 | 1 |
| Nr4a3 | ENSMUSG00000028341 | -3455686946 | 4003579054 | 1848121535 | 403E-07 | 996E-06 | -2558450159 | 4152007537 | 1212381771 | 213E-06 | 651E-05 | 0897236787 | 1 |
| Fam240a | ENSMUSG00000096393 | -3437140204 | 0970284646 | 1211012439 | 228E-06 | 301E-05 | -2781482189 | 0622283176 | 4949870534 | 726E-05 | 0000529789 | 0655658015 | 1 |
| Fam186b | ENSMUSG00000078907 | -3387047311 | 1078442772 | 8992706339 | 752E-06 | 657E-05 | -2888970825 | 0998724199 | 1059330743 | 369E-06 | 853E-05 | 0498076486 | 1 |
| Klrb1c | ENSMUSG00000030325 | -3363331311 | 1957739852 | 4451559157 | 0000111514 | 0000495408 | -3588813354 | 2868521497 | 6281322811 | 293E-05 | 000029606 | -0225482043 | 0 |
| Prag1 | ENSMUSG00000050271 | -3352652694 | 5334758949 | 4528256249 | 939E-09 | 113E-06 | -3126269915 | 5159302663 | 3655092335 | 212E-08 | 673E-06 | 0226382779 | 1 |
| Cnga1 | ENSMUSG00000067220 | -3350727457 | 3148648021 | 1685256213 | 590E-07 | 124E-05 | -3010623844 | 3016151728 | 6293722092 | 208E-09 | 229E-06 | 0340103613 | 1 |
| Esrp2 | ENSMUSG00000084128 | -3283176205 | 0688867285 | 4413375484 | 0000115099 | 0000507972 | -2455791604 | 0524134984 | 7925821421 | 118E-05 | 0000171612 | 0827384601 | 1 |
| Btbd11 | ENSMUSG00000020042 | -3279219773 | 5141898876 | 7148641808 | 135E-09 | 449E-07 | -2484371921 | 5057827945 | 7479094362 | 148E-05 | 0000196801 | 0794847852 | 1 |
| Ggt5 | ENSMUSG00000006344 | -3266731364 | 5274376087 | 8730833518 | 574E-10 | 347E-07 | -3367270425 | 5167843907 | 1619899805 | 648E-07 | 352E-05 | -0100539061 | 0 |
| Cx3cr1 | ENSMUSG00000052336 | -3265193649 | 1610420283 | 505412787 | 696E-05 | 0000347512 | -3727010798 | 2778464802 | 500628296 | 695E-05 | 0000515779 | -0461817149 | 0 |
| Ttc28 | ENSMUSG00000033209 | -3263713274 | 5829881752 | 5192783775 | 525E-09 | 862E-07 | -2967375692 | 5795007071 | 3295840347 | 328E-08 | 772E-06 | 0296337582 | 1 |
| Nfkbiz | ENSMUSG00000035356 | -3230764704 | 6997829521 | 1829642454 | 420E-07 | 103E-05 | -2672256935 | 6864660784 | 8036532062 | 112E-05 | 0000166671 | 0558507769 | 1 |
| Gm19951 | ENSMUSG00000113136 | -3163734392 | 1125590928 | 8472594136 | 952E-06 | 768E-05 | -3230200893 | 0711606075 | 56575459 | 437E-05 | 000038277 | -0066466501 | 0 |
| Txk | ENSMUSG00000054892 | -315683644 | 6712279347 | 1008655643 | 309E-10 | 268E-07 | -2581718935 | 6753324484 | 1924328044 | 317E-07 | 249E-05 | 0575117505 | 1 |
| 2510009E07Rik | ENSMUSG00000043391 | -3151357787 | 183563177 | 832254618 | 102E-05 | 807E-05 | -2729915229 | 1865411912 | 9889831813 | 487E-06 | 0000101284 | 0421442558 | 1 |
| Lrrc23 | ENSMUSG00000030125 | -3135658111 | 1021066641 | 1420127894 | 119E-06 | 195E-05 | -437992963 | 0897496757 | 2882073354 | 580E-08 | 102E-05 | -1244271519 | 0 |
| Tulp2 | ENSMUSG00000023467 | -3129926224 | 1686996999 | 9481464906 | 609E-06 | 568E-05 | -2962332949 | 1719236654 | 3883941657 | 000017753 | 0000971041 | 0167593275 | 1 |
| Sox4 | ENSMUSG00000076431 | -3114783697 | 4389847937 | 3213994965 | 400E-08 | 249E-06 | -3324728837 | 4139415805 | 5776805735 | 300E-09 | 242E-06 | -020994514 | 0 |
| Trpv6 | ENSMUSG00000029868 | -311317107 | 161174554 | 9238042724 | 675E-06 | 612E-05 | -2459963723 | 1500412429 | 1616610681 | 653E-07 | 352E-05 | 0653207347 | 1 |
| Esr1 | ENSMUSG00000019768 | -3111201912 | 147471322 | 1318998499 | 161E-06 | 242E-05 | -2723281203 | 1536184653 | 1376531783 | 127E-06 | 496E-05 | 0387920709 | 1 |
| Gm42725 | ENSMUSG00000105601 | -3109385396 | 1170044735 | 644213128 | 277E-05 | 0000168969 | -2432021215 | 0943534912 | 4363561902 | 0000115991 | 0000730075 | 0677364181 | 1 |
| Arhgap29 | ENSMUSG00000039831 | -3093972629 | 4324875067 | 4024394545 | 155E-08 | 150E-06 | -2935208113 | 4202182524 | 1943708911 | 304E-07 | 246E-05 | 0158764516 | 1 |
| Lrig1 | ENSMUSG00000030029 | -3080171557 | 5709885102 | 7005250933 | 147E-09 | 451E-07 | -3261535556 | 5581549976 | 2381621522 | 130E-07 | 160E-05 | -0181363999 | 0 |
| Il1rl2 | ENSMUSG00000070942 | -3074177483 | 4024758737 | 4582268045 | 893E-09 | 111E-06 | -2505109015 | 4002593672 | 1205308169 | 219E-06 | 652E-05 | 0569068468 | 1 |
| Arhgap5 | ENSMUSG00000035133 | -3058653724 | 4195286556 | 5198537617 | 523E-09 | 862E-07 | -3013012615 | 4147392907 | 4291494513 | 107E-08 | 540E-06 | 0045641109 | 1 |
| Cd69 | ENSMUSG00000030156 | -3052685514 | 700226268 | 9015974744 | 744E-06 | 653E-05 | -2953359735 | 675844881 | 6000172831 | 349E-05 | 0000330345 | 0099325779 | 1 |
| Tnf | ENSMUSG00000024401 | -304138621 | 4171163186 | 2208110133 | 192E-07 | 627E-06 | -2259691489 | 3935064935 | 5521828129 | 480E-05 | 0000404997 | 0781694721 | 1 |
| Zhx2 | ENSMUSG00000071757 | -2980299505 | 5588703961 | 1681234755 | 596E-07 | 125E-05 | -2152511096 | 5525308941 | 3956401854 | 0000166017 | 0000923681 | 0827788409 | 1 |

| | | | | | | | | | | | | | |
|---|---|---|---|---|---|---|---|---|---|---|---|---|---|
| Plk3 | ENSMUSG00000028680 | -2925184969 | 5594195105 | 1155689981 | 275E-06 | 337E-05 | -2398697288 | 5265661143 | 798705265 | 114E-05 | 0000169129 | 0526487681 | 1 |
| E430024P14Rik | ENSMUSG00000090069 | -2924882296 | 2299036433 | 7903395886 | 125E-05 | 938E-05 | -2800666354 | 2096651413 | 6055018826 | 337E-05 | 0000324632 | 0124215942 | 1 |
| Cyp2d22 | ENSMUSG00000061740 | -2896295502 | 2993523992 | 9893540009 | 514E-06 | 510E-05 | -2780182343 | 3068548601 | 9053151277 | 695E-06 | 0000124081 | 0116113159 | 1 |
| Gm37780 | ENSMUSG00000102691 | -2863377637 | 1475914097 | 586362748 | 397E-05 | 000022445 | -3110718058 | 1183673251 | 6743024716 | 222E-05 | 0000253377 | -0247340421 | 0 |
| G0s2 | ENSMUSG00000009633 | -2841898802 | 0721484931 | 366650246 | 000022517 | 0000844586 | -2824063842 | 1025543123 | 616119323 | 315E-05 | 0000311439 | 001783496 | 1 |
| Ramp1 | ENSMUSG00000034353 | -2818480343 | 4869264409 | 1414380842 | 121E-06 | 197E-05 | -3905987932 | 4654913897 | 7630724784 | 137E-05 | 0000188286 | -1087507589 | 0 |
| Dnajb9 | ENSMUSG00000014905 | -2809686661 | 6228554165 | 1863522479 | 389E-07 | 981E-06 | -2601182672 | 6021123041 | 6058799138 | 336E-05 | 0000324546 | 0208503989 | 1 |
| Sgk3 | ENSMUSG00000025915 | -2754596331 | 4903963617 | 6885926209 | 158E-09 | 473E-07 | -2657127262 | 5013052409 | 1435196762 | 107E-06 | 447E-05 | 0097469069 | 1 |
| Ccl3 | ENSMUSG00000000982 | -2743535997 | 2442241893 | 719455489 | 181E-05 | 0000122885 | -2717355296 | 3346484033 | 1643080546 | 611E-07 | 343E-05 | 0026180701 | 1 |
| Zdhhc14 | ENSMUSG00000034265 | -2711058025 | 2132101369 | 1211929555 | 227E-06 | 301E-05 | -3018653205 | 2119462465 | 1336005321 | 143E-06 | 529E-05 | -030759518 | 0 |
| Fgfrl1 | ENSMUSG00000008090 | -2704707304 | 0849339127 | 7303989278 | 170E-05 | 0000117858 | -2486536885 | 1129388491 | 1099673443 | 317E-06 | 783E-05 | 0218170419 | 1 |
| Rapgef4 | ENSMUSG00000049044 | -2699165335 | 5585711586 | 3184249897 | 415E-08 | 255E-06 | -2363871921 | 5489166391 | 5306451497 | 558E-05 | 0000445923 | 0335293414 | 1 |
| Junb | ENSMUSG00000052837 | -2693882431 | 1048541628 | 1871920537 | 382E-07 | 974E-06 | -2494687899 | 10181277 | 5603297084 | 454E-05 | 0000391179 | 0199194532 | 1 |
| App | ENSMUSG00000022892 | -2686626334 | 4260209803 | 5280983241 | 591E-05 | 0000303741 | -234777947 | 4460696908 | 4771807998 | 832E-05 | 0000580692 | 0338846864 | 1 |
| Amz1 | ENSMUSG00000050022 | -268483362 | 260733654 | 9010465904 | 746E-06 | 654E-05 | -2880757359 | 240523469 | 1215525266 | 211E-06 | 651E-05 | -0195923739 | 0 |
| Gadd45a | ENSMUSG00000036390 | -2679438956 | 3637655736 | 1183098849 | 250E-06 | 317E-05 | -2746197372 | 3487035574 | 5136112388 | 631E-05 | 0000484056 | -0066758416 | 0 |
| Rsad2 | ENSMUSG00000020641 | -2653371761 | 2673311442 | 6743749506 | 232E-05 | 0000148526 | -2126574281 | 2831908031 | 9382046586 | 602E-06 | 0000113406 | 052679748 | 1 |
| Ptk2 | ENSMUSG00000022607 | -2652043174 | 2973212447 | 1185308348 | 248E-06 | 316E-05 | -2246429292 | 3064473872 | 1810940927 | 408E-07 | 288E-05 | 0405613882 | 1 |
| Chrna2 | ENSMUSG00000022041 | -2634455648 | 1558175625 | 833907087 | 101E-05 | 804E-05 | -2304022385 | 1127763385 | 3851571422 | 000018299 | 0000989074 | 0330433263 | 1 |
| Gm38257 | ENSMUSG00000102258 | -2630099943 | 0903219803 | 597455608 | 370E-05 | 0000211526 | -2492549649 | 0756417102 | 4621286356 | 938E-05 | 0000633345 | 0137550294 | 1 |
| Nacc2 | ENSMUSG00000026932 | -2613149933 | 338957522 | 1778202043 | 473E-07 | 111E-05 | -3026025034 | 3338814633 | 2769246965 | 686E-08 | 117E-05 | -0412875101 | 0 |
| Irf7 | ENSMUSG00000025498 | -2610066663 | 5994267295 | 4954813755 | 641E-09 | 967E-07 | -2087542152 | 6085462364 | 1748380545 | 472E-07 | 322E-05 | 0522524511 | 1 |
| Ptp4a1 | ENSMUSG00000026064 | -2597318175 | 5563053505 | 3647690552 | 234E-08 | 189E-06 | -2236972732 | 5324999499 | 1108514506 | 307E-06 | 770E-05 | 0360345443 | 1 |
| Frat2 | ENSMUSG00000047604 | -258471047 | 6244152364 | 3004767088 | 530E-08 | 296E-06 | -2506822035 | 5946793925 | 5491084029 | 490E-05 | 0000411725 | 0077888435 | 1 |
| Dnajb4 | ENSMUSG00000028035 | -2565009609 | 5359686347 | 9672664899 | 562E-06 | 537E-05 | -2367059877 | 536256962 | 5653650243 | 439E-05 | 0000383225 | 0197949732 | 1 |
| Ddx60 | ENSMUSG00000037921 | -2557037727 | 5391872821 | 4480292673 | 982E-09 | 116E-06 | -2181785566 | 5350982264 | 5626699362 | 447E-05 | 0000387775 | 0375252161 | 1 |
| Tnfrsf12a | ENSMUSG00000023905 | -2556662947 | 2407451707 | 8222596862 | 107E-05 | 835E-05 | -2817023348 | 2285327759 | 573528789 | 415E-05 | 0000369991 | -0260360401 | 0 |
| Pacsin1 | ENSMUSG00000040276 | -2518201229 | 5927665543 | 3801766552 | 197E-08 | 170E-06 | -2267624594 | 5764333145 | 2118863091 | 212E-07 | 211E-05 | 0250576635 | 1 |
| Csrnp2 | ENSMUSG00000044636 | -250865359 | 3806464006 | 3877066594 | 181E-08 | 163E-06 | -2410699743 | 3765774094 | 1724223739 | 500E-07 | 329E-05 | 0097953847 | 1 |
| Adh1 | ENSMUSG00000074207 | -249614932 | 2265858168 | 5992308703 | 366E-05 | 000021007 | -2120724349 | 2422531819 | 8098628953 | 108E-05 | 0000163276 | 0375424971 | 1 |
| Dse | ENSMUSG00000039497 | -249130406 | 4324728915 | 1038254143 | 424E-06 | 449E-05 | -2114426553 | 4370602316 | 6840451371 | 210E-05 | 0000244897 | 0376877507 | 1 |
| Hexim1 | ENSMUSG00000048878 | -2477220512 | 7379248099 | 961477228 | 576E-06 | 546E-05 | -2072926785 | 702800771 | 6300450371 | 289E-05 | 000029358 | 0404293727 | 1 |
| Rps6ka2 | ENSMUSG00000023809 | -2465437593 | 2849146801 | 2025660553 | 276E-07 | 792E-06 | -2453034704 | 2877185257 | 1155931326 | 259E-06 | 705E-05 | 0012402889 | 1 |
| Emid1 | ENSMUSG00000034164 | -243943583 | 2140400051 | 7876503785 | 127E-05 | 946E-05 | -2828934771 | 2173111268 | 1470568623 | 967E-07 | 428E-05 | -0389498941 | 0 |
| Nfkbid | ENSMUSG00000036931 | -2429806815 | 6323259403 | 7439556914 | 159E-05 | 0000111549 | -2163445022 | 6178862355 | 557982408 | 461E-05 | 0000395881 | 0266361793 | 1 |
| Pdlim4 | ENSMUSG00000020388 | -2408734821 | 5077084793 | 1709943549 | 555E-07 | 120E-05 | -2552474224 | 5096752248 | 1363623789 | 132E-06 | 501E-05 | -0143739403 | 0 |
| Per2 | ENSMUSG00000055866 | -2407044958 | 2668856601 | 7480800324 | 155E-05 | 0000110229 | -2382627397 | 2884319361 | 2317681905 | 145E-07 | 168E-05 | 0024417561 | 1 |
| Cnksr3 | ENSMUSG00000015202 | -2396300123 | 3349938299 | 1775936529 | 475E-07 | 111E-05 | -2170010862 | 3342634097 | 130347209 | 159E-06 | 560E-05 | 0226289261 | 1 |
| Phlda1 | ENSMUSG00000020205 | -2393286589 | 2414099184 | 6178033195 | 325E-05 | 0000191633 | -2159865836 | 229665123 | 5293062119 | 563E-05 | 0000448733 | 0233420753 | 1 |
| Angpt1 | ENSMUSG00000022309 | -2386475965 | 1264926242 | 4085096774 | 0000152619 | 0000634452 | -3290516548 | 2271682782 | 2377114805 | 131E-07 | 160E-05 | -0904040583 | 0 |
| H1f2 | ENSMUSG00000036181 | -2344253322 | 5065789612 | 4131705828 | 0000146458 | 000061365 | -2564222953 | 4739300063 | 6467830145 | 261E-05 | 0000275384 | -0219969631 | 0 |
| Plp1 | ENSMUSG00000031425 | -2336078082 | 2126668048 | 9706635016 | 555E-06 | 536E-05 | -2007232802 | 2118233356 | 1017224414 | 435E-06 | 944E-05 | 032884528 | 1 |
| Rflnb | ENSMUSG00000020846 | -2335852784 | 7246640024 | 1088638395 | 223E-10 | 260E-07 | -2144362466 | 7133909978 | 449760018 | 0000103718 | 000067517 | 0191490318 | 1 |
| Prg4 | ENSMUSG00000006014 | -2307518333 | 1759035708 | 695378121 | 206E-05 | 0000136217 | -2742738673 | 1795426707 | 1274574143 | 174E-06 | 591E-05 | -043522034 | 0 |
| NA | ENSMUSG00000097479 | -2301550544 | 1100122044 | 4798678647 | 845E-05 | 0000399472 | -2476419919 | 0774421783 | 8454709883 | 912E-06 | 0000149488 | -0174869375 | 0 |
| Il6st | ENSMUSG00000021756 | -2289578251 | 7986004944 | 5821019413 | 323E-09 | 692E-07 | -2090599515 | 7980996376 | 1409609537 | 115E-06 | 466E-05 | 0198978736 | 1 |
| Ildr1 | ENSMUSG00000022900 | -2288873576 | 1261876262 | 7016135755 | 199E-05 | 0000132377 | -2657259989 | 1353254729 | 5701438817 | 425E-05 | 0000374916 | -0368386413 | 0 |
| Dntt | ENSMUSG00000025014 | -2284221259 | 2509533937 | 433202319 | 0000123225 | 0000535934 | -4008001411 | 2283335294 | 5888495449 | 375E-05 | 000034934 | -1723780152 | 0 |
| Rab3ip | ENSMUSG00000064181 | -227526246 | 5823814507 | 2593629938 | 983E-08 | 411E-06 | -2336905568 | 5907888564 | 1511277813 | 864E-07 | 402E-05 | -0061643108 | 0 |
| Macir | ENSMUSG00000044768 | -2267793939 | 4374643506 | 1900126365 | 359E-07 | 938E-06 | -2088521847 | 4633156716 | 8954785742 | 726E-06 | 0000127733 | 0179272092 | 1 |
| Gm43127 | ENSMUSG00000105140 | -2265626386 | 3030359212 | 1274370098 | 185E-06 | 265E-05 | -2052467833 | 2933536931 | 762569356 | 137E-05 | 0000188286 | 0213158553 | 1 |
| Lncpint | ENSMUSG00000044471 | -2260217229 | 243895685 | 6641997 | 246E-05 | 0000154968 | -2455857407 | 23318539 | 4336968805 | 0000118635 | 0000739889 | -0195640178 | 0 |
| Ccnd1 | ENSMUSG00000070348 | -2257591943 | 0798786814 | 4023904489 | 00001612 | 0000660246 | -3068846042 | 1713194901 | 5206986387 | 600E-05 | 0000466636 | -0811254099 | 0 |
| Hsd11b1 | ENSMUSG00000016194 | -2254491426 | 5628295456 | 2779045368 | 736E-08 | 352E-06 | -2172031387 | 5682513914 | 1710948177 | 516E-07 | 332E-05 | 0082460039 | 1 |
| Peg13 | ENSMUSG00000106847 | -224100573 | 5448483171 | 2985477329 | 545E-08 | 296E-06 | -2233140431 | 5308599558 | 4563696222 | 983E-05 | 0000651582 | 0007865299 | 1 |
| C230096K16Rik | ENSMUSG00000106205 | -2233884896 | 173461502 | 4776004275 | 860E-05 | 0000405426 | -2674351832 | 171398596 | 1268740255 | 177E-06 | 594E-05 | -0440466936 | 0 |
| Slfn5 | ENSMUSG00000054404 | -2233371522 | 6749677322 | 386246598 | 184E-08 | 163E-06 | -2423150346 | 6761869714 | 5703721898 | 424E-05 | 0000374852 | -0189778824 | 0 |
| Zfp36 | ENSMUSG00000044786 | -2212580774 | 8965826257 | 1246058162 | 203E-06 | 280E-05 | -2214364367 | 8727361101 | 3927070221 | 0000170563 | 0000940348 | -0001783593 | 0 |
| Nhsl1 | ENSMUSG00000039835 | -2205101549 | 2730437517 | 792958485 | 124E-05 | 929E-05 | -2436739072 | 2690494939 | 738027062 | 156E-05 | 000020363 | -0231637523 | 0 |
| Sorcs2 | ENSMUSG00000029093 | -2202204795 | 4012036151 | 4906729439 | 778E-05 | 0000378778 | -2163965498 | 3665294094 | 8823774164 | 770E-06 | 0000132913 | 0038239297 | 1 |
| Spsb1 | ENSMUSG00000039911 | -2198715015 | 4192245935 | 1932087313 | 335E-07 | 905E-06 | -2628391195 | 402660933 | 1467718803 | 974E-07 | 428E-05 | -042967618 | 0 |
| Ptprf | ENSMUSG00000033295 | -2198233586 | 2386767055 | 355445989 | 0000251442 | 0000921798 | -2369694547 | 2055025372 | 1817831802 | 401E-07 | 287E-05 | -0171460961 | 0 |
| Klf3 | ENSMUSG00000029178 | -2186335297 | 7514657362 | 1109036712 | 325E-06 | 378E-05 | -2066766617 | 7514392611 | 7615897708 | 915E-10 | 229E-06 | 011956868 | 1 |
| Ccdc141 | ENSMUSG00000044033 | -218598208 | 1005100535 | 6843103602 | 220E-05 | 0000142936 | -2210037629 | 1074242278 | 8338896093 | 964E-06 | 0000154018 | -0024055549 | 0 |
| Dnajb1 | ENSMUSG00000005483 | -2178967616 | 7532955844 | 1106137335 | 328E-06 | 380E-05 | -2396021885 | 7777244956 | 165838685 | 588E-07 | 338E-05 | -0217054269 | 0 |
| Gabrr2 | ENSMUSG00000023267 | -2172429188 | 3740367174 | 2579566214 | 101E-07 | 413E-06 | -2488970063 | 3875360134 | 5321453311 | 552E-05 | 0000443222 | -0316540875 | 0 |
| Foxp3 | ENSMUSG00000039521 | -2155688359 | 2574582454 | 1263326792 | 192E-06 | 272E-05 | -2049466792 | 2917826602 | 6386172042 | 275E-05 | 0000285068 | 0106221567 | 1 |
| Hdac9 | ENSMUSG00000004698 | -2153592442 | 1845966678 | 484515993 | 815E-05 | 0000390507 | -311327945 | 2029036571 | 425367606 | 0000127409 | 0000771898 | -0959687008 | 0 |
| Plekha1 | ENSMUSG00000040268 | -213189774 | 6063566801 | 4642673062 | 845E-09 | 108E-06 | -2013194886 | 6100631214 | 2164580808 | 194E-07 | 197E-05 | 0118702854 | 1 |
| Trim2 | ENSMUSG00000027993 | -2122020624 | 1200223001 | 5755228369 | 427E-05 | 0000236954 | -2119117949 | 1624095827 | 7510346666 | 146E-05 | 0000195148 | 0002902675 | 1 |
| Acvrl1 | ENSMUSG00000000530 | -2095195357 | 531054418 | 4801026001 | 843E-05 | 0000399247 | -2408177392 | 5005198778 | 1697043535 | 534E-07 | 334E-05 | -0312982035 | 0 |
| Smad7 | ENSMUSG00000025880 | -2074419688 | 7856767722 | 25760826 | 101E-07 | 413E-06 | -2120813152 | 7663369239 | 4800668073 | 814E-05 | 0000572947 | -0046393464 | 0 |
| A430072P03Rik | ENSMUSG00000090030 | 200892922 | 1890541239 | 657105324 | 257E-05 | 0000159378 | 2393260026 | 1881045974 | 1027691945 | 417E-06 | 925E-05 | 0384330806 | 1 |
| Gm14321 | ENSMUSG00000084797 | 2045406072 | 2514354834 | 1089840398 | 349E-06 | 395E-05 | 2024773832 | 2786268529 | 5072113497 | 662E-05 | 000049959 | -002063224 | 0 |
| Kctd17 | ENSMUSG00000033287 | 2086820396 | 4143203609 | 23832375 | 140E-07 | 517E-06 | 2091934224 | 4185726086 | 2755365172 | 701E-08 | 117E-05 | 0005113828 | 1 |
| Tspoap1 | ENSMUSG00000034156 | 2096335718 | 4735163957 | 1233923667 | 211E-06 | 286E-05 | 2314707671 | 4986480598 | 4908543593 | 749E-05 | 0000542285 | 0218371953 | 1 |
| Slc36a3os | ENSMUSG00000085531 | 2099270135 | 1397225807 | 50379957 | 705E-05 | 000035084 | 2317216172 | 1206073723 | 8173739183 | 104E-05 | 0000160199 | 0217946037 | 1 |
| Spo11 | ENSMUSG00000005883 | 2106370478 | 37697114 | 1209388729 | 229E-06 | 302E-05 | 2126208255 | 3776113009 | 1017205679 | 435E-06 | 944E-05 | 0019837777 | 1 |
| Arhgap33 | ENSMUSG00000036882 | 2107349645 | 418710049 | 1070868584 | 374E-06 | 414E-05 | 2111191413 | 4621196471 | 5140773796 | 629E-05 | 0000482707 | 0003841768 | 1 |
| E2f2 | ENSMUSG00000018983 | 2124871176 | 6936767906 | 2517483622 | 111E-07 | 447E-06 | 2120213253 | 7062309628 | 7567217166 | 142E-05 | 0000191793 | -0004657923 | 0 |
| Gprin3 | ENSMUSG00000045441 | 2128254685 | 5630774046 | 4010919057 | 157E-08 | 150E-06 | 2214749165 | 5696822568 | 9420947318 | 366E-10 | 229E-06 | 008649448 | 1 |
| Zbtb42 | ENSMUSG00000037638 | 2151777023 | 4021752226 | 1760059643 | 493E-07 | 114E-05 | 2015489607 | 4080165178 | 378262393 | 183E-08 | 616E-06 | -0136287416 | 0 |
| Tmtc2 | ENSMUSG00000036019 | 2167393067 | 1873151633 | 1014156771 | 465E-06 | 477E-05 | 2438923394 | 2082134193 | 6011902581 | 347E-05 | 0000329158 | 0271530327 | 1 |
| Fabp5 | ENSMUSG00000027533 | 2183452087 | 3835001023 | 9815866391 | 530E-06 | 520E-05 | 2171377776 | 4090277677 | 9177090327 | 658E-06 | 000011909 | -0012074311 | 0 |
| Lgalsl | ENSMUSG00000042363 | 2206257621 | 2786782866 | 9246092324 | 673E-06 | 611E-05 | 2104011239 | 2976481816 | 1221992879 | 207E-06 | 649E-05 | -0102246382 | 0 |
| Tmem237 | ENSMUSG00000038079 | 2213689471 | 2735570735 | 6954661547 | 206E-05 | 0000136217 | 2232319816 | 3115461952 | 6710434136 | 227E-05 | 0000256274 | 0018630345 | 1 |
| Ar | ENSMUSG00000046532 | 2226509522 | 5483502486 | 1536533506 | 862E-07 | 156E-05 | 2395575729 | 5684126357 | 5804316169 | 294E-09 | 242E-06 | 0169066207 | 1 |
| Lxn | ENSMUSG00000047557 | 2232293853 | 3022707928 | 7617253582 | 145E-05 | 0000104487 | 2194260694 | 33107527 | 1139300414 | 275E-06 | 726E-05 | -0038033159 | 0 |
| S100a10 | ENSMUSG00000041959 | 2247991864 | 8093284864 | 2257903275 | 175E-07 | 595E-06 | 2168210882 | 8129041991 | 2165987116 | 193E-07 | 197E-05 | -0079780982 | 0 |
| Tanc2 | ENSMUSG00000053580 | 2273360718 | 3974992297 | 2222944983 | 187E-07 | 616E-06 | 2443135026 | 3890681867 | 5380381827 | 530E-05 | 0000431662 | 0169774308 | 1 |
| Epas1 | ENSMUSG00000024140 | 2281417843 | 6022301037 | 3319052883 | 349E-08 | 233E-06 | 2736471188 | 6325806282 | 8948204574 | 728E-06 | 0000127743 | 0455053345 | 1 |
| Tmem220 | ENSMUSG00000050270 | 2281838057 | 0807134587 | 7432492874 | 159E-05 | 0000111694 | 2590829788 | 0700684443 | 5958552611 | 359E-05 | 0000336055 | 0308991731 | 1 |
| Irf4 | ENSMUSG00000021356 | 2291130329 | 6564033821 | 1638387465 | 662E-07 | 133E-05 | 2266122671 | 6752829114 | 4723374727 | 865E-05 | 000059646 | -0025007658 | 0 |
| E230034D01Rik | ENSMUSG00000111283 | 2293052334 | 3045291464 | 8743648009 | 840E-06 | 705E-05 | 2358237361 | 307734986 | 653937566 | 250E-05 | 0000270722 | 0065185027 | 1 |
| Kank3 | ENSMUSG00000042099 | 2295963144 | 2426130957 | 1219966064 | 221E-06 | 295E-05 | 2130566976 | 2597301295 | 5543694691 | 473E-05 | 0000400981 | -0165396168 | 0 |
| Gm42047 | ENSMUSG00000110631 | 2313961122 | 7437583987 | 3852923248 | 0000188537 | 0000744261 | 306456629 | 8369114671 | 1344222072 | 140E-06 | 519E-05 | 0750605168 | 1 |
| Dubr | ENSMUSG00000022639 | 2322861115 | 4838836163 | 6085396606 | 267E-09 | 619E-07 | 2376646356 | 5058739783 | 3567485584 | 235E-08 | 683E-06 | 0053785241 | 1 |
| Gm8493 | ENSMUSG00000106028 | 2323347999 | 2042007021 | 936621611 | 639E-06 | 587E-05 | 2434675691 | 2051789232 | 137190325 | 129E-06 | 496E-05 | 0111327692 | 1 |
| Hipk2 | ENSMUSG00000061436 | 2351534502 | 6611730572 | 3838416363 | 189E-08 | 165E-06 | 2012214437 | 6480828692 | 1095283457 | 323E-06 | 786E-05 | -0339320065 | 0 |
| Ggt7 | ENSMUSG00000027603 | 2361512877 | 4375471872 | 3545469463 | 264E-08 | 198E-06 | 2258884841 | 4161499551 | 1009153913 | 449E-06 | 962E-05 | -0102628036 | 0 |
| Aldoa | ENSMUSG00000030695 | 2362307779 | 9741945181 | 9920561619 | 508E-06 | 506E-05 | 2087184123 | 9519309853 | 7148707493 | 177E-05 | 0000219944 | -0275123656 | 0 |
| Zmat3 | ENSMUSG00000027663 | 2366688361 | 5666575217 | 2042627268 | 266E-07 | 771E-06 | 2475347306 | 5845513583 | 1397959589 | 119E-06 | 480E-05 | 0108658945 | 1 |
| Capg | ENSMUSG00000056737 | 2368727433 | 6707879426 | 1513029027 | 919E-07 | 162E-05 | 2033601703 | 6739143783 | 5882903797 | 377E-05 | 0000349982 | -033512573 | 0 |
| | ENSMUSG00000095742 | 2381782752 | 5281531965 | 1601311819 | 728E-07 | 142E-05 | 2094064743 | 497840724 | 1631246862 | 630E-07 | 350E-05 | -0287718009 | 0 |
| Sesn2 | ENSMUSG00000028893 | 2385784666 | 4459868866 | 2004517148 | 288E-07 | 819E-06 | 2278359008 | 4785188445 | 5423469456 | 514E-05 | 0000423625 | -0107425658 | 0 |
| Mgmt | ENSMUSG00000054612 | 2386946124 | 276208676 | 1579046176 | 771E-07 | 146E-05 | 223341208 | 2525163946 | 6492842907 | 257E-05 | 0000272953 | -0153534044 | 0 |
| Dhcr24 | ENSMUSG00000034926 | 2402294023 | 6433767388 | 1930584691 | 336E-07 | 906E-06 | 2209104169 | 6455328031 | 1246764495 | 190E-06 | 622E-05 | -0193189854 | 0 |
| Abhd4 | ENSMUSG00000040997 | 2429407012 | 3811399115 | 1771188043 | 480E-07 | 112E-05 | 2136224762 | 42440431 | 4290993215 | 0000123382 | 0000756574 | -029318225 | 0 |
| Fbxo27 | ENSMUSG00000037463 | 2432858792 | 2814537706 | 1071266496 | 374E-06 | 414E-05 | 2473428929 | 2977004827 | 5481577695 | 493E-05 | 0000412413 | 0040570137 | 1 |
| Slc2a3 | ENSMUSG00000003153 | 2443557734 | 7798434856 | 2065882878 | 254E-07 | 752E-06 | 2088818854 | 7372229742 | 7525048699 | 145E-05 | 0000194561 | -035473888 | 0 |
| Zfp760 | ENSMUSG00000067928 | 246163207 | 2912380279 | 187759125 | 377E-07 | 967E-06 | 2101602682 | 3461786252 | 5012181498 | 692E-05 | 0000515391 | -0360029388 | 0 |

| | | | | | | | | | | | | | |
|---|---|---|---|---|---|---|---|---|---|---|---|---|---|
| Abhd14b | ENSMUSG00000042073 | 2462799337 | 3130962806 | 2048749976 | 263E-07 | 763E-06 | 2271586335 | 2980727719 | 1866009077 | 360E-07 | 267E-05 | -0191213002 | 0 |
| Tnfrsf4 | ENSMUSG00000029075 | 2463871738 | 5161244467 | 701502562 | 146E-09 | 451E-07 | 220623608 | 5139183011 | 9654758957 | 537E-06 | 0000104804 | -0257635658 | 0 |
| Bbs12 | ENSMUSG00000051444 | 2468206723 | 378741055 | 2602363129 | 969E-08 | 408E-06 | 235182947 | 3773158908 | 3541503874 | 242E-08 | 683E-06 | -0116377253 | 0 |
| Il3ra | ENSMUSG00000068758 | 2474615836 | 524968085 | 5581970343 | 386E-09 | 727E-07 | 2286998517 | 505533051 | 2485795428 | 108E-07 | 152E-05 | -0187617319 | 0 |
| Armcx4 | ENSMUSG00000049804 | 2475355191 | 36521626 | 1902825912 | 357E-07 | 936E-06 | 2171894416 | 365487919 | 2033798718 | 251E-07 | 223E-05 | -0303460775 | 0 |
| Ero1a | ENSMUSG00000021831 | 2481189505 | 7129364888 | 2021629958 | 278E-07 | 794E-06 | 2166011101 | 6915694449 | 1604891028 | 673E-07 | 353E-05 | -0315178404 | 0 |
| Repin1 | ENSMUSG00000052751 | 24823762 | 3111452144 | 1589901782 | 750E-07 | 144E-05 | 2229378598 | 3362276729 | 1496155654 | 900E-07 | 413E-05 | -0252997602 | 0 |
| Podnl1 | ENSMUSG00000012889 | 2496456101 | 5442465448 | 2786971686 | 727E-08 | 352E-06 | 2269298924 | 546971349 | 1280889472 | 170E-06 | 582E-05 | -0227157177 | 0 |
| Glipr2 | ENSMUSG00000028480 | 2504628022 | 7298680918 | 7516371473 | 109E-09 | 396E-07 | 2458716595 | 7455337004 | 4569223715 | 818E-09 | 472E-06 | -0045911427 | 0 |
| Slc9a5 | ENSMUSG00000014786 | 2505361341 | 2234483528 | 8643256926 | 879E-06 | 726E-05 | 3056294909 | 2882207556 | 4786720691 | 823E-05 | 0000576368 | 0550933568 | 1 |
| Slc52a3 | ENSMUSG00000027463 | 2511902465 | 2740092798 | 1934283154 | 334E-07 | 905E-06 | 2465988979 | 2947387725 | 1548454013 | 781E-07 | 385E-05 | -0045913486 | 0 |
| Gm16976 | ENSMUSG00000085500 | 2522169168 | 0784868243 | 5273820029 | 594E-05 | 0000304759 | 3480183491 | 1192835921 | 9774627184 | 511E-06 | 0000102394 | 0958014323 | 1 |
| Ier5l | ENSMUSG00000089762 | 2527100881 | 2594767751 | 3810921453 | 0000196107 | 0000764196 | 2503287118 | 2492073127 | 9076100149 | 688E-06 | 0000123376 | -0023813763 | 0 |
| S100a1 | ENSMUSG00000044080 | 2573184495 | 193167956 | 1406474344 | 124E-06 | 200E-05 | 2874424701 | 2400480804 | 5098803424 | 649E-05 | 0000493184 | 0301240206 | 1 |
| 9230114K14Rik | ENSMUSG00000097145 | 2583130918 | 1430636954 | 1139975103 | 291E-06 | 350E-05 | 3237932062 | 2163463697 | 5088522305 | 654E-05 | 0000495707 | 0654801144 | 1 |
| Dglucy | ENSMUSG00000021185 | 2590790904 | 613345622 | 1194724137 | 150E-10 | 257E-07 | 2601325226 | 612152305 | 296480644 | 514E-08 | 945E-06 | 0010534322 | 1 |
| Gapdh | ENSMUSG00000057666 | 2594144196 | 1043452863 | 2779949209 | 735E-08 | 352E-06 | 2327363643 | 1027605326 | 1049002817 | 384E-06 | 879E-05 | -0266780553 | 0 |
| Kif22 | ENSMUSG00000030677 | 2597446147 | 519570867 | 1470472919 | 103E-06 | 177E-05 | 2426245063 | 5805438928 | 3873441687 | 0000179279 | 0000976852 | -0171201084 | 0 |
| Insl3 | ENSMUSG00000079019 | 2606649527 | 4509997028 | 1721668414 | 540E-07 | 118E-05 | 2332380835 | 4344214824 | 9497069511 | 573E-06 | 0000109354 | -0274268692 | 0 |
| Oit3 | ENSMUSG00000009654 | 2632934093 | 1054665051 | 6048856903 | 353E-05 | 0000204374 | 2161578095 | 0785330011 | 6911614417 | 202E-05 | 0000239418 | -0471355998 | 0 |
| Gm17173 | ENSMUSG00000090709 | 2634093587 | 2808186861 | 1556914531 | 817E-07 | 152E-05 | 313710466 | 2974561665 | 1494640626 | 904E-07 | 413E-05 | 0503011073 | 1 |
| Carhsp1 | ENSMUSG00000008393 | 2643646046 | 5779947254 | 8393292559 | 679E-10 | 347E-07 | 2038263425 | 5814093841 | 7736505412 | 130E-05 | 0000181848 | -0605382621 | 0 |
| Tspan4 | ENSMUSG00000025511 | 2648821508 | 3621329037 | 3351889394 | 335E-08 | 227E-06 | 2119920672 | 3591481677 | 1995906889 | 272E-07 | 231E-05 | -0528900836 | 0 |
| Castor1 | ENSMUSG00000020424 | 2659186332 | 602682635 | 3988471522 | 161E-08 | 150E-06 | 2614641144 | 5839751625 | 2307126986 | 148E-07 | 168E-05 | -0044545188 | 0 |
| Myo3b | ENSMUSG00000042064 | 2671045697 | 1173422801 | 8291588788 | 104E-05 | 816E-05 | 2626073781 | 1694992947 | 8049027763 | 111E-05 | 0000165905 | -0044971916 | 0 |
| Mcub | ENSMUSG00000027994 | 2676069345 | 3286591229 | 1250299052 | 200E-06 | 278E-05 | 2138495374 | 3366421332 | 1119708543 | 295E-06 | 755E-05 | -0537573971 | 0 |
| Neil2 | ENSMUSG00000035121 | 267910647 | 2685275934 | 1181567213 | 252E-06 | 318E-05 | 2330688678 | 2729121714 | 6814768864 | 213E-05 | 0000246855 | -0348417792 | 0 |
| Klk9 | ENSMUSG00000047884 | 2683054936 | 0752125313 | 6208057091 | 319E-05 | 0000189058 | 3000348323 | 0746612452 | 1255627236 | 185E-06 | 611E-05 | 0317293387 | 1 |
| Alpk2 | ENSMUSG00000032845 | 2696411562 | 1472514458 | 5533058589 | 495E-05 | 0000265995 | 2536521212 | 1604436796 | 1221513139 | 207E-06 | 649E-05 | -015989035 | 0 |
| Nqo2 | ENSMUSG00000046949 | 2696971615 | 4457468531 | 5278515032 | 490E-09 | 857E-07 | 247217518 | 4571792308 | 8768432147 | 789E-06 | 0000134955 | -0224796435 | 0 |
| Kcnq5 | ENSMUSG00000028033 | 2706078813 | 3667274822 | 2327248318 | 155E-07 | 548E-06 | 2668499114 | 386157957 | 3420811062 | 280E-08 | 740E-06 | -0037579699 | 0 |
| Ndrg1 | ENSMUSG00000005125 | 2711921761 | 6039846223 | 1031630193 | 435E-06 | 459E-05 | 2164121173 | 5548400281 | 8411870022 | 931E-06 | 0000151167 | -0547800588 | 0 |
| Bhlhe40 | ENSMUSG00000030103 | 2712263322 | 6138919201 | 368365984 | 225E-08 | 183E-06 | 2601507895 | 6386491633 | 1683367762 | 553E-07 | 337E-05 | -0110755427 | 0 |
| Hgfac | ENSMUSG00000029102 | 2714128842 | 0858181762 | 6367549981 | 290E-05 | 0000175518 | 2476008968 | 0973503874 | 5129779205 | 634E-05 | 0000485394 | -0238119874 | 0 |
| Gpr179 | ENSMUSG00000070337 | 2730339124 | 1819827385 | 1205448485 | 232E-06 | 304E-05 | 2024274067 | 1881094527 | 4508700869 | 0000102775 | 0000671637 | -0706065057 | 0 |
| 4933433G15Rik | ENSMUSG00000097021 | 2742031874 | 1184933549 | 6311032319 | 300E-05 | 0000180027 | 2782847252 | 1307609397 | 8922008211 | 736E-06 | 0000129061 | 0040815378 | 1 |
| Gpt2 | ENSMUSG00000031700 | 2762467492 | 1605670972 | 1403566808 | 125E-06 | 202E-05 | 2560134788 | 1853401453 | 4053948384 | 0000151933 | 0000869655 | -0202332704 | 0 |
| Sema7a | ENSMUSG00000038264 | 2773013527 | 3128333025 | 8673317977 | 867E-06 | 719E-05 | 2374228017 | 3602019944 | 4863857291 | 775E-05 | 0000554764 | -039878551 | 0 |
| Crmp1 | ENSMUSG00000029121 | 2782874276 | 5490457997 | 3862924007 | 184E-08 | 163E-06 | 2701981969 | 5435987474 | 1612678095 | 660E-07 | 353E-05 | -0080892307 | 0 |
| Glis2 | ENSMUSG00000014303 | 2810341398 | 1843924543 | 8017171777 | 118E-05 | 900E-05 | 2155928966 | 190142679 | 8309772398 | 977E-06 | 0000155285 | -0654412432 | 0 |
| Ncs1 | ENSMUSG00000062661 | 2816204945 | 1882794233 | 9819259253 | 530E-06 | 520E-05 | 2364385653 | 2176671009 | 1053331666 | 378E-06 | 868E-05 | -0451819292 | 0 |
| Plekhf1 | ENSMUSG00000074170 | 2835593838 | 1053254963 | 1335689137 | 153E-06 | 234E-05 | 2089980509 | 1418013065 | 4044442459 | 0000153239 | 0000875124 | -0745613329 | 0 |
| Trim46 | ENSMUSG00000042766 | 2837103002 | 3377517168 | 2333926347 | 153E-07 | 544E-06 | 2297809667 | 3342937683 | 3123279016 | 413E-08 | 863E-06 | -0539293335 | 0 |
| Gm39323 | ENSMUSG00000111229 | 2841845317 | 5740259934 | 6814232966 | 165E-09 | 482E-07 | 2958238892 | 5838655682 | 411657051 | 128E-08 | 573E-06 | 0116393575 | 1 |
| Hic1 | ENSMUSG00000043099 | 2849335727 | 3848649464 | 9370635761 | 638E-06 | 587E-05 | 3156752737 | 4110823211 | 1510952615 | 864E-07 | 402E-05 | 030741701 | 1 |
| Raph1 | ENSMUSG00000026014 | 2850872256 | 495739248 | 4361732942 | 110E-08 | 121E-06 | 2474375593 | 5155517115 | 1198320822 | 224E-06 | 652E-05 | -0376496663 | 0 |
| D930048N14Rik | ENSMUSG00000052563 | 2851663634 | 2454368269 | 8281801701 | 104E-05 | 817E-05 | 2554930881 | 2750784455 | 1301659922 | 160E-06 | 561E-05 | -0296732753 | 0 |
| Piwil2 | ENSMUSG00000033644 | 2859558926 | 3311059362 | 1905834532 | 355E-07 | 934E-06 | 2797321123 | 30754045 | 1295230559 | 163E-06 | 568E-05 | -0062237803 | 0 |
| Gemin8 | ENSMUSG00000040621 | 2860060023 | 3839910144 | 4284531533 | 119E-08 | 128E-06 | 2253826199 | 3803041488 | 2413057098 | 123E-07 | 158E-05 | -0606233824 | 0 |
| Afdn | ENSMUSG00000068036 | 2896117655 | 4377068077 | 2372609126 | 143E-07 | 525E-06 | 2181513616 | 4424201067 | 1110491967 | 305E-06 | 768E-05 | -0714604039 | 0 |
| Ica1l | ENSMUSG00000026018 | 2905375259 | 2513547013 | 1610512483 | 711E-07 | 139E-05 | 2550652582 | 2278566446 | 9217242575 | 646E-06 | 0000118299 | -0354722677 | 0 |
| Fggy | ENSMUSG00000028573 | 290582621 | 2626344952 | 2090073809 | 242E-07 | 726E-06 | 238077939 | 2467663773 | 996766653 | 472E-06 | 992E-05 | -052504682 | 0 |
| Fah | ENSMUSG00000030630 | 2909788322 | 2501271119 | 7152405824 | 185E-05 | 0000124926 | 2321530926 | 2405208568 | 114093653 | 273E-06 | 724E-05 | -0588257396 | 0 |
| Bcl2l2 | ENSMUSG00000089682 | 2921543247 | 3576845665 | 2834450443 | 678E-08 | 341E-06 | 2687103104 | 3634328374 | 143989021 | 105E-06 | 444E-05 | -0234440143 | 0 |
| Yes1 | ENSMUSG00000014932 | 2921663819 | 2021867999 | 1717123976 | 546E-07 | 119E-05 | 2399026554 | 2086329571 | 2272691328 | 158E-07 | 171E-05 | -0522637265 | 0 |
| Ecm1 | ENSMUSG00000028108 | 293217829 | 7639467272 | 1451519846 | 651E-11 | 182E-07 | 226577554 | 6882087517 | 5490957978 | 490E-05 | 0000411725 | -066640275 | 0 |
| Vim | ENSMUSG00000026728 | 2946104728 | 1064453124 | 2662407624 | 881E-08 | 385E-06 | 2565807452 | 1055687818 | 681968773 | 213E-05 | 0000246855 | -0380297276 | 0 |
| Ttc16 | ENSMUSG00000039021 | 2952928576 | 0431040396 | 1095538024 | 341E-06 | 390E-05 | 211233515 | 0584961761 | 4464940115 | 0000106555 | 000068799 | -0840593426 | 0 |
| Ccr8 | ENSMUSG00000042262 | 2957184566 | 3689785008 | 1391178778 | 130E-06 | 208E-05 | 2824550157 | 3358245665 | 5570931999 | 464E-05 | 0000396402 | -0132634409 | 0 |
| Slc22a15 | ENSMUSG00000033147 | 2966962698 | 4172414928 | 2631576297 | 925E-08 | 393E-06 | 234915237 | 3925768422 | 5816764311 | 393E-05 | 0000358984 | -0617810328 | 0 |
| Itih5 | ENSMUSG00000025780 | 2981008685 | 6676804553 | 2126481686 | 225E-07 | 693E-06 | 2260545552 | 6215204034 | 3870101852 | 0000179839 | 0000978149 | -0720463133 | 0 |
| Tmem231 | ENSMUSG00000031951 | 2989039071 | 1406562773 | 102818036 | 440E-06 | 461E-05 | 2232914754 | 1617745518 | 5251954581 | 580E-05 | 0000457072 | -0756124317 | 0 |
| Otos | ENSMUSG00000044055 | 2989682028 | 0565143986 | 8793862466 | 821E-06 | 699E-05 | 3004911168 | 0957777588 | 5518135512 | 481E-05 | 0000405568 | 001522914 | 1 |
| Nlgn2 | ENSMUSG00000051790 | 2991687343 | 4536482011 | 1820173844 | 429E-07 | 103E-05 | 2500135259 | 4213058773 | 6478256344 | 260E-05 | 0000274399 | -0491552084 | 0 |
| Impa2 | ENSMUSG00000024525 | 2997400029 | 5883639932 | 1787475428 | 463E-07 | 109E-05 | 2446612599 | 5581904142 | 1233345279 | 199E-06 | 635E-05 | -055078743 | 0 |
| Il4 | ENSMUSG00000000869 | 3015038553 | 1594041927 | 15451723 | 843E-07 | 155E-05 | 278781485 | 1506721157 | 1218446113 | 209E-06 | 651E-05 | -0227223703 | 0 |
| 5730409E04Rik | ENSMUSG00000073755 | 3018578304 | 1288562928 | 9461558152 | 614E-06 | 572E-05 | 3211527111 | 0904189996 | 5504628134 | 486E-05 | 0000408697 | 0192948807 | 1 |
| Cish | ENSMUSG00000032578 | 3038403595 | 5361931839 | 1649437919 | 644E-07 | 131E-05 | 312450187 | 5497635929 | 1604426595 | 674E-07 | 353E-05 | 0086098275 | 1 |
| Slc25a24 | ENSMUSG00000040322 | 3044148948 | 6082876761 | 7860229224 | 898E-10 | 361E-07 | 2447319277 | 6112228626 | 2448815611 | 115E-07 | 157E-05 | -0596829671 | 0 |
| Myl6b | ENSMUSG00000039824 | 3045598234 | 2586690543 | 8630669635 | 885E-06 | 728E-05 | 2227026228 | 2827288559 | 1119720696 | 295E-06 | 755E-05 | -0818572006 | 0 |
| Tmbim1 | ENSMUSG00000006301 | 3063837408 | 3490143407 | 292944123 | 590E-08 | 312E-06 | 228456454 | 3544670024 | 9542238073 | 563E-06 | 000010763 | -0779272868 | 0 |
| Cldn12 | ENSMUSG00000046798 | 3069479922 | 2066438111 | 1438369944 | 113E-06 | 189E-05 | 2617813941 | 2290645353 | 1297238987 | 162E-06 | 567E-05 | -0451665981 | 0 |
| Mrgpre | ENSMUSG00000048965 | 3092462577 | 2869999552 | 1207055654 | 231E-06 | 303E-05 | 243441207 | 2966258452 | 9747216655 | 517E-06 | 0000103386 | -0658050507 | 0 |
| Cpa3 | ENSMUSG00000001865 | 3102674267 | 4064584095 | 3359459306 | 332E-08 | 227E-06 | 3042592903 | 474294797 | 8120121422 | 107E-05 | 0000163012 | -0060081364 | 0 |
| Emp1 | ENSMUSG00000030208 | 3119738991 | 3181992248 | 3950043955 | 0000172365 | 0000696901 | 2038323285 | 2983833238 | 4885951289 | 762E-05 | 00005482 | -1081415706 | 0 |
| Rbpj | ENSMUSG00000039191 | 3135087694 | 7773066158 | 983816445 | 344E-10 | 268E-07 | 2770715751 | 7600461511 | 4477113554 | 892E-09 | 492E-06 | -0364371943 | 0 |
| Prrg4 | ENSMUSG00000027171 | 3139642415 | 3682704573 | 2645538495 | 905E-08 | 391E-06 | 3135958228 | 398241373 | 9038296295 | 699E-06 | 0000124532 | -0003684187 | 0 |
| C130083M11Rik | ENSMUSG00000106464 | 3154205569 | 142666812 | 119022462 | 244E-06 | 313E-05 | 3035576791 | 2261531743 | 4062299665 | 0000150797 | 0000865319 | -0118628778 | 0 |
| Glrx | ENSMUSG00000021591 | 3156500259 | 5660825385 | 3730959078 | 213E-08 | 179E-06 | 2570523997 | 5524097439 | 265597641 | 819E-08 | 129E-05 | -0585976262 | 0 |
| Syne3 | ENSMUSG00000054150 | 3168578248 | 6545035509 | 7664901929 | 100E-09 | 380E-07 | 2714828951 | 6579966741 | 1217230852 | 210E-06 | 651E-05 | -0453749297 | 0 |
| Gm28875 | ENSMUSG00000100975 | 3174298589 | 4323467234 | 3603745892 | 247E-08 | 189E-06 | 2701533273 | 4233110603 | 1015366918 | 438E-06 | 947E-05 | -0472765316 | 0 |
| Srgap3 | ENSMUSG00000030257 | 319937929 | 4516100764 | 1240065478 | 207E-06 | 283E-05 | 2221884411 | 4551236351 | 444038283 | 0000108751 | 0000697291 | -0977494879 | 0 |
| H1f0 | ENSMUSG00000096210 | 3202906651 | 6220305878 | 1290133601 | 176E-06 | 257E-05 | 2982323076 | 6162515641 | 1241111058 | 194E-06 | 625E-05 | -0220583575 | 0 |
| AA467197 | ENSMUSG00000033213 | 3205781664 | 1104966143 | 8813044457 | 814E-06 | 697E-05 | 3761161261 | 1095184424 | 1130183429 | 284E-06 | 742E-05 | 0555379597 | 1 |
| Pcgf2 | ENSMUSG00000018537 | 3216454587 | 3301394191 | 2152460866 | 214E-07 | 673E-06 | 2738787811 | 3514776148 | 1643175092 | 611E-07 | 343E-05 | -0477666776 | 0 |
| Paqr4 | ENSMUSG00000023909 | 3225025033 | 3604698177 | 1600056954 | 730E-07 | 142E-05 | 3107276113 | 3982160294 | 5202279181 | 602E-05 | 0000467424 | -011774892 | 0 |
| Ptgir | ENSMUSG00000043017 | 3226272038 | 4997448356 | 195072599 | 322E-07 | 884E-06 | 286118531 | 496449953 | 8631434606 | 840E-06 | 0000141457 | -0365086728 | 0 |
| Nol4l | ENSMUSG00000061411 | 3230457551 | 3441502069 | 3235830091 | 388E-08 | 249E-06 | 2613964964 | 3388840079 | 8396902426 | 938E-06 | 0000152037 | -0616492587 | 0 |
| Ifitm1 | ENSMUSG00000025491 | 3242213809 | 3806809285 | 5244745624 | 606E-05 | 0000310068 | 261232625 | 4433729283 | 629303109 | 291E-05 | 0000294428 | -0629887559 | 0 |
| Eif4ebp1 | ENSMUSG00000031490 | 3251191115 | 4622586523 | 2260361253 | 175E-07 | 594E-06 | 2366469172 | 4521726274 | 8458642953 | 911E-06 | 0000149486 | -0884721943 | 0 |
| Fbxo44 | ENSMUSG00000029001 | 3259970803 | 2702447116 | 1251214152 | 199E-06 | 277E-05 | 3220144887 | 2491968021 | 1905464206 | 330E-07 | 252E-05 | -0039825916 | 0 |
| Smo | ENSMUSG00000001761 | 3274036293 | 5031921734 | 8129890464 | 778E-10 | 347E-07 | 2556371021 | 4677639011 | 1128494507 | 286E-06 | 744E-05 | -0717665272 | 0 |
| Lmtk3 | ENSMUSG00000062044 | 3278158628 | 2925142292 | 2861124386 | 651E-08 | 332E-06 | 3070364966 | 3143731578 | 1574658652 | 729E-07 | 368E-05 | -0207793662 | 0 |
| Ly6a | ENSMUSG00000075602 | 3283635723 | 8264707238 | 7041677439 | 144E-09 | 451E-07 | 3094161746 | 818559432 | 6518972874 | 179E-09 | 229E-06 | -0189473977 | 0 |
| Rapsn | ENSMUSG00000002104 | 3301552767 | 0685903459 | 6836133005 | 220E-05 | 0000143342 | 3069735098 | 111622405 | 8573428394 | 863E-06 | 0000144403 | -0231817669 | 0 |
| Hip1 | ENSMUSG00000039959 | 3304969935 | 4317572279 | 1587013387 | 755E-07 | 144E-05 | 2254944247 | 458866967 | 5448570354 | 505E-05 | 0000418532 | -1050025688 | 0 |
| Ston2 | ENSMUSG00000020961 | 3332883528 | 3442508316 | 9953351891 | 502E-06 | 503E-05 | 2956014081 | 3397588524 | 8620913675 | 844E-06 | 0000141457 | -0376869447 | 0 |
| 9530082P21Rik | ENSMUSG00000097006 | 3333250168 | 2071295889 | 1208425485 | 230E-06 | 302E-05 | 3231860849 | 1980085908 | 8270365787 | 996E-06 | 0000156583 | -0101389319 | 0 |
| Ccdc40 | ENSMUSG00000039963 | 3336421307 | 2788430029 | 2370467202 | 143E-07 | 525E-06 | 3026042075 | 3023686918 | 1038670839 | 400E-06 | 901E-05 | -0310379232 | 0 |
| B3galt5 | ENSMUSG00000074892 | 3350713779 | 1344192765 | 1095317682 | 342E-06 | 390E-05 | 3479584425 | 1577863086 | 9887583064 | 488E-06 | 0000101284 | 0128870646 | 1 |
| Kremen2 | ENSMUSG00000040680 | 3354029987 | 3588121877 | 1778729441 | 472E-07 | 111E-05 | 2925149337 | 3188392831 | 1378463797 | 126E-06 | 495E-05 | -042888065 | 0 |
| Bcat1 | ENSMUSG00000030268 | 3356883266 | 2031072656 | 5362370997 | 558E-05 | 0000292019 | 3234686714 | 2778008139 | 5619439847 | 449E-05 | 00003887 | -0122196552 | 0 |
| Scn1b | ENSMUSG00000019194 | 3374984567 | 1405037977 | 1582701316 | 764E-07 | 146E-05 | 3315982245 | 1469575728 | 9843388387 | 497E-06 | 0000101592 | -0059002322 | 0 |
| Mybl1 | ENSMUSG00000025912 | 3382500758 | 208009405 | 1515847738 | 912E-07 | 161E-05 | 3795055973 | 3020968743 | 5715474164 | 421E-05 | 0000372381 | 0412555215 | 1 |
| Dkkl1 | ENSMUSG00000030792 | 3387120495 | 21222399 | 8270017844 | 105E-05 | 820E-05 | 269235307 | 2304316247 | 433939899 | 000011839 | 0000739625 | -0694767425 | 0 |
| Hif3a | ENSMUSG00000004328 | 3388096453 | 4399267931 | 119185426 | 243E-06 | 311E-05 | 2607736813 | 4279885948 | 1004181115 | 458E-06 | 970E-05 | -078035964 | 0 |
| Gm35035 | ENSMUSG00000112082 | 3394943389 | 1061267942 | 6716615124 | 236E-05 | 0000150284 | 2932320492 | 0879575366 | 5748842548 | 411E-05 | 0000367746 | -0462622897 | 0 |
| Mdfic | ENSMUSG00000041390 | 3413310229 | 4753735763 | 4113785076 | 141E-08 | 142E-06 | 2846653017 | 489271907 | 1064454 | 362E-06 | 845E-05 | -0566657212 | 0 |
| Slc35f5 | ENSMUSG00000026342 | 3423194735 | 205910289 | 1981069977 | 302E-07 | 852E-06 | 2293986532 | 2258660694 | 4146276474 | 0000139934 | 0000823167 | -1129208203 | 0 |
| Syngr3 | ENSMUSG00000007021 | 3425845984 | 088235188 | 5882843044 | 392E-05 | 0000221884 | 4241720369 | 082672009 | 4323168764 | 0000120036 | 0000745805 | 0815874385 | 1 |
| Tpi1 | ENSMUSG00000023456 | 3431213192 | 801177499 | 1072355149 | 372E-06 | 412E-05 | 3140142873 | 8122147785 | 883096849 | 767E-06 | 0000132861 | -0291070319 | 0 |

| | | | | | | | | | | | | | |
|---|---|---|---|---|---|---|---|---|---|---|---|---|---|
| Ahnak | ENSMUSG00000069833 | 3435521943 | 1020433862 | 6430394309 | 211E-09 | 561E-07 | 2644788867 | 1002244717 | 1538879488 | 801E-07 | 390E-05 | -0790733076 | 0 |
| Fam83h | ENSMUSG00000046761 | 3436697123 | 191915043 | 1960619368 | 315E-07 | 879E-06 | 3574607304 | 1899054751 | 9598754403 | 549E-06 | 0000105943 | 0137910181 | 1 |
| Cdc25b | ENSMUSG00000027330 | 3439929796 | 8361017636 | 6635420704 | 185E-09 | 502E-07 | 2983026441 | 8239060446 | 1935448559 | 309E-07 | 247E-05 | -0456903355 | 0 |
| Muc3a | ENSMUSG00000094840 | 3450196213 | 2596429964 | 159228257 | 745E-07 | 144E-05 | 3383675994 | 2440062795 | 1323796013 | 149E-06 | 540E-05 | -0066520219 | 0 |
| Cxcr6 | ENSMUSG00000048521 | 3484719924 | 5090794782 | 1325128908 | 158E-06 | 238E-05 | 2720943819 | 5106902915 | 4073631835 | 0000149272 | 0000858874 | -0763776105 | 0 |
| Lgals1 | ENSMUSG00000068220 | 3494450627 | 8046744154 | 5160933116 | 539E-09 | 862E-07 | 3448614641 | 842898618 | 1465144179 | 981E-07 | 430E-05 | -0045835986 | 0 |
| Scd2 | ENSMUSG00000025203 | 3503117897 | 8906186756 | 2578429626 | 101E-07 | 413E-06 | 300034171 | 877889889 | 1210779436 | 215E-06 | 651E-05 | -0502776187 | 0 |
| Bmpr1a | ENSMUSG00000021796 | 3518509356 | 3116848645 | 2298258695 | 163E-07 | 569E-06 | 3455582065 | 3063347747 | 3132847912 | 407E-08 | 863E-06 | -0062927291 | 0 |
| Cst7 | ENSMUSG00000068129 | 3539050719 | 5914380722 | 4237801976 | 124E-08 | 131E-06 | 3019945363 | 5836492945 | 2467586925 | 112E-07 | 154E-05 | -0519105356 | 0 |
| Kcnk5 | ENSMUSG00000023243 | 3541125346 | 3929867497 | 128062304 | 182E-06 | 261E-05 | 3006085985 | 4143873042 | 5622725029 | 448E-05 | 0000388533 | -0535039361 | 0 |
| Hmcn2 | ENSMUSG00000055632 | 3543450017 | 290537785 | 2899103752 | 616E-08 | 320E-06 | 3500089887 | 3129229604 | 1239745161 | 195E-06 | 625E-05 | -004336013 | 0 |
| Gpr68 | ENSMUSG00000047415 | 3554955412 | 639957541 | 4816519874 | 723E-09 | 105E-06 | 2854936301 | 6348126372 | 240615418 | 124E-07 | 158E-05 | -0700019111 | 0 |
| Lockd | ENSMUSG00000098318 | 3567016941 | 1864805192 | 1091947169 | 346E-06 | 393E-05 | 3535597527 | 2532963742 | 4809666083 | 808E-05 | 000057044 | -0031419414 | 0 |
| Gpat3 | ENSMUSG00000029314 | 3589355937 | 1460778419 | 6623508798 | 249E-05 | 0000155888 | 3329379483 | 1269994585 | 7588477481 | 140E-05 | 0000190725 | -0259976454 | 0 |
| Zdhhc2 | ENSMUSG00000039470 | 3593391995 | 3153664851 | 1002278778 | 488E-06 | 495E-05 | 3078730324 | 375989893 | 4992978504 | 702E-05 | 0000518113 | -0514661671 | 0 |
| Cdc20 | ENSMUSG00000006398 | 3598011969 | 5427144989 | 2411055383 | 133E-07 | 505E-06 | 2903729688 | 5973043415 | 3951014573 | 0000166841 | 0000926989 | -0694282281 | 0 |
| Gm17745 | ENSMUSG00000110647 | 360249513 | 1787647349 | 8790573834 | 823E-06 | 699E-05 | 3052885515 | 1837820569 | 1282085488 | 170E-06 | 582E-05 | -0549609615 | 0 |
| Farp1 | ENSMUSG00000025555 | 3613795021 | 4183020537 | 6176730533 | 251E-09 | 603E-07 | 3146369317 | 4428008262 | 3636477909 | 216E-08 | 673E-06 | -0467425704 | 0 |
| Lag3 | ENSMUSG00000030124 | 3644873714 | 4974410904 | 5278916166 | 490E-09 | 857E-07 | 2945817487 | 5225841261 | 4782746142 | 825E-05 | 0000577144 | -0699056227 | 0 |
| Tktl1 | ENSMUSG00000031397 | 3664561681 | 1288578906 | 5486563629 | 511E-05 | 0000272025 | 3830936151 | 0964422022 | 5735903149 | 415E-05 | 0000369991 | 016637447 | 1 |
| Dcxr | ENSMUSG00000039450 | 3674986031 | 3038699511 | 1526486213 | 886E-07 | 159E-05 | 2497622472 | 3047279193 | 5523954461 | 479E-05 | 0000404685 | -1177363559 | 0 |
| Ccr1 | ENSMUSG00000025804 | 3679043827 | 2410625813 | 6555244445 | 259E-05 | 0000160524 | 2888009904 | 2554192752 | 4654559567 | 913E-05 | 0000622205 | -0791033923 | 0 |
| Atg9b | ENSMUSG00000038295 | 3680540924 | 3832567221 | 728152819 | 124E-09 | 440E-07 | 3521972914 | 3484432262 | 2221620773 | 174E-07 | 183E-05 | -015856801 | 0 |
| Il12rb2 | ENSMUSG00000018341 | 3686586291 | 402184924 | 1653814054 | 637E-07 | 131E-05 | 3588700497 | 4616062917 | 1304456616 | 158E-06 | 560E-05 | -0097885794 | 0 |
| Map6 | ENSMUSG00000055407 | 3694465246 | 3195886962 | 2138049901 | 220E-07 | 683E-06 | 4188642069 | 3441611358 | 9868220929 | 492E-06 | 0000101391 | 0494176823 | 1 |
| Prr36 | ENSMUSG00000064125 | 3705933664 | 1359457137 | 4803667802 | 842E-05 | 0000399079 | 3303117165 | 1326888694 | 1433417726 | 107E-06 | 447E-05 | -0402816499 | 0 |
| Ctla2a | ENSMUSG00000044258 | 3736922643 | 4314879446 | 161288468 | 707E-07 | 139E-05 | 2769118972 | 3944422227 | 2789046698 | 666E-08 | 115E-05 | -0967803671 | 0 |
| Pgk1 | ENSMUSG00000062070 | 3743302255 | 951925952 | 6167688295 | 253E-09 | 603E-07 | 3190366972 | 8980918516 | 9279038346 | 629E-06 | 0000116724 | -0552935283 | 0 |
| Ppbp | ENSMUSG00000029372 | 3747210153 | 2609468482 | 3603084684 | 00002396 | 0000887022 | 4962891623 | 3141240231 | 1323087881 | 149E-06 | 540E-05 | 121568147 | 1 |
| Mlkl | ENSMUSG00000012519 | 3753468829 | 2881084779 | 2180273322 | 203E-07 | 648E-06 | 2997478839 | 3602509631 | 4602215106 | 952E-05 | 0000639597 | -075598999 | 0 |
| Serpine2 | ENSMUSG00000026249 | 3754186997 | 2454255945 | 2768107599 | 748E-08 | 354E-06 | 5272706931 | 4071052066 | 6327481035 | 284E-05 | 0000291183 | 1518519934 | 1 |
| Slc41a2 | ENSMUSG00000034591 | 3769580087 | 1954017653 | 9894204981 | 514E-06 | 510E-05 | 2018147465 | 1697714752 | 5632770969 | 445E-05 | 000038728 | -1751432622 | 0 |
| Rab26 | ENSMUSG00000079657 | 378426426 | 0764106557 | 1022193286 | 451E-06 | 469E-05 | 3769933766 | 0691440609 | 15523637 | 773E-07 | 383E-05 | -0014330494 | 0 |
| Gzma | ENSMUSG00000023132 | 3799555554 | 7612034637 | 3397124732 | 316E-08 | 223E-06 | 2396913925 | 7472873509 | 4481194627 | 0000105131 | 0000682395 | -1402641629 | 0 |
| B930095G15Rik | ENSMUSG00000085133 | 3818431291 | 231396026 | 8779508416 | 827E-06 | 701E-05 | 3615230938 | 265686341 | 1623039851 | 643E-07 | 351E-05 | -0203200353 | 0 |
| Nebl | ENSMUSG00000053702 | 3820034706 | 5090771484 | 2100506282 | 237E-07 | 716E-06 | 2616582811 | 4917039245 | 8259854197 | 100E-05 | 0000156616 | -1203451895 | 0 |
| Poglut3 | ENSMUSG00000034487 | 3827672732 | 2750206355 | 1571560827 | 786E-07 | 148E-05 | 2845750151 | 2730341316 | 4999195904 | 699E-05 | 0000516919 | -0981922581 | 0 |
| Ptpn13 | ENSMUSG00000034573 | 3830006692 | 4695895648 | 2054132601 | 260E-07 | 760E-06 | 2662560702 | 4409042994 | 6663979568 | 233E-05 | 0000260389 | -116744599 | 0 |
| Cyb561 | ENSMUSG00000019590 | 38358457 | 1063291857 | 5485230715 | 512E-05 | 0000272025 | 3492910545 | 1309906037 | 164740239 | 604E-07 | 343E-05 | -0342935155 | 0 |
| Plp2 | ENSMUSG00000031146 | 3839503176 | 6743720247 | 6042375897 | 276E-09 | 619E-07 | 3311999381 | 6654461858 | 2333005137 | 141E-07 | 166E-05 | -0527503795 | 0 |
| Gzmc | ENSMUSG00000079186 | 384906371 | 2996871883 | 2130995723 | 223E-07 | 691E-06 | 4267992571 | 3473634565 | 3466935419 | 265E-08 | 728E-06 | 0418928861 | 1 |
| Ndrg4 | ENSMUSG00000036564 | 3882789128 | 1856484201 | 2239820707 | 181E-07 | 610E-06 | 3580187257 | 1642253038 | 138496083 | 124E-06 | 492E-05 | -0302601871 | 0 |
| Itga7 | ENSMUSG00000025348 | 3890798159 | 6939364475 | 2446927456 | 125E-07 | 483E-06 | 2881153593 | 6492196475 | 4480255033 | 0000105213 | 0000682422 | -1009644566 | 0 |
| Apbb1 | ENSMUSG00000037032 | 3892722199 | 3314984589 | 8435966294 | 968E-06 | 777E-05 | 3352371689 | 3162261916 | 1734232091 | 488E-07 | 325E-05 | -054035051 | 0 |
| Lurap1 | ENSMUSG00000028701 | 3906626092 | 0732447715 | 1517069441 | 909E-07 | 161E-05 | 3720463264 | 1301500631 | 8117259584 | 107E-05 | 0000163012 | -0186162828 | 0 |
| Adam8 | ENSMUSG00000025473 | 3919033433 | 504345939 | 970266215 | 555E-06 | 536E-05 | 3472631027 | 5430468589 | 1963319558 | 291E-07 | 239E-05 | -0446402406 | 0 |
| Anxa2 | ENSMUSG00000032231 | 3920506575 | 7775581446 | 1870779627 | 383E-07 | 974E-06 | 2899688022 | 7876989266 | 6060496916 | 336E-05 | 0000324455 | -1020818553 | 0 |
| Gm973 | ENSMUSG00000047361 | 3931894473 | 188892157 | 1432636944 | 115E-06 | 191E-05 | 3831911682 | 1911012539 | 7864687674 | 122E-05 | 0000174871 | -0099982791 | 0 |
| Cacnb3 | ENSMUSG00000003352 | 3952715879 | 5833432851 | 1187315868 | 154E-10 | 257E-07 | 3372519115 | 570861301 | 2573428864 | 935E-08 | 140E-05 | -0580196764 | 0 |
| Adgra3 | ENSMUSG00000029090 | 3960913222 | 4239093465 | 3501502681 | 278E-08 | 203E-06 | 3500079318 | 3877476355 | 7092836759 | 182E-05 | 0000224208 | -0460833904 | 0 |
| Maf | ENSMUSG00000055435 | 3969950251 | 7233990474 | 4070229603 | 147E-08 | 146E-06 | 2769488872 | 6686018859 | 6529450662 | 252E-05 | 0000271113 | -1200461379 | 0 |
| Plod2 | ENSMUSG00000032374 | 3981909034 | 3103305584 | 1142733612 | 288E-06 | 347E-05 | 3393591254 | 3149895947 | 136927027 | 130E-06 | 496E-05 | -058831778 | 0 |
| Hcn2 | ENSMUSG00000020331 | 4000105434 | 1298964332 | 8734278678 | 844E-06 | 706E-05 | 4554222825 | 1535885711 | 9655816572 | 536E-06 | 0000104804 | 0554117391 | 1 |
| Bnip3 | ENSMUSG00000078566 | 4003246287 | 4059321419 | 669338647 | 239E-05 | 0000151537 | 332457579 | 3702841104 | 9606919788 | 548E-06 | 000010575 | -0678670497 | 0 |
| Troap | ENSMUSG00000032783 | 40135506 | 2991378923 | 1800595464 | 449E-07 | 107E-05 | 318483667 | 3682905961 | 3966138417 | 0000164541 | 0000917151 | -082871393 | 0 |
| Rnf128 | ENSMUSG00000031438 | 4015366559 | 1906208817 | 1164674633 | 267E-06 | 329E-05 | 3338855989 | 220894411 | 4122067606 | 0000142964 | 0000836474 | -067651057 | 0 |
| Rasgrp4 | ENSMUSG00000030589 | 4026392187 | 5353026847 | 6731957615 | 174E-09 | 483E-07 | 2716673632 | 5134260287 | 4443683732 | 0000108453 | 0000697098 | -1309718555 | 0 |
| Gna15 | ENSMUSG00000034792 | 4031318203 | 4828300048 | 1849708857 | 402E-07 | 996E-06 | 3021199659 | 4992823067 | 8299555208 | 982E-06 | 00001553 | -1010118544 | 0 |
| Car5b | ENSMUSG00000031373 | 4041723459 | 2306294881 | 2695586373 | 836E-08 | 382E-06 | 3162377715 | 2734014607 | 8119342583 | 107E-05 | 0000163012 | -0879345744 | 0 |
| Tnip3 | ENSMUSG00000044162 | 4079575543 | 142884392 | 4834733909 | 822E-05 | 0000392222 | 3631514669 | 1681789131 | 7585281758 | 140E-05 | 0000190828 | -0448060874 | 0 |
| Unc13a | ENSMUSG00000034799 | 4118886782 | 5435967318 | 2224489511 | 187E-07 | 616E-06 | 3791002215 | 5377612262 | 9850029341 | 495E-06 | 0000101592 | -0327884567 | 0 |
| Mir155hg | ENSMUSG00000097418 | 4119865401 | 1896819144 | 1707820991 | 558E-07 | 120E-05 | 4841228964 | 2417960957 | 1913904704 | 324E-07 | 252E-05 | 0721363563 | 1 |
| Ppp2r3a | ENSMUSG00000043154 | 4126189027 | 4610168056 | 6700618584 | 238E-05 | 0000151309 | 3630339997 | 4439504316 | 9188337029 | 655E-06 | 0000119041 | -049584903 | 0 |
| Adap1 | ENSMUSG00000056413 | 4145590173 | 5621014132 | 6059539848 | 272E-09 | 619E-07 | 2914386565 | 5630551966 | 7107962056 | 181E-05 | 0000222798 | -1231203608 | 0 |
| Gm19391 | ENSMUSG00000105139 | 4167324782 | 0595734606 | 7461916323 | 157E-05 | 0000110649 | 4119493495 | 0908691366 | 1213844672 | 212E-06 | 651E-05 | -0047831287 | 0 |
| Il12rb1 | ENSMUSG00000000791 | 4181738866 | 6013574557 | 336317708 | 330E-08 | 227E-06 | 3467588747 | 6020601794 | 1989824364 | 275E-07 | 232E-05 | -0714150119 | 0 |
| Gucy2e | ENSMUSG00000020890 | 419712166 | 1453088113 | 1787777677 | 462E-07 | 109E-05 | 466015279 | 2408230117 | 1199298615 | 223E-06 | 652E-05 | 046303113 | 1 |
| Ctla4 | ENSMUSG00000026011 | 4211570075 | 7518549534 | 1060310691 | 250E-10 | 265E-07 | 3457968417 | 7034835593 | 1129286226 | 285E-06 | 743E-05 | -0753601658 | 0 |
| Cep55 | ENSMUSG00000024989 | 4225955042 | 423277631 | 154242881 | 849E-07 | 155E-05 | 3631508132 | 4832849086 | 50675611 | 664E-05 | 0000500971 | -059444691 | 0 |
| Srxn1 | ENSMUSG00000032802 | 4227781604 | 197459014 | 1291012562 | 176E-06 | 257E-05 | 3405966552 | 2346705986 | 1097938841 | 319E-06 | 783E-05 | -0821815052 | 0 |
| Prelid2 | ENSMUSG00000056671 | 4252517622 | 2826553953 | 1184144355 | 249E-06 | 317E-05 | 273565407 | 2588397307 | 4641958806 | 922E-05 | 0000627105 | -1516863552 | 0 |
| Tnfsf4 | ENSMUSG00000026700 | 4259022672 | 1969260963 | 2324522003 | 155E-07 | 549E-06 | 4466906533 | 2388381715 | 4104178904 | 0000145255 | 0000843369 | 0207883861 | 1 |
| Atp2b4 | ENSMUSG00000026463 | 4265832129 | 6786862005 | 2792640933 | 721E-08 | 352E-06 | 3038792901 | 6770571633 | 7560895058 | 142E-05 | 000019197 | -1227039228 | 0 |
| Tspan17 | ENSMUSG00000025875 | 4286009648 | 2175046662 | 2448828313 | 125E-07 | 483E-06 | 2898894342 | 1975166709 | 4450532043 | 0000107837 | 0000693875 | -1387115306 | 0 |
| Lgals3 | ENSMUSG00000050335 | 4288044125 | 5900033432 | 1331390722 | 155E-06 | 236E-05 | 2964172645 | 5923857781 | 5749749654 | 411E-05 | 0000367746 | -132387148 | 0 |
| Mylk | ENSMUSG00000022836 | 4307325108 | 1023829624 | 3951605623 | 0000172119 | 0000696188 | 4600525213 | 1697201036 | 1344195574 | 140E-06 | 519E-05 | 0293200105 | 1 |
| D630039A03Rik | ENSMUSG00000052117 | 4321885035 | 3642145734 | 6738846523 | 173E-09 | 483E-07 | 4517817355 | 3796677717 | 8525693275 | 563E-10 | 229E-06 | 019593232 | 1 |
| Lilr4b | ENSMUSG00000112023 | 4338024722 | 5955842639 | 1854012992 | 398E-07 | 992E-06 | 3269744614 | 6081050634 | 137058574 | 129E-06 | 496E-05 | -1068280108 | 0 |
| Adarb1 | ENSMUSG00000020262 | 4344378569 | 4492561551 | 7648421107 | 101E-09 | 380E-07 | 3868237458 | 421055784 | 4617406278 | 782E-09 | 472E-06 | -0476141111 | 0 |
| Gstt3 | ENSMUSG00000001665 | 4368260972 | 0942230165 | 4867064944 | 802E-05 | 0000386279 | 3920286882 | 1484032806 | 1201517806 | 221E-06 | 652E-05 | -044797409 | 0 |
| Ajuba | ENSMUSG00000022178 | 4376349558 | 27622485 | 176906425 | 483E-07 | 112E-05 | 4344825692 | 32412829 | 3110287717 | 420E-08 | 863E-06 | -0031523866 | 0 |
| Tbc1d2 | ENSMUSG00000039813 | 4383770603 | 3716513003 | 4394946012 | 107E-08 | 121E-06 | 3583459911 | 3584911485 | 7639912154 | 136E-05 | 0000188198 | -0800310692 | 0 |
| Fam71b | ENSMUSG00000020401 | 4389189327 | 4088765452 | 4875735314 | 686E-09 | 101E-06 | 3752802596 | 3789610429 | 6842371617 | 145E-09 | 229E-06 | -0636386731 | 0 |
| C030034L19Rik | ENSMUSG00000097365 | 4413673751 | 2830146599 | 1088516135 | 350E-06 | 396E-05 | 3797716707 | 3138730321 | 5539334445 | 474E-05 | 0000400981 | -0615957044 | 0 |
| Chst2 | ENSMUSG00000033350 | 4416286574 | 4516891024 | 2841939744 | 670E-08 | 339E-06 | 3203612841 | 4317603252 | 5630041137 | 446E-05 | 0000387719 | -1212673733 | 0 |
| Nrbp2 | ENSMUSG00000075590 | 4434627453 | 2707198979 | 1073413379 | 371E-06 | 411E-05 | 3034615762 | 2489630602 | 5348861955 | 542E-05 | 0000436696 | -1400011691 | 0 |
| Batf3 | ENSMUSG00000026630 | 4459974295 | 2503014135 | 4124994836 | 0000147326 | 0000616844 | 3028566662 | 2721801939 | 394660464 | 0000167519 | 0000929903 | -1431407633 | 0 |
| Nfil3 | ENSMUSG00000056749 | 4468711157 | 5919197077 | 1129361144 | 191E-10 | 260E-07 | 3988520929 | 5856052201 | 3289601111 | 331E-08 | 772E-06 | -0480190228 | 0 |
| 6430571L13Rik | ENSMUSG00000037977 | 4479147445 | 2035644736 | 3380314395 | 323E-08 | 224E-06 | 4534406347 | 2059872535 | 1116613385 | 298E-06 | 758E-05 | 0055258902 | 1 |
| Tg | ENSMUSG00000053469 | 4486596009 | 3253069178 | 2939986237 | 581E-08 | 311E-06 | 3540763377 | 3035292635 | 4291535213 | 0000123325 | 0000756574 | -0945832632 | 0 |
| Tnfsf13b | ENSMUSG00000031497 | 4493926641 | 1649112638 | 1844443377 | 406E-07 | 100E-05 | 330501317 | 1541474104 | 6264912514 | 296E-05 | 0000297578 | -1188913471 | 0 |
| Lilrb4a | ENSMUSG00000112148 | 452161456 | 4495032438 | 2735072943 | 787E-08 | 366E-06 | 2889554164 | 4657485617 | 7568983087 | 141E-05 | 0000191793 | -1632060396 | 0 |
| Sapcd2 | ENSMUSG00000026955 | 4532932607 | 3034548127 | 2577779915 | 101E-07 | 413E-06 | 3113468481 | 3426609502 | 3862403885 | 000018114 | 0000983899 | -1419464126 | 0 |
| Ccr4 | ENSMUSG00000047898 | 4537141426 | 4999930056 | 1951696376 | 321E-07 | 884E-06 | 3769747012 | 5121810918 | 9807523723 | 504E-06 | 0000102153 | -0767394414 | 0 |
| Hk2 | ENSMUSG00000000628 | 4549050275 | 5312184091 | 352509667 | 271E-08 | 200E-06 | 3416755986 | 5452843655 | 3919893953 | 0000171699 | 0000945319 | -1132294289 | 0 |
| Filip1 | ENSMUSG00000034898 | 4561843101 | 0951441237 | 7890124339 | 126E-05 | 941E-05 | 4390326275 | 1552033217 | 6600387851 | 242E-05 | 000026513 | -0171516826 | 0 |
| Tigit | ENSMUSG00000071552 | 4575490066 | 4861246168 | 17290832 | 531E-07 | 117E-05 | 295994616 | 4559687801 | 1024884161 | 422E-06 | 925E-05 | -1615543906 | 0 |
| B3galt2 | ENSMUSG00000033849 | 4577864049 | 1615986909 | 1647576746 | 647E-07 | 131E-05 | 4696612249 | 140207497 | 7636083787 | 137E-05 | 0000188198 | 01187482 | 1 |
| Lif | ENSMUSG00000034394 | 4584251683 | 4018245513 | 3519390683 | 273E-08 | 200E-06 | 5473004831 | 4750086839 | 8556897074 | 870E-06 | 0000145116 | 0888753148 | 1 |
| Zbtb32 | ENSMUSG00000006310 | 4614975931 | 4418446354 | 2353347756 | 148E-07 | 538E-06 | 3659171289 | 4699276079 | 9635517784 | 541E-06 | 0000105107 | -0955804642 | 0 |
| Nrn1 | ENSMUSG00000039114 | 4634114413 | 3629947299 | 1794744263 | 455E-07 | 108E-05 | 3087328091 | 2495698472 | 457350546 | 975E-05 | 0000648892 | -1546786322 | 0 |
| Cenpf | ENSMUSG00000026605 | 4645330221 | 5888817105 | 2179797489 | 203E-07 | 648E-06 | 3813073965 | 6513262772 | 4016655852 | 0000157137 | 0000889368 | -0832256256 | 0 |
| Aifm2 | ENSMUSG00000020085 | 4658706935 | 2816652109 | 3217269196 | 398E-08 | 249E-06 | 3225682287 | 2841646844 | 9510534488 | 570E-06 | 0000108905 | -1433024648 | 0 |
| Ccr5 | ENSMUSG00000079227 | 4677325565 | 5929208681 | 1553808005 | 824E-07 | 153E-05 | 3042449572 | 5506964099 | 4756343101 | 842E-05 | 00005845 | -1634875993 | 0 |
| Prnp | ENSMUSG00000079037 | 4686420775 | 5115628761 | 6248105675 | 239E-09 | 603E-07 | 2904445021 | 3639406691 | 6412921399 | 270E-05 | 0000282182 | -1781975754 | 0 |
| Ptger2 | ENSMUSG00000037759 | 468673162 | 533576001 | 8826775609 | 547E-10 | 347E-07 | 4383059089 | 5085117424 | 1440217169 | 105E-06 | 444E-05 | -0303672531 | 0 |
| Celf5 | ENSMUSG00000034818 | 4711915008 | 1307096699 | 8281751369 | 104E-05 | 817E-05 | 3864232148 | 1543726615 | 8257020066 | 100E-05 | 0000156616 | -084768286 | 0 |
| Tmem159 | ENSMUSG00000030917 | 4762539782 | 2574245124 | 4003236444 | 158E-08 | 150E-06 | 3944574094 | 2454885901 | 1213560374 | 213E-06 | 651E-05 | -0817965688 | 0 |
| Ctsf | ENSMUSG00000083282 | 4780436124 | 3790234535 | 4509640382 | 955E-09 | 114E-06 | 4288103853 | 2236460256 | 1431678601 | 108E-06 | 447E-05 | -0492332271 | 0 |
| Cd80 | ENSMUSG00000075122 | 4785874422 | 1914338862 | 1567392849 | 795E-07 | 149E-05 | 3497166321 | 2644242986 | 5261401775 | 576E-05 | 0000455684 | -1288708101 | 0 |
| Dach2 | ENSMUSG00000025592 | 4788916423 | 1595842935 | 2477082791 | 119E-07 | 471E-06 | 4110411967 | 171079524 | 7680426239 | 133E-05 | 0000185176 | -0678504456 | 0 |

| | | | | | | | | | | | | | |
|---|---|---|---|---|---|---|---|---|---|---|---|---|---|
| Bcam | ENSMUSG00000002980 | 4811638811 | 3768928906 | 3924852942 | 172E-08 | 157E-06 | 415350523 | 3512444527 | 1305912988 | 157E-06 | 560E-05 | -0658133581 | 0 |
| Perp | ENSMUSG00000019851 | 4842351661 | 1580176595 | 2302856959 | 162E-07 | 566E-06 | 4081049902 | 1689737007 | 7253525657 | 167E-05 | 0000213368 | -0761301759 | 0 |
| Gm16712 | ENSMUSG00000097077 | 4899801108 | 1264570464 | 2188196298 | 200E-07 | 641E-06 | 3378652062 | 1227534818 | 444121198 | 0000108676 | 0000697291 | -1521149046 | 0 |
| Slc16a3 | ENSMUSG00000025161 | 4936696402 | 574397076 | 1454760397 | 108E-06 | 183E-05 | 4480348804 | 5851939703 | 1271765778 | 176E-06 | 594E-05 | -0456347598 | 0 |
| Ppfia3 | ENSMUSG00000003863 | 4975680063 | 0470215557 | 9923423608 | 508E-06 | 506E-05 | 4514643268 | 0310525057 | 6594800068 | 242E-05 | 0000265523 | -0461036795 | 0 |
| Gm47705 | ENSMUSG00000114033 | 498456552 | 1029916561 | 1262487148 | 192E-06 | 272E-05 | 3065965086 | 111116443 | 1100595225 | 316E-06 | 783E-05 | -1918600434 | 0 |
| Vegfa | ENSMUSG00000023951 | 4994289052 | 5346511988 | 4236195875 | 125E-08 | 131E-06 | 4609678979 | 50357296 | 8652432406 | 832E-06 | 0000140973 | -0384610073 | 0 |
| Ebi3 | ENSMUSG00000003206 | 4997400791 | 2924480358 | 1053396319 | 400E-06 | 434E-05 | 3842131374 | 2826741601 | 6591720178 | 243E-05 | 0000265766 | -1155269417 | 0 |
| Slco4a1 | ENSMUSG00000038963 | 501394024 | 5318605467 | 5256880297 | 498E-09 | 857E-07 | 4723090909 | 5354506847 | 6390139381 | 194E-09 | 229E-06 | -0290849331 | 0 |
| Arfgef3 | ENSMUSG00000019852 | 5015465553 | 2338366262 | 3121154281 | 452E-08 | 267E-06 | 3888449757 | 1797775248 | 5795596133 | 399E-05 | 0000361192 | -1127015796 | 0 |
| Eml1 | ENSMUSG00000058070 | 5028136421 | 020249578 | 8893216463 | 786E-06 | 678E-05 | 5032150219 | 0089886871 | 7732660784 | 130E-05 | 0000181994 | 0004013798 | 1 |
| Sema6b | ENSMUSG00000001227 | 5042495679 | 1106650008 | 2176313336 | 204E-07 | 650E-06 | 3180316602 | 1489430348 | 5891971183 | 374E-05 | 0000349013 | -1862179077 | 0 |
| Cfap54 | ENSMUSG00000020014 | 5081934447 | 1797989378 | 157619784 | 777E-07 | 147E-05 | 3919616638 | 1860108239 | 1006526522 | 454E-06 | 962E-05 | -1162317809 | 0 |
| Gm37881 | ENSMUSG00000104101 | 5093012669 | 0842164353 | 8276206241 | 104E-05 | 818E-05 | 5672807115 | 1044149082 | 6959096304 | 197E-05 | 0000235866 | 0579794446 | 1 |
| Il10 | ENSMUSG00000016529 | 5129468094 | 2237275255 | 3640117995 | 236E-08 | 189E-06 | 3324636224 | 1544993279 | 5096192752 | 650E-05 | 0000493827 | -180483187 | 0 |
| Aipl1 | ENSMUSG00000040554 | 5131521157 | 0552038517 | 12476617 | 202E-06 | 279E-05 | 3970176086 | 092390156 | 4885502543 | 762E-05 | 00005482 | -1161345071 | 0 |
| Gzmb | ENSMUSG00000015437 | 5166170201 | 7627303304 | 2777208273 | 738E-08 | 352E-06 | 5171086151 | 8326991828 | 418190134 | 119E-08 | 573E-06 | 000491595 | 1 |
| Tex15 | ENSMUSG00000009628 | 5170911999 | 2604963982 | 3166312187 | 425E-08 | 257E-06 | 4641164258 | 2839035269 | 6768726386 | 219E-05 | 0000251304 | -0529747741 | 0 |
| Ak4 | ENSMUSG00000028527 | 5214197464 | 3011346207 | 1020087089 | 455E-06 | 471E-05 | 4326886087 | 2866869463 | 667105151 | 232E-05 | 0000260027 | -0887311377 | 0 |
| Mt1 | ENSMUSG00000031765 | 5274177957 | 4530911916 | 4560541456 | 911E-09 | 112E-06 | 3681144029 | 4765450652 | 5855874577 | 383E-05 | 0000353663 | -1593033928 | 0 |
| Lamc1 | ENSMUSG00000026478 | 5314294403 | 5162031336 | 5207966255 | 519E-09 | 862E-07 | 3653546821 | 5027282937 | 7741516211 | 129E-05 | 0000181594 | -1660747582 | 0 |
| Il1r2 | ENSMUSG00000026073 | 5314961864 | 2197494882 | 3257494606 | 378E-08 | 249E-06 | 2682348375 | 1260615227 | 3976987494 | 0000162916 | 0000911069 | -2632613489 | 0 |
| Thbs1 | ENSMUSG00000040152 | 5333556395 | 3972744089 | 1508920499 | 929E-07 | 163E-05 | 4333744342 | 4268985689 | 6724037023 | 225E-05 | 0000254976 | -0999812053 | 0 |
| 2900052L18Rik | ENSMUSG00000043993 | 5350317197 | 1653223834 | 7906670525 | 125E-05 | 937E-05 | 4506889696 | 1723949423 | 5682167078 | 430E-05 | 000037785 | -0843427501 | 0 |
| Il13 | ENSMUSG00000020383 | 5381014028 | 0929003069 | 775630342 | 135E-05 | 992E-05 | 6155055134 | 1140411076 | 176512174 | 454E-07 | 314E-05 | 0774041106 | 1 |
| Cetn4 | ENSMUSG00000045031 | 5383740151 | 165285542 | 2897331638 | 618E-08 | 320E-06 | 4977358066 | 160442116 | 8103734059 | 108E-05 | 0000163275 | -0406382085 | 0 |
| Gpm6b | ENSMUSG00000031342 | 541975244 | 5825151036 | 559816004 | 381E-09 | 727E-07 | 4172987015 | 5301420184 | 1192809144 | 228E-06 | 652E-05 | -1246765425 | 0 |
| S100a6 | ENSMUSG00000001025 | 5426950346 | 6603061509 | 2797907744 | 715E-08 | 352E-06 | 442206126 | 6750646084 | 134787266 | 138E-06 | 516E-05 | -1004889086 | 0 |
| Phlda3 | ENSMUSG00000041801 | 5508068641 | 4295661788 | 3757937772 | 207E-08 | 177E-06 | 5592765839 | 4447486145 | 1803772861 | 415E-07 | 291E-05 | 0084697198 | 1 |
| Pimreg | ENSMUSG00000020808 | 5531638181 | 4517174374 | 2687785144 | 847E-08 | 382E-06 | 4991082348 | 5019809928 | 982443926 | 500E-06 | 0000101836 | -0540555833 | 0 |
| Pf4 | ENSMUSG00000029373 | 5540432069 | 2560838522 | 9850806466 | 523E-06 | 517E-05 | 4502176842 | 2717092506 | 5287163438 | 566E-05 | 0000449451 | -1038255227 | 0 |
| Gpnmb | ENSMUSG00000029816 | 5541381649 | 5216734421 | 5068366843 | 582E-09 | 906E-07 | 467078994 | 4890958127 | 2572993074 | 936E-08 | 140E-05 | -0870591709 | 0 |
| Akr1c18 | ENSMUSG00000021214 | 5552387799 | 1185749415 | 7346044627 | 167E-05 | 0000115724 | 4656656224 | 077102802 | 5845491376 | 386E-05 | 00003552 | -0895731575 | 0 |
| Gm37145 | ENSMUSG00000102278 | 5559210726 | 2885795429 | 258991773 | 989E-08 | 411E-06 | 4984156014 | 277170525 | 1603216374 | 676E-07 | 353E-05 | -0575054712 | 0 |
| Lamc2 | ENSMUSG00000026479 | 5589758277 | 0474047216 | 9684283965 | 560E-06 | 537E-05 | 4797294208 | 0207754247 | 4296016254 | 0000122852 | 0000754843 | -0792464069 | 0 |
| Socs2 | ENSMUSG00000020027 | 5638315316 | 5015895025 | 8233726851 | 737E-10 | 347E-07 | 4144842125 | 4524555577 | 1044540475 | 391E-06 | 888E-05 | -1493473191 | 0 |
| Pros1 | ENSMUSG00000022912 | 5724629917 | 5249623321 | 5807671234 | 326E-09 | 692E-07 | 4895561968 | 5363499156 | 3003182845 | 487E-08 | 909E-06 | -0829067949 | 0 |
| Galr3 | ENSMUSG00000114755 | 575812861 | 1104810239 | 1484329403 | 994E-07 | 172E-05 | 5243848538 | 1111527564 | 5030637008 | 683E-05 | 0000510824 | -0514280072 | 0 |
| Cdkn1a | ENSMUSG00000023067 | 5960292599 | 5916355953 | 459028858 | 886E-09 | 111E-06 | 5650199469 | 6041708046 | 2108471139 | 216E-07 | 211E-05 | -031009313 | 0 |
| Pkp2 | ENSMUSG00000041957 | 5993604869 | 1423509438 | 5349327792 | 563E-05 | 0000294097 | 3702275375 | 0644701774 | 5961926039 | 358E-05 | 0000335944 | -2291329494 | 0 |
| Aldoc | ENSMUSG00000017390 | 6043259107 | 5959934782 | 2054231565 | 260E-07 | 760E-06 | 4988934551 | 5457178285 | 170134682 | 529E-07 | 334E-05 | -1054324556 | 0 |
| Il2ra | ENSMUSG00000026770 | 6057064766 | 7670490936 | 9069546621 | 487E-10 | 335E-07 | 6067839822 | 8003236693 | 1834866896 | 386E-07 | 279E-05 | 0010775056 | 1 |
| Dnm1 | ENSMUSG00000026825 | 6117390592 | 1721360631 | 8237486149 | 106E-05 | 830E-05 | 565497915 | 1371209353 | 6031378965 | 342E-05 | 0000326366 | -0462411442 | 0 |
| Mgarp | ENSMUSG00000037161 | 613576731 | 3288515808 | 1361646415 | 141E-06 | 222E-05 | 4609872214 | 248852487 | 659802833 | 242E-05 | 0000265259 | -1525895096 | 0 |
| 1500009L16Rik | ENSMUSG00000087651 | 6217884547 | 3957155336 | 1552260454 | 827E-07 | 153E-05 | 5641764779 | 4241783356 | 1649611478 | 601E-07 | 342E-05 | -0576119768 | 0 |
| Egln3 | ENSMUSG00000035105 | 6223345449 | 6601302759 | 8174915256 | 760E-10 | 347E-07 | 5417919274 | 6136189284 | 2085203878 | 226E-07 | 214E-05 | -0805426175 | 0 |
| Gm20139 | ENSMUSG00000111052 | 6289118184 | 2068176883 | 1170858029 | 261E-06 | 324E-05 | 5965275396 | 2260079427 | 1982662605 | 280E-07 | 232E-05 | -0323842788 | 0 |
| Scrt1 | ENSMUSG00000048385 | 6406603489 | 2061646742 | 5967265678 | 372E-05 | 0000212411 | 6120584246 | 0898144374 | 4500136647 | 0000103502 | 0000674923 | -0286019243 | 0 |
| Psrc1 | ENSMUSG00000068744 | 642019869 | 5847186085 | 304190818 | 271E-12 | 278E-08 | 5402668841 | 5868141024 | 8254945605 | 100E-05 | 0000156616 | -1017529849 | 0 |
| Serpinf1 | ENSMUSG00000000753 | 6426134436 | 4774163251 | 559896661 | 381E-09 | 727E-07 | 3968497439 | 3260207315 | 4607065348 | 949E-05 | 0000638426 | -2457636997 | 0 |
| Inhba | ENSMUSG00000041324 | 6465587195 | 276026447 | 2434772009 | 128E-07 | 490E-06 | 4384741895 | 2258240176 | 6462055604 | 262E-05 | 0000275423 | -20808453 | 0 |
| Rnf152 | ENSMUSG00000047496 | 6622663081 | 1444010967 | 1653535784 | 638E-07 | 131E-05 | 6126466297 | 1288415412 | 5604671722 | 453E-05 | 0000391091 | -0496196784 | 0 |
| Cyp11a1 | ENSMUSG00000032323 | 6645292782 | 6177982361 | 1109205519 | 206E-10 | 260E-07 | 554052073 | 6232181767 | 1069889697 | 355E-06 | 834E-05 | -1104772052 | 0 |
| Hc | ENSMUSG00000026874 | 6754225516 | 3754895252 | 2386331111 | 139E-07 | 516E-06 | 5682385799 | 3321250015 | 270394494 | 759E-08 | 124E-05 | -1071839717 | 0 |
| Spp1 | ENSMUSG00000029304 | 6836488218 | 1652300456 | 1316324022 | 162E-06 | 244E-05 | 4050396025 | 1854402727 | 1885938928 | 344E-07 | 258E-05 | -2786092193 | 0 |
| Trp53cor1 | ENSMUSG00000085912 | 6859545258 | 0747903715 | 1887684278 | 369E-07 | 955E-06 | 6055603785 | 1183907685 | 2024462963 | 256E-07 | 224E-05 | -0803941473 | 0 |
| Azin2 | ENSMUSG00000028789 | 6916865466 | 1698154802 | 1545949035 | 841E-07 | 155E-05 | 515962622 | 1987058723 | 1444960699 | 104E-06 | 442E-05 | -1757239246 | 0 |
| Esm1 | ENSMUSG00000042379 | 6979410796 | 1482323225 | 1251277589 | 199E-06 | 277E-05 | 3878391381 | 1052749984 | 3976497485 | 0000162989 | 0000911069 | -3101019415 | 0 |
| Gm17767 | ENSMUSG00000099413 | 6992936768 | 098199756 | 1750699037 | 504E-07 | 114E-05 | 5111514948 | 0983969346 | 618271552 | 311E-05 | 0000308283 | -188142182 | 0 |
| Gm17322 | ENSMUSG00000097604 | 6998129013 | 0735702758 | 1649968198 | 643E-07 | 131E-05 | 6441434963 | 1282692194 | 8237418031 | 101E-05 | 0000157075 | -055669405 | 0 |
| Ccl22 | ENSMUSG00000031779 | 7284440974 | 4560203111 | 5119450772 | 558E-09 | 880E-07 | 6086138922 | 55763073 | 1071802784 | 352E-06 | 831E-05 | -1198302052 | 0 |
| Stc2 | ENSMUSG00000020303 | 7472287914 | 4931843034 | 1390770836 | 782E-11 | 182E-07 | 6363549558 | 4368872077 | 1463960223 | 985E-07 | 430E-05 | -1108738356 | 0 |
| Obsl1 | ENSMUSG00000026211 | 7667149668 | 2410515933 | 1051327127 | 403E-06 | 436E-05 | 5198000987 | 2072935985 | 6873513964 | 206E-05 | 0000241817 | -2469148681 | 0 |
| Plcd1 | ENSMUSG00000010660 | 7695347397 | 3318119466 | 2554800153 | 105E-07 | 424E-06 | 4951799745 | 2332138283 | 1162400131 | 253E-06 | 705E-05 | -2743547652 | 0 |
| Penk | ENSMUSG00000045573 | 7713062655 | 4604803036 | 4205651581 | 128E-08 | 134E-06 | 5979973768 | 463368722 | 7725427791 | 130E-05 | 0000182457 | -1733088887 | 0 |
| Gm37795 | ENSMUSG00000102813 | 7767864556 | 3331182614 | 3293911508 | 360E-08 | 239E-06 | 7475939516 | 3740664709 | 228227837 | 155E-07 | 169E-05 | -029192504 | 0 |
| Ermn | ENSMUSG00000026830 | 8249209114 | 13585667 | 2683968564 | 852E-08 | 382E-06 | 7648018029 | 1013292859 | 1602034924 | 678E-07 | 353E-05 | -0601191085 | 0 |
| Gm5127 | ENSMUSG00000073010 | 8508154065 | 148782903 | 2052122627 | 261E-07 | 761E-06 | 4931243929 | 0966632698 | 5626657032 | 447E-05 | 0000387775 | -3576910136 | 0 |
| Lama5 | ENSMUSG00000015647 | 8649871143 | 6312318611 | 5254987074 | 499E-09 | 857E-07 | 6377005948 | 5741081358 | 1171198046 | 246E-06 | 695E-05 | -2272865195 | 0 |
| Tnfrsf8 | ENSMUSG00000028602 | 8695606504 | 5866570356 | 5670578937 | 361E-09 | 727E-07 | 7449432317 | 6012243162 | 1122690809 | 292E-06 | 751E-05 | -1246174187 | 0 |
| Cysltr1 | ENSMUSG00000052821 | 9104972222 | 5841427943 | 9879377769 | 338E-10 | 268E-07 | 6694139586 | 5159266982 | 1625492902 | 639E-07 | 351E-05 | -2410832636 | 0 |
| Inha | ENSMUSG00000032968 | 9692052743 | 2328782768 | 3113291319 | 457E-08 | 267E-06 | 7411553787 | 1921427861 | 1279407631 | 171E-06 | 584E-05 | -2280498956 | 0 |
| Upp1 | ENSMUSG00000020407 | 9962565332 | 1717827932 | 8445229852 | 964E-06 | 775E-05 | 552692347 | 0742962228 | 7980679144 | 115E-05 | 0000169251 | -4435641862 | 0 |
| Olfr60 | ENSMUSG00000060112 | 9967819806 | 3339380194 | 5669994823 | 361E-09 | 727E-07 | 8695054491 | 3094592485 | 3875919815 | 165E-08 | 595E-06 | -1272765315 | 0 |
| Vldlr | ENSMUSG00000024924 | 1107709039 | 3195124984 | 8316452351 | 102E-05 | 809E-05 | 6141330168 | 1770308408 | 5031861535 | 682E-05 | 0000510671 | -4935760222 | 0 |
| Adamtsl3 | ENSMUSG00000070469 | 1136814664 | 4983691175 | 2676715759 | 861E-08 | 382E-06 | 1128006141 | 3975163371 | 9204055864 | 650E-06 | 0000118763 | -008808523 | 0 |
| Gm38563 | ENSMUSG00000116498 | 1140738677 | 0305827254 | 3264526619 | 698E-07 | 138E-05 | 7358597771 | 0430323235 | 8512856102 | 888E-06 | 0000147316 | -4048788999 | 0 |
| Slc17a6 | ENSMUSG00000030500 | 1166913588 | 2150492958 | 2341562106 | 151E-07 | 541E-06 | 808740969 | 1170538112 | 9658078644 | 536E-06 | 0000104804 | -358172619 | 0 |
| Tmprss11e | ENSMUSG00000054537 | 1203269425 | 0973098716 | 2094894956 | 294E-06 | 353E-05 | 7947866472 | 064983677 | 9856577083 | 494E-06 | 0000101529 | -4084827778 | 0 |
| Tpbg | ENSMUSG00000035274 | 1221087296 | 1141841834 | 3286348908 | 683E-07 | 136E-05 | 7124034331 | 0195971651 | 9658537916 | 536E-06 | 0000104804 | -5086838629 | 0 |
| Tph1 | ENSMUSG00000040046 | 1232651721 | 1260266624 | 3875505476 | 399E-07 | 993E-06 | 9477949567 | 1694165488 | 104176876 | 395E-06 | 892E-05 | -2848567643 | 0 |
| Gzmg | ENSMUSG00000040284 | 1308999834 | 2041058084 | 1694135093 | 581E-06 | 550E-05 | 1177447977 | 1706914246 | 3374527671 | 278E-06 | 734E-05 | -131551857 | 0 |
| Gzmd | ENSMUSG00000059256 | 1328626681 | 2242957616 | 2790345838 | 116E-06 | 192E-05 | 127738957 | 2703347792 | 8162108058 | 231E-07 | 214E-05 | -051237111 | 0 |

**TABLE S4**

| Gene | ENSMBL | logFC_WT_F | logCPM_WT_F | F_WT_F | PValue_WT_F | FDR_WT_F |
|---|---|---|---|---|---|---|
| Slc28a2b | ENSMUSG00000079071 | -5241431416 | 6753534905 | 7185889485 | 132E-09 | 449E-07 |
| A530021J07Rik | ENSMUSG00000053528 | -518816046 | 3240114898 | 3600950587 | 247E-08 | 189E-06 |
| Gm48369 | ENSMUSG00000113349 | -5030158071 | 0687116353 | 4766272283 | 866E-05 | 0000406773 |
| Drc1 | ENSMUSG00000073102 | -4547585221 | 1136044701 | 156193936 | 806E-07 | 150E-05 |
| Wnt5b | ENSMUSG00000030170 | -4528480052 | 0429791925 | 1127450522 | 304E-06 | 360E-05 |
| Gm49492 | ENSMUSG00000116180 | -4105192355 | 0550738305 | 1278302753 | 183E-06 | 262E-05 |
| Kcnc1 | ENSMUSG00000058975 | -3562132004 | 1246825184 | 7864175503 | 128E-05 | 950E-05 |
| Ddx43 | ENSMUSG00000070291 | -3530548914 | 0816857326 | 1226036067 | 217E-06 | 291E-05 |
| St6galnac3 | ENSMUSG00000052544 | -3227911203 | 086993593 | 9254316402 | 671E-06 | 609E-05 |
| Bambi-ps1 | ENSMUSG00000081219 | -3139781602 | 4385009917 | 1937949949 | 331E-07 | 901E-06 |
| Samd3 | ENSMUSG00000051354 | -3134113272 | 1917755941 | 1016238771 | 462E-06 | 475E-05 |
| E030030I06Rik | ENSMUSG00000097327 | -3099825006 | 1888368641 | 8724232438 | 848E-06 | 708E-05 |
| Calhm6 | ENSMUSG00000046031 | -3044880692 | 3007125861 | 2590928157 | 987E-08 | 411E-06 |
| Trav15-1-dv6-1 | ENSMUSG00000094016 | -2931159823 | 1196833022 | 129325364 | 174E-06 | 257E-05 |
| Cldn10 | ENSMUSG00000022132 | -287623747 | 1106788 | 8528380075 | 927E-06 | 753E-05 |
| Hsph1 | ENSMUSG00000029657 | -2853897603 | 7167763401 | 4629099529 | 965E-05 | 0000443351 |
| Cdk5r1 | ENSMUSG00000048895 | -2813264583 | 1727087484 | 1474708653 | 102E-06 | 176E-05 |
| Gm26581 | ENSMUSG00000097108 | -2790438347 | 1101953562 | 4110945321 | 0000149163 | 0000622973 |
| Gm11455 | ENSMUSG00000085538 | -2765554179 | 0399866561 | 4235583736 | 0000133794 | 0000572723 |
| Rapgef4os3 | ENSMUSG00000086231 | -2714008841 | 1942058682 | 8691740771 | 860E-06 | 716E-05 |
| Ifi206 | ENSMUSG00000037849 | -2703424198 | 6805451068 | 3488023529 | 283E-08 | 205E-06 |
| Kifc3 | ENSMUSG00000031788 | -2669438148 | 0929409194 | 5619896533 | 467E-05 | 0000253562 |
| Usp18 | ENSMUSG00000030107 | -2652778813 | 3879969808 | 1152650469 | 278E-06 | 339E-05 |
| Gm2682 | ENSMUSG00000115681 | -2643965294 | 6747940046 | 3606009783 | 246E-08 | 189E-06 |
| Gm44941 | ENSMUSG00000107509 | -263857715 | 113626257 | 7805152371 | 131E-05 | 974E-05 |
| Arl4d | ENSMUSG00000034936 | -2636153818 | 2453797177 | 1344483832 | 149E-06 | 231E-05 |
| Gm37968 | ENSMUSG00000102570 | -2635091891 | 1785120877 | 4217186615 | 0000135935 | 0000578705 |
| Traj5 | ENSMUSG00000076923 | -2615774051 | 1273347224 | 4492262142 | 0000107842 | 000048326 |
| Slc25a29 | ENSMUSG00000021265 | -259865648 | 2252639854 | 5682666776 | 448E-05 | 0000245404 |
| Gm18911 | ENSMUSG00000116438 | -2588975658 | 0873682127 | 7661581949 | 141E-05 | 0000103033 |
| Actn1 | ENSMUSG00000015143 | -2577154975 | 8666981497 | 3417305496 | 309E-08 | 220E-06 |
| Ift80 | ENSMUSG00000027778 | -2560450576 | 7040464482 | 4398819867 | 106E-08 | 121E-06 |
| Iigp1 | ENSMUSG00000054072 | -2550789597 | 4019589108 | 1759788413 | 493E-07 | 114E-05 |
| NA | ENSMUSG00000103126 | -2547761328 | 4269461528 | 2278838772 | 169E-07 | 583E-06 |
| Acoxl | ENSMUSG00000027380 | -2545767324 | 1549834343 | 105308723 | 400E-06 | 434E-05 |
| Qser1 | ENSMUSG00000074994 | -2508110072 | 4253634934 | 1351351966 | 146E-06 | 227E-05 |
| Heatr9 | ENSMUSG00000018925 | -2497907315 | 0783306456 | 4119490032 | 0000148042 | 00006194 |
| Ccl5 | ENSMUSG00000035042 | -2483018846 | 4409582319 | 8721620569 | 849E-06 | 709E-05 |
| Nav1 | ENSMUSG00000009418 | -2477898385 | 148029373 | 4774805813 | 861E-05 | 000040564 |
| Gm38118 | ENSMUSG00000103089 | -2452760009 | 2478163465 | 7509532022 | 153E-05 | 0000109055 |
| Gm10382 | ENSMUSG00000072612 | -242833812 | 1804453716 | 5707976864 | 440E-05 | 0000242417 |
| Cd7 | ENSMUSG00000025163 | -2409770349 | 4001114 | 3879956359 | 0000183854 | 0000730976 |
| Tnfaip3 | ENSMUSG00000019850 | -2404406393 | 8912141124 | 8241821185 | 106E-05 | 830E-05 |
| Nod1 | ENSMUSG00000038058 | -2403643962 | 5167219525 | 1867090925 | 386E-07 | 978E-06 |
| Pou6f1 | ENSMUSG00000009739 | -239673546 | 5640184239 | 273652079 | 785E-08 | 366E-06 |
| Gm13842 | ENSMUSG00000084992 | -2396479736 | 1983316709 | 5060037832 | 693E-05 | 000034634 |
| NA | ENSMUSG00000097158 | -2392681149 | 0946011391 | 5248218705 | 605E-05 | 0000309433 |
| Lypd6b | ENSMUSG00000026765 | -2374858958 | 4403158073 | 3845804989 | 187E-08 | 164E-06 |
| Mid1 | ENSMUSG00000035299 | -2371146822 | 1906240169 | 418505807 | 0000139777 | 0000591608 |
| Gbp10 | ENSMUSG00000105096 | -2367212397 | 221161203 | 901659055 | 744E-06 | 653E-05 |
| Traj11 | ENSMUSG00000076918 | -2365765247 | 1344427537 | 8830711109 | 808E-06 | 693E-05 |
| Gm7027 | ENSMUSG00000067121 | -2365690991 | 0901672531 | 9678910692 | 561E-06 | 537E-05 |
| Ifit1bl2 | ENSMUSG00000067297 | -2360885825 | 0508867287 | 5532006426 | 496E-05 | 0000266064 |
| Hid1 | ENSMUSG00000034586 | -2359241803 | 5566267004 | 2461043143 | 122E-07 | 478E-06 |

| | | | | | | |
|---|---|---|---|---|---|---|
| Cers6 | ENSMUSG00000027035 | -2324017657 | 4898687398 | 4321051215 | 114E-08 | 125E-06 |
| Nim1k | ENSMUSG00000095930 | -2318432149 | 1189282949 | 6809131514 | 224E-05 | 0000144909 |
| Ksr2 | ENSMUSG00000061578 | -2309965297 | 2856198893 | 8657384463 | 874E-06 | 724E-05 |
| Tob2 | ENSMUSG00000048546 | -2271957218 | 7148859922 | 1083019853 | 357E-06 | 401E-05 |
| BC065397 | ENSMUSG00000087368 | -2258331393 | 2348899677 | 1097387997 | 339E-06 | 390E-05 |
| Rbm3os | ENSMUSG00000055188 | -2253052556 | 2213025588 | 667873066 | 241E-05 | 0000152523 |
| Trav14-1 | ENSMUSG00000076840 | -2247857563 | 2483104536 | 7195304038 | 181E-05 | 0000122885 |
| Perm1 | ENSMUSG00000078486 | -2243462533 | 2817931418 | 5349109683 | 563E-05 | 0000294097 |
| Ifi214 | ENSMUSG00000070501 | -224049973 | 4941964123 | 2745123187 | 775E-08 | 365E-06 |
| Tmie | ENSMUSG00000049555 | -2240350938 | 4845357147 | 1355171792 | 144E-06 | 225E-05 |
| Art2b | ENSMUSG00000030651 | -2237786411 | 5499039925 | 1019477721 | 456E-06 | 471E-05 |
| Whrn | ENSMUSG00000039137 | -2227764392 | 3404870717 | 8491740293 | 943E-06 | 763E-05 |
| Gm47234 | ENSMUSG00000111030 | -2224644314 | 0997977059 | 3637157738 | 000023171 | 000086402 |
| Gm17494 | ENSMUSG00000057359 | -2224422059 | 1359388526 | 5312829473 | 577E-05 | 000029939 |
| Traj13 | ENSMUSG00000076916 | -2224262813 | 0960147173 | 7770431777 | 134E-05 | 987E-05 |
| Cdh24 | ENSMUSG00000059674 | -2215020826 | 5128204408 | 3622167081 | 241E-08 | 189E-06 |
| Sntb1 | ENSMUSG00000060429 | -221436529 | 5305307846 | 5812189513 | 325E-09 | 692E-07 |
| Ifitm10 | ENSMUSG00000045777 | -2212519841 | 1285451985 | 44917519 | 0000107887 | 0000483277 |
| Gm17092 | ENSMUSG00000091623 | -2205209903 | 1734954022 | 8996785236 | 750E-06 | 656E-05 |
| Adgrg3 | ENSMUSG00000060470 | -2188185358 | 3085741776 | 1214763244 | 225E-06 | 299E-05 |
| Hmgn1 | ENSMUSG00000040681 | -218229227 | 6272482772 | 1723215817 | 538E-07 | 118E-05 |
| Gm49553 | ENSMUSG00000116433 | -2178011236 | 4407490642 | 1530860453 | 876E-07 | 158E-05 |
| Mctp2 | ENSMUSG00000032776 | -2164942845 | 4897922482 | 73738936 | 164E-05 | 0000114442 |
| Arl5c | ENSMUSG00000038352 | -2164308398 | 589952891 | 4125524768 | 139E-08 | 142E-06 |
| Il18r1 | ENSMUSG00000026070 | -2161124439 | 5564819347 | 7308129363 | 170E-05 | 0000117668 |
| Car2 | ENSMUSG00000027562 | -213568968 | 3133218317 | 4916095051 | 772E-05 | 0000376967 |
| Vipr1 | ENSMUSG00000032528 | -2132300103 | 7565309583 | 2632820999 | 923E-08 | 393E-06 |
| Dzip1 | ENSMUSG00000042156 | -2129123422 | 5884816935 | 4464229902 | 997E-09 | 116E-06 |
| Hspa5 | ENSMUSG00000026864 | -2125154016 | 9315155164 | 7798534144 | 132E-05 | 976E-05 |
| Tgm4 | ENSMUSG00000025787 | -2125078647 | 0678256279 | 4643253277 | 955E-05 | 0000440455 |
| Gm14453 | ENSMUSG00000087442 | -2123355452 | 2379238476 | 4694545485 | 917E-05 | 0000426276 |
| Nr4a2 | ENSMUSG00000026826 | -2122583088 | 3328976338 | 4193644001 | 0000138738 | 000058913 |
| Kcnh3 | ENSMUSG00000037579 | -2118413096 | 2203150162 | 6914815425 | 211E-05 | 0000138353 |
| Jaml | ENSMUSG00000048534 | -2107563903 | 2816411157 | 4570580753 | 0000101187 | 0000459436 |
| Ier5 | ENSMUSG00000056708 | -2106187384 | 8178166639 | 1096283594 | 340E-06 | 390E-05 |
| Traj7 | ENSMUSG00000076921 | -2095194225 | 1909606415 | 5493141003 | 509E-05 | 000027121 |
| Gm17024 | ENSMUSG00000090582 | -207482868 | 2083337172 | 3880879593 | 0000183697 | 0000730841 |
| Lpar6 | ENSMUSG00000033446 | -2064906266 | 3926396761 | 2687132009 | 848E-08 | 382E-06 |
| Otud1 | ENSMUSG00000043415 | -2063935853 | 5751305521 | 3604281305 | 246E-08 | 189E-06 |
| Apol7e | ENSMUSG00000071716 | -2063028911 | 537680843 | 321419414 | 399E-08 | 249E-06 |
| 4930562C15Rik | ENSMUSG00000022518 | -205500487 | 0769796334 | 534132192 | 566E-05 | 0000295318 |
| Ets2 | ENSMUSG00000022895 | -2054862212 | 5384084978 | 4993173743 | 620E-09 | 952E-07 |
| Gprasp1 | ENSMUSG00000043384 | -2047205404 | 5758360614 | 1299221149 | 171E-06 | 254E-05 |
| Traj2 | ENSMUSG00000076926 | -2046014904 | 2161247631 | 5682567696 | 448E-05 | 0000245404 |
| Tspan9 | ENSMUSG00000030352 | -2045589261 | 4372094932 | 5010912696 | 719E-05 | 0000356612 |
| Eomes | ENSMUSG00000032446 | -2042520802 | 3461834342 | 165073047 | 642E-07 | 131E-05 |
| Csrnp1 | ENSMUSG00000032515 | -2041564662 | 6770217239 | 8221380276 | 107E-05 | 835E-05 |
| Guca1b | ENSMUSG00000023979 | -2040140283 | 159590333 | 3982074809 | 000016741 | 0000680663 |
| Traj8 | ENSMUSG00000076920 | -2034169043 | 1357949181 | 5164027306 | 642E-05 | 0000325807 |
| Traj1 | ENSMUSG00000076927 | -2033988704 | 2115092349 | 5913967379 | 385E-05 | 000021863 |
| Id3 | ENSMUSG00000007872 | -2030666876 | 5029311972 | 1494328651 | 967E-07 | 169E-05 |
| Trav7n-4 | ENSMUSG00000095736 | -2026690129 | 1073602366 | 3566828138 | 0000248363 | 000091309 |
| Siah2 | ENSMUSG00000036432 | -2025835432 | 5496059203 | 4915229744 | 773E-05 | 0000376967 |
| Zbtb10 | ENSMUSG00000069114 | -2005416898 | 3380995129 | 3980789521 | 0000167605 | 0000681193 |
| Snora31 | ENSMUSG00000065147 | -2004847671 | 2762830214 | 1273671581 | 186E-06 | 265E-05 |
| Gm15283 | ENSMUSG00000087700 | 2004505377 | 204962159 | 3815853401 | 0000195199 | 0000761545 |
| D730005E14Rik | ENSMUSG00000115902 | 200555469 | 1519748253 | 9036749477 | 737E-06 | 648E-05 |

| | | | | | | |
|---|---|---|---|---|---|---|
| Gng2 | ENSMUSG00000043004 | 2005844902 | 5919624607 | 287882879 | 635E-08 | 326E-06 |
| Rilpl1 | ENSMUSG00000029392 | 2007638159 | 2271697235 | 9340051068 | 647E-06 | 592E-05 |
| Gapdhs | ENSMUSG00000061099 | 201175998 | 1351750482 | 3775705564 | 0000202746 | 0000782096 |
| Arhgap18 | ENSMUSG00000039031 | 2013405814 | 5804291432 | 2742292364 | 778E-08 | 365E-06 |
| Smim24 | ENSMUSG00000078439 | 2013555239 | 2685269073 | 3650976196 | 0000228601 | 0000854983 |
| Gm2a | ENSMUSG00000000594 | 2014795193 | 8303623673 | 3944106089 | 168E-08 | 155E-06 |
| Hmgcs1 | ENSMUSG00000093930 | 2020620004 | 6290347934 | 1864270207 | 389E-07 | 981E-06 |
| D430020J02Rik | ENSMUSG00000112980 | 2026288267 | 1842941361 | 4307964289 | 0000125762 | 0000543928 |
| C1qtnf12 | ENSMUSG00000023571 | 2036987079 | 3899849335 | 1187293414 | 247E-06 | 315E-05 |
| Slc39a4 | ENSMUSG00000063354 | 2037045182 | 3669892041 | 1065638049 | 382E-06 | 420E-05 |
| Abcb10 | ENSMUSG00000031974 | 2041160148 | 3517581435 | 1239527436 | 207E-06 | 283E-05 |
| Mybl2 | ENSMUSG00000017861 | 2041883847 | 3667897401 | 568968711 | 446E-05 | 0000244811 |
| Cep170b | ENSMUSG00000072825 | 2042589392 | 5581725746 | 3594178602 | 249E-08 | 189E-06 |
| Eno2 | ENSMUSG00000004267 | 2043552746 | 2019304536 | 5710917503 | 439E-05 | 0000242057 |
| Endod1 | ENSMUSG00000037419 | 2046655415 | 4947211201 | 1848882863 | 402E-07 | 996E-06 |
| Stmn1 | ENSMUSG00000028832 | 2047017426 | 6351453596 | 4234919815 | 0000133871 | 000057284 |
| Rhoq | ENSMUSG00000024143 | 2051649154 | 3677621404 | 8653165582 | 875E-06 | 724E-05 |
| Atxn1 | ENSMUSG00000046876 | 2053797335 | 5394956006 | 4411989352 | 105E-08 | 121E-06 |
| Pdcd1lg2 | ENSMUSG00000016498 | 2055878868 | 1556852671 | 7376756249 | 164E-05 | 0000114337 |
| Ncapd2 | ENSMUSG00000038252 | 2056753776 | 7231552156 | 8631843507 | 884E-06 | 728E-05 |
| Gpr171 | ENSMUSG00000050075 | 2057334508 | 6312632729 | 1604351003 | 722E-07 | 141E-05 |
| Plin2 | ENSMUSG00000028494 | 2060504219 | 454497096 | 102932179 | 438E-06 | 461E-05 |
| Cyfip1 | ENSMUSG00000030447 | 2063006454 | 4787036709 | 2223765195 | 187E-07 | 616E-06 |
| 4930556H04Rik | ENSMUSG00000114190 | 2063904157 | 1073918265 | 6052325268 | 352E-05 | 0000204088 |
| Pole | ENSMUSG00000007080 | 2065668367 | 4881013309 | 7656475238 | 142E-05 | 0000103238 |
| Wee1 | ENSMUSG00000031016 | 2066586419 | 3396791474 | 9111187152 | 714E-06 | 636E-05 |
| Niban1 | ENSMUSG00000026483 | 2069319733 | 5802996146 | 1902334984 | 357E-07 | 936E-06 |
| Cyp4f16 | ENSMUSG00000048440 | 2070276319 | 4302407515 | 2023492535 | 277E-07 | 793E-06 |
| Susd3 | ENSMUSG00000021384 | 2070669546 | 5741251741 | 4787471153 | 741E-09 | 105E-06 |
| 1810044D09Rik | ENSMUSG00000100680 | 2075001526 | 0773496546 | 5985251476 | 367E-05 | 0000210547 |
| Sema4f | ENSMUSG00000000627 | 2081248225 | 7129492776 | 5450789302 | 427E-09 | 779E-07 |
| Gsto1 | ENSMUSG00000025068 | 2082536201 | 4821818022 | 3695310995 | 222E-08 | 183E-06 |
| Cd81 | ENSMUSG00000037706 | 2083497621 | 5311958876 | 2934952965 | 585E-08 | 311E-06 |
| Gm14718 | ENSMUSG00000085808 | 2089122251 | 1253597733 | 6814835173 | 223E-05 | 0000144681 |
| Pglyrp1 | ENSMUSG00000030413 | 2090942092 | 4468508538 | 2028349917 | 274E-07 | 789E-06 |
| Ikbip | ENSMUSG00000019975 | 2095669354 | 1738797568 | 5179805423 | 635E-05 | 0000322519 |
| Rhoc | ENSMUSG00000002233 | 209571255 | 3072581098 | 1152368146 | 278E-06 | 339E-05 |
| Renbp | ENSMUSG00000031387 | 2095765543 | 1899329156 | 4316094282 | 0000124898 | 0000541192 |
| Eng | ENSMUSG00000026814 | 2099424363 | 4292083041 | 112967406 | 302E-06 | 358E-05 |
| Hip1r | ENSMUSG00000000915 | 2102662612 | 6967029074 | 2171157579 | 206E-07 | 654E-06 |
| Smim3 | ENSMUSG00000038059 | 2106224115 | 2217175555 | 4666223159 | 937E-05 | 0000434204 |
| Ptpn3 | ENSMUSG00000038764 | 2107648237 | 3699424392 | 1221295563 | 220E-06 | 294E-05 |
| Nicn1 | ENSMUSG00000032606 | 2111934638 | 3050942205 | 6293130204 | 303E-05 | 0000181277 |
| Mocos | ENSMUSG00000039616 | 211247153 | 2114451524 | 6712189035 | 237E-05 | 0000150585 |
| Slamf7 | ENSMUSG00000038179 | 2113298122 | 4665696818 | 1010628005 | 472E-06 | 483E-05 |
| Icos | ENSMUSG00000026009 | 2113938326 | 7083147955 | 1278381818 | 183E-06 | 262E-05 |
| Odf3b | ENSMUSG00000047394 | 2117528415 | 0635291673 | 7473690387 | 156E-05 | 0000110304 |
| Bcl2l14 | ENSMUSG00000030200 | 2123840545 | 1620388833 | 7111038483 | 189E-05 | 0000126968 |
| Csf2rb2 | ENSMUSG00000071714 | 2126064344 | 32219583 | 5126808534 | 660E-05 | 0000333316 |
| Fam161a | ENSMUSG00000049811 | 2127017696 | 1181360353 | 6897695454 | 213E-05 | 0000139302 |
| Trip10 | ENSMUSG00000019487 | 2128605534 | 3457543639 | 9984157344 | 495E-06 | 500E-05 |
| Spock2 | ENSMUSG00000058297 | 2128830393 | 4260553936 | 5036057022 | 706E-05 | 0000351046 |
| Ccl6 | ENSMUSG00000018927 | 2129757351 | 3337818212 | 5298945593 | 583E-05 | 0000301479 |
| Fzd7 | ENSMUSG00000041075 | 2129896193 | 2048247571 | 4574952787 | 0000100831 | 000045819 |
| Ift122 | ENSMUSG00000030323 | 2134634363 | 2629664237 | 6088313027 | 344E-05 | 0000200449 |
| F730043M19Rik | ENSMUSG00000052125 | 2135579741 | 2725445309 | 1063625061 | 384E-06 | 422E-05 |
| Prr13 | ENSMUSG00000023048 | 2136474257 | 6784683036 | 3982302314 | 162E-08 | 150E-06 |

| | | | | | | |
|---|---|---|---|---|---|---|
| Pafah1b3 | ENSMUSG00000005447 | 2136861681 | 3735317964 | 7021583761 | 199E-05 | 0000132305 |
| Eme1 | ENSMUSG00000039055 | 2143051969 | 2264176488 | 4178794023 | 0000140542 | 0000594413 |
| Il1rl1 | ENSMUSG00000026069 | 2178049626 | 2562078799 | 1125257256 | 306E-06 | 362E-05 |
| Camk2n1 | ENSMUSG00000046447 | 2179393209 | 2722775013 | 1136902851 | 294E-06 | 353E-05 |
| Angptl4 | ENSMUSG00000002289 | 2180263898 | 2740273855 | 9329199337 | 650E-06 | 594E-05 |
| Soat2 | ENSMUSG00000023045 | 2191554991 | 2544948295 | 6100156407 | 342E-05 | 0000199361 |
| Kpna2 | ENSMUSG00000018362 | 2205110691 | 5648342632 | 1255824646 | 197E-06 | 275E-05 |
| Tasl | ENSMUSG00000025058 | 221239465 | 1516664644 | 7025958099 | 198E-05 | 0000132136 |
| Jdp2 | ENSMUSG00000034271 | 2213157193 | 2048700317 | 6624647966 | 249E-05 | 0000155869 |
| Stard4 | ENSMUSG00000024378 | 2217953064 | 624592468 | 2672754494 | 867E-08 | 382E-06 |
| Slc9b2 | ENSMUSG00000037994 | 2218232021 | 1921549006 | 8472003396 | 952E-06 | 768E-05 |
| Brca1 | ENSMUSG00000017146 | 2219011943 | 4071286849 | 5369331349 | 555E-05 | 0000290725 |
| Itga2 | ENSMUSG00000015533 | 2222934364 | 4487605922 | 1736213521 | 522E-07 | 115E-05 |
| 2700099C18Rik | ENSMUSG00000098090 | 2223995505 | 2414932483 | 6507025363 | 267E-05 | 000016394 |
| Slc4a7 | ENSMUSG00000021733 | 2224045118 | 7235786096 | 6163240736 | 253E-09 | 603E-07 |
| Efcab11 | ENSMUSG00000021176 | 2224157379 | 0809906882 | 5412751023 | 538E-05 | 0000283939 |
| Rap1gap | ENSMUSG00000041351 | 2227422715 | 1009506212 | 4767370808 | 866E-05 | 0000406773 |
| Spdl1 | ENSMUSG00000069910 | 2231964313 | 2902793206 | 4922495453 | 768E-05 | 0000375917 |
| Gab2 | ENSMUSG00000004508 | 2235612752 | 2537292899 | 5616531955 | 468E-05 | 0000253902 |
| Ccdc39 | ENSMUSG00000027676 | 2236222856 | 206832558 | 990258915 | 512E-06 | 509E-05 |
| Lpxn | ENSMUSG00000024696 | 2244590169 | 6444728552 | 2777884574 | 737E-08 | 352E-06 |
| Slc43a3 | ENSMUSG00000027074 | 224869943 | 3692175198 | 5434566538 | 530E-05 | 0000280676 |
| Hook2 | ENSMUSG00000052566 | 2253337403 | 4427271624 | 2868814811 | 644E-08 | 330E-06 |
| Gm13241 | ENSMUSG00000073716 | 2253528531 | 0656763068 | 4883767188 | 791E-05 | 0000382893 |
| Igf2bp3 | ENSMUSG00000029814 | 2258069008 | 3413636406 | 5075806767 | 685E-05 | 0000343212 |
| Cdkn2c | ENSMUSG00000028551 | 2258989723 | 4031104902 | 6635896963 | 247E-05 | 0000155294 |
| Inppl1 | ENSMUSG00000032737 | 2261209954 | 4427421486 | 1096105558 | 341E-06 | 390E-05 |
| Bmp7 | ENSMUSG00000008999 | 2266306488 | 3136111569 | 5818262037 | 409E-05 | 0000229742 |
| 4930520O04Rik | ENSMUSG00000074039 | 2271202299 | 139407534 | 4779366568 | 858E-05 | 0000404531 |
| Cdc6 | ENSMUSG00000017499 | 2276227603 | 3260263168 | 5223043387 | 616E-05 | 0000313953 |
| Metrnl | ENSMUSG00000039208 | 2278170114 | 1714035469 | 9096206004 | 718E-06 | 637E-05 |
| Cdc45 | ENSMUSG00000000028 | 228679348 | 3445975296 | 7352655467 | 166E-05 | 0000115461 |
| BC064078 | ENSMUSG00000087150 | 22872495 | 1646304328 | 5682091265 | 448E-05 | 0000245404 |
| Pask | ENSMUSG00000026274 | 2289938643 | 255009062 | 633303805 | 296E-05 | 0000178386 |
| Evi5 | ENSMUSG00000011831 | 2294224227 | 2470203376 | 1105054467 | 330E-06 | 381E-05 |
| H2-Oa | ENSMUSG00000024334 | 230620328 | 3811227241 | 1189900373 | 245E-06 | 313E-05 |
| Gm10505 | ENSMUSG00000073430 | 2312870759 | 2446269459 | 1080396314 | 361E-06 | 404E-05 |
| Spef2 | ENSMUSG00000072663 | 2320990465 | 4792380337 | 4642039671 | 845E-09 | 108E-06 |
| Fbxo48 | ENSMUSG00000044966 | 2328039224 | 1769933018 | 6132901606 | 335E-05 | 00001962 |
| Amigo1 | ENSMUSG00000050947 | 2330331292 | 3571561851 | 2988730403 | 542E-08 | 296E-06 |
| Cks1b | ENSMUSG00000028044 | 2333659287 | 4109522103 | 6750891093 | 231E-05 | 0000148435 |
| Basp1 | ENSMUSG00000045763 | 2335897996 | 4267515742 | 6834656092 | 221E-05 | 0000143382 |
| Pla2g12a | ENSMUSG00000027999 | 2347184609 | 4731989195 | 3234285133 | 389E-08 | 249E-06 |
| Bag2 | ENSMUSG00000042215 | 235735457 | 1839748058 | 4786369622 | 853E-05 | 0000402826 |
| Agpat4 | ENSMUSG00000023827 | 2362416377 | 3632128313 | 103887954 | 423E-06 | 449E-05 |
| Tlcd2 | ENSMUSG00000038217 | 2369394762 | 0946136279 | 7564687498 | 149E-05 | 0000106499 |
| Ybx3 | ENSMUSG00000030189 | 2373389085 | 641916215 | 2998869912 | 535E-08 | 296E-06 |
| Fancd2 | ENSMUSG00000034023 | 2373561998 | 3259231479 | 4610057514 | 980E-05 | 000044804 |
| Arhgap11a | ENSMUSG00000041219 | 2374168926 | 5538586244 | 104356237 | 415E-06 | 444E-05 |
| Map3k20 | ENSMUSG00000004085 | 2377281014 | 1589178994 | 8690134591 | 861E-06 | 716E-05 |
| Ltbp4 | ENSMUSG00000040488 | 2379468898 | 1393705703 | 6577837713 | 256E-05 | 0000159025 |
| Cxcr3 | ENSMUSG00000050232 | 2387738396 | 4386544959 | 5558547669 | 487E-05 | 0000262374 |
| Gpld1 | ENSMUSG00000021340 | 2389161555 | 2455939188 | 7119418242 | 188E-05 | 0000126461 |
| C1qtnf4 | ENSMUSG00000040794 | 2394323373 | 1234019397 | 9348256105 | 644E-06 | 591E-05 |
| Arhgap21 | ENSMUSG00000036591 | 2398732596 | 4235382472 | 1680664707 | 596E-07 | 125E-05 |
| Tent5a | ENSMUSG00000032265 | 2403097826 | 7091506792 | 8049710833 | 811E-10 | 347E-07 |
| Slc2a6 | ENSMUSG00000036067 | 2407570686 | 2383464804 | 8340500913 | 101E-05 | 804E-05 |

| | | | | | | |
|---|---|---|---|---|---|---|
| Fzd6 | ENSMUSG00000022297 | 2408312132 | 1089886667 | 3701953121 | 000021757 | 000082456 |
| Acot7 | ENSMUSG00000028937 | 2411858586 | 5426886633 | 1896493052 | 362E-07 | 941E-06 |
| Serinc3 | ENSMUSG00000017707 | 2418524234 | 9629957425 | 8503300786 | 642E-10 | 347E-07 |
| Casp3 | ENSMUSG00000031628 | 2422871814 | 6210056559 | 192776219 | 338E-07 | 906E-06 |
| Ier3 | ENSMUSG00000003541 | 2428948759 | 295373947 | 7739335324 | 136E-05 | 100E-04 |
| Plscr1 | ENSMUSG00000032369 | 2429882158 | 3290260373 | 1252054582 | 199E-06 | 277E-05 |
| Stau2 | ENSMUSG00000025920 | 2431440035 | 1279381309 | 6407754383 | 283E-05 | 0000171855 |
| Stx11 | ENSMUSG00000039232 | 2448250663 | 3965195996 | 1238915678 | 208E-06 | 284E-05 |
| Dmwd | ENSMUSG00000030410 | 2449755757 | 1797808177 | 4162974671 | 0000142496 | 000060092 |
| Igkv4-57 | ENSMUSG00000076556 | 2455870644 | 096997995 | 4126750479 | 0000147098 | 0000616112 |
| Fam78b | ENSMUSG00000060568 | 2470121327 | 1317242413 | 4307290969 | 0000125834 | 0000544038 |
| Pltp | ENSMUSG00000017754 | 2482364843 | 1049629419 | 4247202737 | 0000132463 | 0000567858 |
| Smtn | ENSMUSG00000020439 | 2482982881 | 1696925963 | 3491029559 | 0000267996 | 0000970893 |
| Gins2 | ENSMUSG00000031821 | 2483596423 | 2371177542 | 7051126447 | 195E-05 | 0000130537 |
| Knstrn | ENSMUSG00000027331 | 2484569483 | 4483913857 | 1145227656 | 285E-06 | 346E-05 |
| NA | ENSMUSG00000091562 | 2485597878 | 1523009569 | 644278279 | 277E-05 | 0000168969 |
| Pxmp2 | ENSMUSG00000029499 | 2485789301 | 0415333753 | 429109629 | 000012758 | 0000549548 |
| Akap1 | ENSMUSG00000018428 | 2488291989 | 3794618022 | 1738415158 | 519E-07 | 115E-05 |
| Serpinb6a | ENSMUSG00000060147 | 2491070543 | 5747205756 | 1611269163 | 709E-07 | 139E-05 |
| Cercam | ENSMUSG00000039787 | 2494124796 | 2330110486 | 5103436992 | 672E-05 | 0000337465 |
| Tmprss13 | ENSMUSG00000037129 | 2497083903 | 2004173128 | 3804762034 | 0000197249 | 0000767489 |
| Tacc3 | ENSMUSG00000037313 | 2498516589 | 5302213407 | 9201023494 | 686E-06 | 618E-05 |
| Tk1 | ENSMUSG00000025574 | 250160136 | 4748726132 | 2269812677 | 172E-07 | 588E-06 |
| Plpp2 | ENSMUSG00000052151 | 2507220916 | 2418261216 | 1042300446 | 417E-06 | 445E-05 |
| Marchf8 | ENSMUSG00000025702 | 2512059689 | 2662700333 | 1242960189 | 205E-06 | 282E-05 |
| Fignl1 | ENSMUSG00000035455 | 2523143048 | 3719766483 | 7333806869 | 168E-05 | 0000116346 |
| Scpep1 | ENSMUSG00000000278 | 2526816253 | 4206494707 | 1161511833 | 270E-06 | 331E-05 |
| Tbkbp1 | ENSMUSG00000038517 | 2534853102 | 3144075181 | 7522436573 | 152E-05 | 0000108524 |
| Six5 | ENSMUSG00000040841 | 2538412132 | 0602626656 | 5357279364 | 560E-05 | 0000292935 |
| Cox6b2 | ENSMUSG00000051811 | 2541675989 | 1406817363 | 5270485102 | 595E-05 | 000030535 |
| Med12l | ENSMUSG00000056476 | 255175938 | 3582745304 | 2386963472 | 139E-07 | 516E-06 |
| Cryz | ENSMUSG00000028199 | 2556034657 | 2722031232 | 139140713 | 129E-06 | 208E-05 |
| Plcxd1 | ENSMUSG00000064247 | 2558174607 | 2525655776 | 490477787 | 779E-05 | 0000379019 |
| Gabarapl1 | ENSMUSG00000030161 | 2568569049 | 355593426 | 1613495059 | 705E-07 | 139E-05 |
| Neb | ENSMUSG00000026950 | 2570527897 | 5095073561 | 1123241822 | 309E-06 | 364E-05 |
| Nsl1 | ENSMUSG00000062510 | 2570603162 | 3010454913 | 8356330543 | 100E-05 | 800E-05 |
| Dhdh | ENSMUSG00000011382 | 2570987241 | 2202110623 | 9545130361 | 593E-06 | 559E-05 |
| Uhrf1 | ENSMUSG00000001228 | 2571643137 | 5566843772 | 9006681223 | 747E-06 | 654E-05 |
| Acsbg1 | ENSMUSG00000032281 | 2575982768 | 5707307084 | 3155717293 | 432E-08 | 260E-06 |
| Ccdc18 | ENSMUSG00000056531 | 2576125325 | 1965400102 | 661234842 | 251E-05 | 0000156654 |
| Anxa1 | ENSMUSG00000024659 | 2579971118 | 4139535456 | 1319689825 | 161E-06 | 242E-05 |
| Arhgap19 | ENSMUSG00000025154 | 2581784924 | 4367644509 | 1019476567 | 456E-06 | 471E-05 |
| Spc24 | ENSMUSG00000074476 | 2587581825 | 3146763651 | 7367802208 | 165E-05 | 0000114743 |
| E2f8 | ENSMUSG00000046179 | 2587733432 | 3493338274 | 4473233525 | 000010954 | 00004888 |
| Sgo1 | ENSMUSG00000023940 | 2588336869 | 3668691927 | 938954542 | 633E-06 | 584E-05 |
| 5830418P13Rik | ENSMUSG00000086236 | 2589372196 | 0390507577 | 8921713262 | 776E-06 | 674E-05 |
| Chaf1a | ENSMUSG00000002835 | 2590105822 | 3972202461 | 905452592 | 731E-06 | 645E-05 |
| Rapgef3 | ENSMUSG00000022469 | 260861652 | 1410183434 | 6902024382 | 212E-05 | 0000139119 |
| Ndnf | ENSMUSG00000049001 | 2615047051 | 1012809815 | 3962944922 | 0000170348 | 0000691191 |
| Septin8 | ENSMUSG00000018398 | 261629487 | 4653584049 | 5539743066 | 399E-09 | 739E-07 |
| Rap1gap2 | ENSMUSG00000038807 | 2625419479 | 5527004989 | 1671995694 | 609E-07 | 126E-05 |
| Cela1 | ENSMUSG00000023031 | 2629157724 | 1565236322 | 6498556328 | 268E-05 | 0000164505 |
| Ect2 | ENSMUSG00000027699 | 2632257319 | 4075288146 | 1165181547 | 266E-06 | 329E-05 |
| Plk1 | ENSMUSG00000030867 | 2633196012 | 4116887124 | 9717749422 | 552E-06 | 535E-05 |
| Myo1f | ENSMUSG00000024300 | 2634565164 | 672721605 | 1000508263 | 491E-06 | 497E-05 |
| Cip2a | ENSMUSG00000033031 | 2636457237 | 4505805475 | 1416366186 | 120E-06 | 197E-05 |
| Sh3gl3 | ENSMUSG00000030638 | 2648739627 | 1679177241 | 1287399316 | 178E-06 | 258E-05 |

| | | | | | | |
|---|---|---|---|---|---|---|
| Ugt1a7c | ENSMUSG00000090124 | 2670558525 | 0689786392 | 578623278 | 418E-05 | 0000233384 |
| Racgap1 | ENSMUSG00000023015 | 2677386066 | 6088299057 | 1226124971 | 217E-06 | 291E-05 |
| Casp1 | ENSMUSG00000025888 | 2678031592 | 415235931 | 3147046331 | 437E-08 | 261E-06 |
| Hpse | ENSMUSG00000035273 | 2690944798 | 2730623146 | 5050913777 | 698E-05 | 0000347941 |
| Fgl2 | ENSMUSG00000039899 | 2704984726 | 4398904949 | 9065382707 | 728E-06 | 643E-05 |
| Anxa9 | ENSMUSG00000015702 | 2710218842 | 1796653097 | 1685916517 | 589E-07 | 124E-05 |
| St14 | ENSMUSG00000031995 | 2716455208 | 3377330549 | 1744987047 | 511E-07 | 115E-05 |
| Rad54l | ENSMUSG00000028702 | 2723431481 | 3154039471 | 8210895328 | 108E-05 | 838E-05 |
| Cenpi | ENSMUSG00000031262 | 2735367845 | 2631191733 | 8781849925 | 826E-06 | 700E-05 |
| Dlgap5 | ENSMUSG00000037544 | 2748954432 | 4934714918 | 1204381031 | 233E-06 | 304E-05 |
| Rnf208 | ENSMUSG00000044628 | 2751480063 | 0787666719 | 7929523275 | 124E-05 | 929E-05 |
| Gtse1 | ENSMUSG00000022385 | 2751712956 | 4142663747 | 1794899908 | 455E-07 | 108E-05 |
| Cacna1c | ENSMUSG00000051331 | 2752008614 | 0523454776 | 3693064438 | 0000219445 | 0000828975 |
| Akap7 | ENSMUSG00000039166 | 2753474453 | 3456886589 | 2682690945 | 853E-08 | 382E-06 |
| St6galnac6 | ENSMUSG00000026811 | 2773505059 | 3239035564 | 1897371138 | 361E-07 | 941E-06 |
| Hemgn | ENSMUSG00000028332 | 2778446604 | 2167517704 | 4497154389 | 0000107411 | 00004826 |
| Snx9 | ENSMUSG00000002365 | 2790758068 | 5254300627 | 4688890852 | 810E-09 | 108E-06 |
| Ticrr | ENSMUSG00000046591 | 2801298943 | 3753398952 | 8056596206 | 116E-05 | 887E-05 |
| Ctnnbip1 | ENSMUSG00000028988 | 2806732476 | 143122423 | 1226040922 | 217E-06 | 291E-05 |
| Tgfb3 | ENSMUSG00000021253 | 281199046 | 2243965386 | 9227827449 | 678E-06 | 614E-05 |
| Dtl | ENSMUSG00000037474 | 2813161131 | 3935607172 | 7442445392 | 158E-05 | 0000111447 |
| Tmem107 | ENSMUSG00000020895 | 2838760198 | 0698510261 | 5774237832 | 421E-05 | 0000234671 |
| Niban2 | ENSMUSG00000026796 | 2842556001 | 3447720851 | 8176214352 | 110E-05 | 846E-05 |
| Havcr2 | ENSMUSG00000020399 | 2856945925 | 1550280251 | 4411387901 | 000011529 | 0000508435 |
| Pip5k1b | ENSMUSG00000024867 | 2857765641 | 329618914 | 2195230948 | 197E-07 | 636E-06 |
| Ncaph | ENSMUSG00000034906 | 2866836568 | 4626580872 | 1099782104 | 336E-06 | 387E-05 |
| Asb2 | ENSMUSG00000021200 | 2866896228 | 4230645174 | 1427092406 | 117E-06 | 193E-05 |
| Aurka | ENSMUSG00000027496 | 2866932927 | 4097949392 | 105092533 | 403E-06 | 436E-05 |
| Rad51ap1 | ENSMUSG00000030346 | 2869385732 | 3095224185 | 1133878989 | 297E-06 | 354E-05 |
| Stk32c | ENSMUSG00000015981 | 2874377926 | 1404637229 | 553772449 | 494E-05 | 0000265515 |
| Entpd1 | ENSMUSG00000048120 | 2875318474 | 3260001897 | 1258303254 | 195E-06 | 274E-05 |
| Foxm1 | ENSMUSG00000001517 | 2893953624 | 4805338458 | 1087665735 | 351E-06 | 396E-05 |
| Anln | ENSMUSG00000036777 | 2896010374 | 3601327938 | 1229357397 | 214E-06 | 290E-05 |
| Krt18 | ENSMUSG00000023043 | 2897149109 | 1030316097 | 348176715 | 0000270524 | 0000976336 |
| Ncapg2 | ENSMUSG00000042029 | 2901221375 | 4918035391 | 1146431301 | 284E-06 | 345E-05 |
| Pygl | ENSMUSG00000021069 | 2906382412 | 3739651979 | 5829879289 | 406E-05 | 0000228223 |
| Cpd | ENSMUSG00000020841 | 2917780521 | 3272980972 | 9967604763 | 499E-06 | 502E-05 |
| Cenph | ENSMUSG00000045273 | 2924993288 | 2296146789 | 7302149952 | 171E-05 | 0000117904 |
| Patl2 | ENSMUSG00000027233 | 2929206973 | 0759624173 | 4518199779 | 000010558 | 0000476232 |
| Cenpk | ENSMUSG00000021714 | 2933033276 | 1673172827 | 1569850526 | 790E-07 | 148E-05 |
| Tiam1 | ENSMUSG00000002489 | 2937334704 | 6074630993 | 1919653539 | 344E-07 | 916E-06 |
| Rgs9 | ENSMUSG00000020599 | 2942448666 | 0775282654 | 5502244252 | 506E-05 | 000027049 |
| Trip13 | ENSMUSG00000021569 | 2947705864 | 2611946706 | 7980122022 | 120E-05 | 911E-05 |
| Anxa4 | ENSMUSG00000029994 | 2977350105 | 2931064013 | 8146843735 | 111E-05 | 856E-05 |
| Aqp9 | ENSMUSG00000032204 | 2977729377 | 3953387755 | 2558070556 | 104E-07 | 423E-06 |
| Micall2 | ENSMUSG00000036718 | 2985462157 | 109159138 | 7587856579 | 147E-05 | 0000105622 |
| Rrm2 | ENSMUSG00000020649 | 2991539194 | 5491692285 | 8798751844 | 820E-06 | 698E-05 |
| Alcam | ENSMUSG00000022636 | 2994934635 | 4427521462 | 1101793599 | 334E-06 | 385E-05 |
| Gm31718 | ENSMUSG00000110697 | 2997490755 | 129645169 | 8759779211 | 834E-06 | 703E-05 |
| Mzb1 | ENSMUSG00000024353 | 30077976 | 4226028961 | 3725252208 | 0000212746 | 0000810228 |
| Ska3 | ENSMUSG00000021965 | 3017404285 | 2912831163 | 1060364317 | 389E-06 | 426E-05 |
| Kif20a | ENSMUSG00000003779 | 3024257156 | 4524983012 | 1185612331 | 248E-06 | 316E-05 |
| Fam20a | ENSMUSG00000020614 | 3026755069 | 1277093558 | 3835244781 | 0000191679 | 0000753769 |
| Diaph3 | ENSMUSG00000022021 | 3027588228 | 4272113134 | 9704913902 | 555E-06 | 536E-05 |
| Dyrk3 | ENSMUSG00000016526 | 3029540005 | 2398416373 | 1292038271 | 175E-06 | 257E-05 |
| Rad54b | ENSMUSG00000078773 | 3038857278 | 2234457538 | 7736804234 | 136E-05 | 0000100015 |
| Cxcr1 | ENSMUSG00000048480 | 3042052418 | 2037084209 | 1059935498 | 390E-06 | 426E-05 |

| | | | | | | |
|---|---|---|---|---|---|---|
| Ell2 | ENSMUSG00000001542 | 3062829814 | 4278579286 | 2966488089 | 560E-08 | 302E-06 |
| Oip5 | ENSMUSG00000072980 | 3062877709 | 1901077601 | 7443396363 | 158E-05 | 0000111447 |
| Spc25 | ENSMUSG00000005233 | 3067078908 | 3239312017 | 1027757093 | 441E-06 | 462E-05 |
| Mcm10 | ENSMUSG00000026669 | 3078459636 | 3283637137 | 7994657725 | 120E-05 | 907E-05 |
| Knl1 | ENSMUSG00000027326 | 3090095405 | 4535590997 | 1132122544 | 299E-06 | 356E-05 |
| Itpripl2 | ENSMUSG00000095115 | 3100796819 | 4193918832 | 1977934985 | 304E-07 | 855E-06 |
| Mis18bp1 | ENSMUSG00000047534 | 3102960171 | 3990779121 | 1325425139 | 158E-06 | 238E-05 |
| Svip | ENSMUSG00000074093 | 310417289 | 1361615031 | 789795449 | 125E-05 | 939E-05 |
| Prc1 | ENSMUSG00000038943 | 3104449216 | 4817771395 | 1153260881 | 277E-06 | 339E-05 |
| Casp4 | ENSMUSG00000033538 | 3107387839 | 3430888518 | 7636823445 | 143E-05 | 0000103829 |
| Sgo2a | ENSMUSG00000026039 | 3143902188 | 1907202793 | 6022544615 | 359E-05 | 0000206986 |
| Gm15987 | ENSMUSG00000085786 | 3145365154 | 2202749948 | 4232148355 | 0000134191 | 0000573789 |
| Xkr5 | ENSMUSG00000039814 | 3152954134 | 1578836349 | 8095787603 | 114E-05 | 874E-05 |
| E2f7 | ENSMUSG00000020185 | 3180866573 | 2250126832 | 4892123576 | 786E-05 | 0000381095 |
| Arrdc4 | ENSMUSG00000042659 | 3182263164 | 3392455161 | 1002843531 | 487E-06 | 494E-05 |
| Trim16 | ENSMUSG00000047821 | 3185843456 | 2905064309 | 1677857133 | 601E-07 | 125E-05 |
| Hrob | ENSMUSG00000034773 | 3194385573 | 2409135912 | 5810248069 | 411E-05 | 0000230396 |
| Mctp1 | ENSMUSG00000021596 | 3211125004 | 2927619206 | 1442328626 | 112E-06 | 187E-05 |
| Rad51 | ENSMUSG00000027323 | 321504709 | 383584867 | 9417327845 | 626E-06 | 579E-05 |
| Clspn | ENSMUSG00000042489 | 3219777416 | 4300136725 | 8388030488 | 990E-06 | 790E-05 |
| Cenpp | ENSMUSG00000021391 | 3227988233 | 1548661044 | 3664532069 | 0000225602 | 0000845934 |
| Dnah12 | ENSMUSG00000021879 | 3231029924 | 3089526089 | 2901200832 | 614E-08 | 320E-06 |
| Kntc1 | ENSMUSG00000029414 | 3231497008 | 4350134708 | 9684702552 | 560E-06 | 537E-05 |
| Esco2 | ENSMUSG00000022034 | 3240101797 | 3474008097 | 905334006 | 732E-06 | 645E-05 |
| Pbk | ENSMUSG00000022033 | 3256339512 | 2447658753 | 5929526173 | 381E-05 | 0000216767 |
| Cdca5 | ENSMUSG00000024791 | 3259056445 | 3545068875 | 1292126742 | 175E-06 | 257E-05 |
| Tmem40 | ENSMUSG00000059900 | 3275353051 | 1697665572 | 9058845756 | 730E-06 | 644E-05 |
| Rai14 | ENSMUSG00000022246 | 3276120017 | 0869744961 | 1169594361 | 262E-06 | 325E-05 |
| Cit | ENSMUSG00000029516 | 3290400317 | 4750824304 | 169005828 | 583E-07 | 124E-05 |
| Ccnf | ENSMUSG00000072082 | 3303690922 | 4106043086 | 1213307522 | 226E-06 | 300E-05 |
| Plxnb3 | ENSMUSG00000031385 | 3335830907 | 0479628559 | 1360452574 | 142E-06 | 222E-05 |
| Tgm2 | ENSMUSG00000037820 | 3339008071 | 2446357094 | 1586034887 | 757E-07 | 145E-05 |
| Cdca8 | ENSMUSG00000028873 | 3344100861 | 4654403176 | 1446078057 | 111E-06 | 186E-05 |
| Rapgef5 | ENSMUSG00000041992 | 334968312 | 2704934011 | 9216136546 | 682E-06 | 615E-05 |
| NA | ENSMUSG00000072812 | 3356169955 | 079962137 | 3719045806 | 0000214018 | 0000813213 |
| Cysltr2 | ENSMUSG00000033470 | 3361431558 | 347887975 | 6278949233 | 306E-05 | 0000182666 |
| Cdca2 | ENSMUSG00000048922 | 3368236798 | 4179199452 | 178786946 | 462E-07 | 109E-05 |
| Cdkn3 | ENSMUSG00000037628 | 3370894277 | 2516119281 | 1211361952 | 227E-06 | 301E-05 |
| Depdc1b | ENSMUSG00000021697 | 3373273099 | 3294056109 | 8904141352 | 782E-06 | 676E-05 |
| Vwf | ENSMUSG00000001930 | 3377270925 | 2627922366 | 6530187245 | 263E-05 | 0000162395 |
| Cdca3 | ENSMUSG00000023505 | 338921478 | 4844435117 | 1106911479 | 327E-06 | 379E-05 |
| Bub1b | ENSMUSG00000040084 | 3390436827 | 5311139542 | 1254388302 | 197E-06 | 276E-05 |
| Itgb5 | ENSMUSG00000022817 | 3392535239 | 2342575969 | 5417217266 | 537E-05 | 0000283304 |
| Hacd1 | ENSMUSG00000063275 | 3396118522 | 0688931841 | 1624619744 | 686E-07 | 137E-05 |
| Bub1 | ENSMUSG00000027379 | 3414549455 | 3934849672 | 1086497651 | 353E-06 | 398E-05 |
| Ankle1 | ENSMUSG00000046295 | 3420870157 | 3346777363 | 1442903551 | 112E-06 | 187E-05 |
| Fgfr1 | ENSMUSG00000031565 | 3426063971 | 2977434109 | 1039872054 | 421E-06 | 448E-05 |
| Cma1 | ENSMUSG00000022225 | 3447567866 | 305203774 | 1802816962 | 446E-07 | 107E-05 |
| Melk | ENSMUSG00000035683 | 344761673 | 3063902 | 8743768886 | 840E-06 | 705E-05 |
| Kif11 | ENSMUSG00000012443 | 3468217583 | 5798031916 | 1542832834 | 848E-07 | 155E-05 |
| Vdr | ENSMUSG00000022479 | 3471418345 | 1713685285 | 1539573092 | 855E-07 | 155E-05 |
| Cenpm | ENSMUSG00000068101 | 3485864651 | 2473566345 | 1217414023 | 223E-06 | 298E-05 |
| Ckap2l | ENSMUSG00000048327 | 3496949039 | 4074898541 | 1711645926 | 553E-07 | 120E-05 |
| Axl | ENSMUSG00000002602 | 3505153973 | 5345370783 | 14761981 | 102E-06 | 175E-05 |
| Kif15 | ENSMUSG00000036768 | 351582615 | 5049063299 | 1291259126 | 176E-06 | 257E-05 |
| Gas2l3 | ENSMUSG00000074802 | 3528924274 | 2524239073 | 9958674557 | 501E-06 | 503E-05 |
| Kif2c | ENSMUSG00000028678 | 356238776 | 3940233891 | 2226293914 | 186E-07 | 616E-06 |

| | | | | | | |
|---|---|---|---|---|---|---|
| Shcbp1 | ENSMUSG00000022322 | 3581792995 | 3325474857 | 1016121505 | 462E-06 | 475E-05 |
| P2ry1 | ENSMUSG00000027765 | 360551012 | 1969637458 | 4916204713 | 772E-05 | 0000376967 |
| Depdc1a | ENSMUSG00000028175 | 3605930561 | 2954707116 | 1260741771 | 193E-06 | 273E-05 |
| Nhsl2 | ENSMUSG00000079481 | 361975741 | 3695566276 | 964132518 | 570E-06 | 540E-05 |
| Mki67 | ENSMUSG00000031004 | 3631747563 | 8116930171 | 1696600072 | 574E-07 | 122E-05 |
| Ska1 | ENSMUSG00000036223 | 3646090485 | 1929251002 | 1166923265 | 265E-06 | 327E-05 |
| Iqgap3 | ENSMUSG00000028068 | 3662603262 | 3053891879 | 879609773 | 821E-06 | 698E-05 |
| Stil | ENSMUSG00000028718 | 3698323359 | 371433245 | 1314243555 | 163E-06 | 245E-05 |
| Ckap2 | ENSMUSG00000037725 | 3707189835 | 3914945823 | 1877507008 | 377E-07 | 967E-06 |
| Oaf | ENSMUSG00000032014 | 3719617984 | 164848845 | 9560842791 | 589E-06 | 556E-05 |
| Pclaf | ENSMUSG00000040204 | 3724430133 | 5221458994 | 112615402 | 305E-06 | 361E-05 |
| Kif18b | ENSMUSG00000051378 | 3735348491 | 4052743314 | 1742829344 | 513E-07 | 115E-05 |
| Top2a | ENSMUSG00000020914 | 3735651702 | 7933454038 | 1703182528 | 565E-07 | 120E-05 |
| Tpx2 | ENSMUSG00000027469 | 3751429741 | 5447248462 | 1547959256 | 837E-07 | 154E-05 |
| Selenbp1 | ENSMUSG00000068874 | 3762373192 | 4179893512 | 2102343757 | 236E-07 | 716E-06 |
| Cdk1 | ENSMUSG00000019942 | 381520242 | 4478133727 | 1480328106 | 100E-06 | 174E-05 |
| Parpbp | ENSMUSG00000035365 | 381570505 | 2795097114 | 1750085118 | 505E-07 | 114E-05 |
| Mastl | ENSMUSG00000026779 | 3822874623 | 3126752363 | 9444535277 | 618E-06 | 575E-05 |
| Ttk | ENSMUSG00000038379 | 383779241 | 301701323 | 9383251974 | 635E-06 | 585E-05 |
| Mxd3 | ENSMUSG00000021485 | 3847987704 | 3286352091 | 1068185808 | 378E-06 | 417E-05 |
| Spag5 | ENSMUSG00000002055 | 3873712078 | 4769336804 | 1326445313 | 157E-06 | 238E-05 |
| Kif4 | ENSMUSG00000034311 | 3880362569 | 4335558615 | 863651187 | 882E-06 | 728E-05 |
| Cenpe | ENSMUSG00000045328 | 3882519738 | 5674756758 | 187990593 | 375E-07 | 967E-06 |
| Slc22a21 | ENSMUSG00000063652 | 3884413642 | 0529248724 | 1162902281 | 268E-06 | 331E-05 |
| Ccnb2 | ENSMUSG00000032218 | 3894067012 | 5076577953 | 1424783173 | 117E-06 | 193E-05 |
| Ncapg | ENSMUSG00000015880 | 3905948007 | 4813558018 | 2106489659 | 234E-07 | 713E-06 |
| Bhlhe41 | ENSMUSG00000030256 | 3927437214 | 2889430392 | 2068336977 | 253E-07 | 752E-06 |
| Nuf2 | ENSMUSG00000026683 | 393038098 | 387769539 | 1645724693 | 650E-07 | 131E-05 |
| Espl1 | ENSMUSG00000058290 | 3989387388 | 5042726689 | 192943841 | 337E-07 | 906E-06 |
| Kif14 | ENSMUSG00000041498 | 4027929439 | 4114110922 | 1192705254 | 242E-06 | 311E-05 |
| Nek2 | ENSMUSG00000026622 | 4032072082 | 43479741 | 2143131102 | 218E-07 | 678E-06 |
| Aurkb | ENSMUSG00000020897 | 4061686819 | 4888975141 | 1660486596 | 627E-07 | 129E-05 |
| Neil3 | ENSMUSG00000039396 | 4078015193 | 4167487299 | 2232253957 | 184E-07 | 616E-06 |
| Neurl1b | ENSMUSG00000034413 | 40806512 | 2186721911 | 133150467 | 155E-06 | 236E-05 |
| Pif1 | ENSMUSG00000041064 | 4083479277 | 337260304 | 1029968469 | 437E-06 | 460E-05 |
| Prr11 | ENSMUSG00000020493 | 4088157875 | 4024856169 | 1915398724 | 347E-07 | 922E-06 |
| Nusap1 | ENSMUSG00000027306 | 4117718344 | 5312564453 | 220058288 | 195E-07 | 631E-06 |
| Syngr1 | ENSMUSG00000022415 | 4123451561 | 1881577008 | 9785585128 | 537E-06 | 525E-05 |
| Ccnb1 | ENSMUSG00000041431 | 4127135876 | 4842617747 | 1531767507 | 873E-07 | 158E-05 |
| Ube2c | ENSMUSG00000001403 | 4149705144 | 5177823082 | 3171599529 | 423E-08 | 257E-06 |
| Ltbp3 | ENSMUSG00000024940 | 4166577509 | 2798573117 | 1462508817 | 106E-06 | 181E-05 |
| Birc5 | ENSMUSG00000017716 | 4166743866 | 4959174908 | 136259939 | 141E-06 | 222E-05 |
| Ccna2 | ENSMUSG00000027715 | 4177951664 | 5452845283 | 1342529267 | 150E-06 | 232E-05 |
| Lsr | ENSMUSG00000001247 | 4214403258 | 2267588029 | 1592031889 | 745E-07 | 144E-05 |
| Hmmr | ENSMUSG00000020330 | 4287122365 | 4570851692 | 1035127776 | 429E-06 | 454E-05 |
| Aspm | ENSMUSG00000033952 | 430784643 | 5344557873 | 2794897 | 719E-08 | 352E-06 |
| Atp6v0d2 | ENSMUSG00000028238 | 4339204353 | 1803353065 | 1191808397 | 243E-06 | 311E-05 |
| Ceacam16 | ENSMUSG00000014686 | 4352310215 | 2277349917 | 9218213803 | 681E-06 | 615E-05 |
| Cdh17 | ENSMUSG00000028217 | 4444675914 | 1528513277 | 1106894678 | 327E-06 | 379E-05 |
| Cdc25c | ENSMUSG00000044201 | 4466156426 | 2449935209 | 1275537429 | 184E-06 | 264E-05 |
| Pcyt1b | ENSMUSG00000035246 | 4570804385 | 3329873205 | 235043356 | 148E-07 | 538E-06 |
| Spred3 | ENSMUSG00000037239 | 4581296421 | 2967159102 | 1523927431 | 892E-07 | 159E-05 |
| Ryk | ENSMUSG00000032547 | 4642567757 | 2852705961 | 1537205497 | 861E-07 | 156E-05 |
| Lrrc32 | ENSMUSG00000090958 | 4683756305 | 4638249588 | 125368606 | 198E-06 | 276E-05 |
| Fads2 | ENSMUSG00000024665 | 4718197787 | 2934347773 | 1749624402 | 505E-07 | 114E-05 |
| Ccr2 | ENSMUSG00000049103 | 4907108805 | 7862206958 | 1739490599 | 518E-07 | 115E-05 |
| Mcpt1 | ENSMUSG00000022227 | 4925658983 | 1409045822 | 998429646 | 495E-06 | 500E-05 |

| | | | | | | |
|---|---|---|---|---|---|---|
| Asns | ENSMUSG00000029752 | 4937687121 | 3656387293 | 3603403826 | 247E-08 | 189E-06 |
| Jcad | ENSMUSG00000033960 | 5110883833 | 2893792932 | 2670558046 | 870E-08 | 382E-06 |
| Il21 | ENSMUSG00000027718 | 5393973609 | 3686484944 | 1716969532 | 546E-07 | 119E-05 |
| Muc2 | ENSMUSG00000025515 | 57169715 | 1053403901 | 8212818881 | 108E-05 | 838E-05 |
| Ltb4r1 | ENSMUSG00000046908 | 5744075609 | 3734980846 | 1673586202 | 607E-07 | 126E-05 |
| Gm44659 | ENSMUSG00000109214 | 5908516205 | 0284414852 | 3453682845 | 0000278372 | 0000999178 |
| Mgl2 | ENSMUSG00000040950 | 6068783318 | 1108547593 | 9441176747 | 619E-06 | 575E-05 |
| Mcpt4 | ENSMUSG00000061068 | 6134152421 | 1584300256 | 1969211826 | 310E-07 | 869E-06 |
| Mmp12 | ENSMUSG00000049723 | 6167088994 | 2645063473 | 6818082488 | 223E-05 | 0000144575 |
| Olfr525 | ENSMUSG00000061489 | 6292913884 | 0208601692 | 1376208082 | 135E-06 | 216E-05 |
| Mgll | ENSMUSG00000033174 | 6439781022 | 4464331906 | 2658352397 | 887E-08 | 386E-06 |
| Gm49180 | ENSMUSG00000115735 | 7239182789 | 0482183597 | 4516918616 | 000010569 | 0000476361 |
| Timd2 | ENSMUSG00000040413 | 7241272674 | 1064177765 | 5718005134 | 437E-05 | 000024149 |
| Pparg | ENSMUSG00000000440 | 7348959875 | 3563119135 | 9277924214 | 442E-10 | 323E-07 |
| Akr1b8 | ENSMUSG00000029762 | 9151027511 | 0342821368 | 1438028703 | 113E-06 | 189E-05 |
| Fam183b | ENSMUSG00000049154 | 1253432188 | 1474247942 | 2629254217 | 141E-06 | 222E-05 |

**TABLE S5**

| Gene | ENSMBL | logFC_KO_F | logCPM_KO_F | F_KO_F | PValue_KO_F | FDR_KO_F |
|---|---|---|---|---|---|---|
| Cd209a | ENSMUSG00000031494 | -1186295271 | 1808285601 | 9691514627 | 875E-05 | 0000601903 |
| Ighg2c | ENSMUSG00000076612 | -8555348452 | 5389418847 | 50579294 | 669E-05 | 0000503946 |
| Treml4 | ENSMUSG00000051682 | -849001336 | 0828531959 | 7415263298 | 153E-05 | 000020144 |
| Pklr | ENSMUSG00000041237 | -833712057 | 2036168879 | 5740680952 | 414E-05 | 0000369206 |
| Hbb-bs | ENSMUSG00000052305 | -7441408512 | 3617436048 | 5438211739 | 508E-05 | 0000420995 |
| Hpgd | ENSMUSG00000031613 | -7086131075 | 1207256177 | 4803580742 | 812E-05 | 000057248 |
| Tspo2 | ENSMUSG00000023995 | -6613541598 | 1265206508 | 4733499283 | 858E-05 | 0000593064 |
| Adgre4 | ENSMUSG00000032915 | -6587086422 | 2148954275 | 7307241452 | 162E-05 | 0000209486 |
| Slc40a1 | ENSMUSG00000025993 | -6577423469 | 2591415042 | 5971747755 | 356E-05 | 0000334863 |
| Siglech | ENSMUSG00000051504 | -6443494894 | 0698677244 | 4643843161 | 921E-05 | 000062651 |
| Vcam1 | ENSMUSG00000027962 | -6024850825 | 253113907 | 4793354384 | 818E-05 | 0000574986 |
| Csf3r | ENSMUSG00000028859 | -5769646482 | 183629828 | 8382793914 | 944E-06 | 0000152852 |
| Shank3 | ENSMUSG00000022623 | -541906441 | 1649558916 | 1054716982 | 376E-06 | 865E-05 |
| Dach1 | ENSMUSG00000055639 | -5381704803 | 1093674478 | 1150669806 | 264E-06 | 708E-05 |
| Klrb1a | ENSMUSG00000030361 | -525750246 | 0590547957 | 1572615604 | 733E-07 | 369E-05 |
| Cyp27a1 | ENSMUSG00000026170 | -4971663715 | 2018977669 | 7083577849 | 183E-05 | 0000224901 |
| Ifi207 | ENSMUSG00000073490 | -4762041863 | 1674089838 | 6510424801 | 255E-05 | 0000272339 |
| Itgad | ENSMUSG00000070369 | -462497497 | 1360722017 | 4163999262 | 0000137766 | 000081497 |
| Csf1r | ENSMUSG00000024621 | -4501685309 | 427861798 | 4346322425 | 0000117696 | 0000736984 |
| Dnase1l3 | ENSMUSG00000025279 | -4409029895 | 1496342282 | 5259792111 | 577E-05 | 0000455684 |
| Clec9a | ENSMUSG00000046080 | -4401382736 | 079874601 | 4947089295 | 727E-05 | 0000530473 |
| Ptpro | ENSMUSG00000030223 | -437786495 | 1705463088 | 6075298529 | 333E-05 | 0000322192 |
| Mpeg1 | ENSMUSG00000046805 | -4177645418 | 5228526679 | 4408255726 | 0000111709 | 0000710499 |
| Plbd1 | ENSMUSG00000030214 | -4007997874 | 3794653507 | 471556593 | 870E-05 | 0000598791 |
| Adgrl2 | ENSMUSG00000028184 | -389731406 | 1400700163 | 8548326454 | 873E-06 | 0000145297 |
| Elane | ENSMUSG00000020125 | -3869208397 | 3246467672 | 3965313692 | 0000164666 | 0000917423 |
| Tmem121b | ENSMUSG00000094626 | -3862632974 | 0501385636 | 6928376822 | 200E-05 | 0000238912 |
| Klf1 | ENSMUSG00000054191 | -38104529 | 2628238152 | 4406590287 | 0000111865 | 0000710746 |
| Klri2 | ENSMUSG00000043932 | -3764042896 | 2931656111 | 1112365484 | 303E-06 | 767E-05 |
| Ly6d | ENSMUSG00000034634 | -3736462532 | 2503364407 | 4874036695 | 769E-05 | 0000551412 |
| Fibcd1 | ENSMUSG00000026841 | -3682668519 | 0590335325 | 1033448438 | 408E-06 | 915E-05 |
| Gfi1b | ENSMUSG00000026815 | -3602080735 | 1904264289 | 5002960902 | 697E-05 | 0000516752 |
| Gm22748 | ENSMUSG00000064427 | -3597931242 | 089227378 | 5425249186 | 513E-05 | 0000423385 |
| Gm6377 | ENSMUSG00000048621 | -3589660732 | 0712701744 | 7172422593 | 175E-05 | 000021861 |
| Trim30b | ENSMUSG00000052749 | -3507976919 | 1621923427 | 4541064413 | 0000100085 | 0000659747 |
| Cr2 | ENSMUSG00000026616 | -3498199512 | 2969260621 | 7398717075 | 155E-05 | 0000202732 |
| Dmbt1 | ENSMUSG00000047517 | -348501558 | 0464029793 | 4074617571 | 000014914 | 0000858872 |
| Mpo | ENSMUSG00000009350 | -3471691894 | 4576343456 | 417664536 | 0000136244 | 0000808468 |
| Il13ra1 | ENSMUSG00000017057 | -3466795769 | 115727405 | 8370712342 | 949E-06 | 0000153132 |
| Cd300a | ENSMUSG00000034652 | -3377505401 | 2620512251 | 4791214782 | 820E-05 | 0000575015 |
| Rnd3 | ENSMUSG00000017144 | -3375190177 | 126680446 | 4908753734 | 749E-05 | 0000542285 |
| Mdga1 | ENSMUSG00000043557 | -3314414979 | 0568283273 | 4941780668 | 730E-05 | 0000531772 |
| Cpne7 | ENSMUSG00000034796 | -3254629517 | 1007026868 | 959188521 | 551E-06 | 0000106079 |
| Irgc1 | ENSMUSG00000062028 | -3223292413 | 0325826881 | 658443028 | 244E-05 | 000026667 |
| Gm43063 | ENSMUSG00000105731 | -3187813565 | 0506341079 | 4931656068 | 736E-05 | 0000534917 |
| Fcer2a | ENSMUSG00000005540 | -3153136009 | 2750055164 | 5458912532 | 501E-05 | 0000416951 |
| H2bc8 | ENSMUSG00000058385 | -3084491749 | 0710126088 | 3934359268 | 000016942 | 0000937163 |
| Spns2 | ENSMUSG00000040447 | -3058518247 | 2123532505 | 7715904284 | 131E-05 | 0000182925 |
| Adgrg1 | ENSMUSG00000031785 | -2960499923 | 4010664605 | 1111447371 | 304E-06 | 768E-05 |
| Gm24119 | ENSMUSG00000077167 | -2920516633 | 0521473106 | 5581815362 | 460E-05 | 0000395825 |
| Gm3739 | ENSMUSG00000091472 | -2808201441 | 0825439518 | 474378534 | 851E-05 | 0000589287 |
| Slc45a3 | ENSMUSG00000026435 | -2799976196 | 1901331916 | 4431199696 | 0000109587 | 0000700674 |
| Klhl4 | ENSMUSG00000025597 | -2772506804 | 0633505631 | 6501333262 | 256E-05 | 0000272526 |
| Cyp4f18 | ENSMUSG00000003484 | -2771254074 | 1665837897 | 5054919556 | 670E-05 | 0000504485 |

| | | | | | | |
|---|---|---|---|---|---|---|
| Cmklr1 | ENSMUSG00000042190 | -2725514112 | 0984350732 | 5053971917 | 671E-05 | 0000504485 |
| Mfsd2b | ENSMUSG00000037336 | -2710955955 | 3991819087 | 5239620065 | 586E-05 | 0000459366 |
| Nfam1 | ENSMUSG00000058099 | -2553833863 | 4029483347 | 4560790448 | 985E-05 | 0000652768 |
| Pyroxd2 | ENSMUSG00000060224 | -2534746533 | 2479671247 | 529135807 | 564E-05 | 0000448985 |
| Apoe | ENSMUSG00000002985 | -2523518197 | 5280445166 | 5341751579 | 544E-05 | 000043774 |
| Maged1 | ENSMUSG00000025151 | -251497244 | 1404753294 | 1262095881 | 181E-06 | 603E-05 |
| Cebpa | ENSMUSG00000034957 | -2420838521 | 2346455177 | 5256874825 | 578E-05 | 0000456072 |
| Cxxc5 | ENSMUSG00000046668 | -2398721932 | 2945395763 | 6073698033 | 333E-05 | 0000322261 |
| Srsf12 | ENSMUSG00000054679 | -2373136903 | 1706942352 | 6111894551 | 325E-05 | 0000317358 |
| Ptgs2 | ENSMUSG00000032487 | -2351193935 | 1181990697 | 3979300507 | 0000162572 | 000091037 |
| Snord49a | ENSMUSG00000065734 | -2342198577 | 0761788859 | 4307754036 | 0000121625 | 0000752144 |
| Tyrobp | ENSMUSG00000030579 | -2341001678 | 3769667369 | 4828189833 | 796E-05 | 000056436 |
| Klrb1b | ENSMUSG00000079298 | -2332382234 | 0701292891 | 4630229294 | 931E-05 | 0000631267 |
| Tppp3 | ENSMUSG00000014846 | -230376666 | 1218207053 | 5957818412 | 359E-05 | 0000336055 |
| Ncmap | ENSMUSG00000043924 | -2282498232 | 2044732502 | 3942794897 | 0000168107 | 0000932316 |
| Snord55 | ENSMUSG00000092680 | -2265393722 | 0473733207 | 4874397621 | 769E-05 | 0000551412 |
| Gm12207 | ENSMUSG00000086920 | -2199385863 | 1122848954 | 4298473174 | 0000122594 | 0000754152 |
| Ovgp1 | ENSMUSG00000074340 | -2175825224 | 2889642352 | 1155345077 | 260E-06 | 705E-05 |
| Boll | ENSMUSG00000025977 | -2171892655 | 1314163242 | 534656467 | 542E-05 | 0000437002 |
| Gm17096 | ENSMUSG00000090397 | -2162954873 | 0789999452 | 4626546414 | 934E-05 | 0000632079 |
| Nckap5los | ENSMUSG00000089968 | -2130284754 | 0639978595 | 3987616642 | 0000161342 | 000090598 |
| Trav8-2 | ENSMUSG00000085302 | -2086676073 | 0672396586 | 5390316016 | 526E-05 | 0000430096 |
| Tsc22d3 | ENSMUSG00000031431 | -2061721351 | 7648313658 | 1452062363 | 102E-06 | 440E-05 |
| Afap1 | ENSMUSG00000029094 | -2053586144 | 4300524647 | 1659141983 | 587E-07 | 338E-05 |
| Selenow | ENSMUSG00000041571 | -2052649897 | 645322409 | 5047233965 | 534E-09 | 381E-06 |
| Auts2 | ENSMUSG00000029673 | -2027801209 | 0837642044 | 9933233644 | 479E-06 | 0000100281 |
| Amigo2 | ENSMUSG00000048218 | -200812627 | 5095514695 | 9194731837 | 653E-06 | 0000118888 |
| Il6ra | ENSMUSG00000027947 | -2006238035 | 7978193169 | 1325658889 | 148E-06 | 540E-05 |
| Gfi1 | ENSMUSG00000029275 | 2002681902 | 5774102475 | 3345659847 | 308E-08 | 772E-06 |
| Gm28187 | ENSMUSG00000099375 | 2020620887 | 2437445288 | 6990824987 | 193E-05 | 0000232738 |
| Il6 | ENSMUSG00000025746 | 202876128 | 0708483128 | 7124750651 | 179E-05 | 0000221876 |
| Fzd10 | ENSMUSG00000081683 | 2033412919 | 0451909733 | 3917029945 | 0000172155 | 0000946968 |
| Hk1os | ENSMUSG00000085347 | 2035799523 | 0563997718 | 8846815139 | 762E-06 | 0000132466 |
| Bspry | ENSMUSG00000028392 | 2036321121 | 1915077881 | 4501474986 | 0000103388 | 0000674774 |
| Gm33682 | ENSMUSG00000112971 | 2037862361 | 1906071952 | 9800590113 | 505E-06 | 0000102153 |
| Gm45250 | ENSMUSG00000110317 | 2038109091 | 2162553618 | 5844378506 | 386E-05 | 00003552 |
| AI480526 | ENSMUSG00000090086 | 205858027 | 4268159904 | 6202894224 | 307E-05 | 0000306434 |
| Ankrd37 | ENSMUSG00000050914 | 2070307724 | 2971472287 | 4209560172 | 000013238 | 0000792513 |
| NA | ENSMUSG00000097732 | 2075331797 | 0658017612 | 5844509354 | 386E-05 | 00003552 |
| Mapk12 | ENSMUSG00000022610 | 2081874415 | 2341797658 | 5855631476 | 383E-05 | 0000353663 |
| Gm28053 | ENSMUSG00000098715 | 2118550539 | 2608684769 | 102765324 | 417E-06 | 925E-05 |
| Phf19 | ENSMUSG00000026873 | 2128568706 | 4197417824 | 400907579 | 0000158222 | 0000892794 |
| Atp6v1g2 | ENSMUSG00000024403 | 21429448 | 2900778785 | 1055109594 | 375E-06 | 865E-05 |
| Gas2l1 | ENSMUSG00000034201 | 2167006613 | 1734934347 | 43876438 | 0000113659 | 0000719131 |
| Gm37169 | ENSMUSG00000103175 | 2204437952 | 4583774591 | 2538296996 | 991E-08 | 145E-05 |
| Gm38102 | ENSMUSG00000104156 | 2220064889 | 2254241771 | 4337512546 | 000011858 | 0000739889 |
| Npas2 | ENSMUSG00000026077 | 2226716619 | 0993248477 | 1011257562 | 445E-06 | 958E-05 |
| Mir142b | ENSMUSG00000065420 | 2231642202 | 1733034788 | 4123967816 | 0000142723 | 0000835469 |
| Selp | ENSMUSG00000026580 | 2275913708 | 1329162649 | 847801414 | 902E-06 | 0000148883 |
| Nrp2 | ENSMUSG00000025969 | 2279800402 | 3257558041 | 3940907819 | 00001684 | 0000933511 |
| A930004D18Rik | ENSMUSG00000054057 | 2290951368 | 110299209 | 1096542341 | 321E-06 | 784E-05 |
| Gm47345 | ENSMUSG00000110815 | 2301863136 | 2431077045 | 4165432475 | 0000137593 | 000081497 |
| Gm10688 | ENSMUSG00000111637 | 2375947714 | 0526025867 | 6789887112 | 216E-05 | 0000248502 |
| Cdkl2 | ENSMUSG00000029403 | 2405157269 | 1832310812 | 779413267 | 126E-05 | 0000178664 |
| 2500002B13Rik | ENSMUSG00000096917 | 2418302097 | 0989405067 | 5166955725 | 617E-05 | 0000475948 |
| 5033430I15Rik | ENSMUSG00000038152 | 2464802725 | 0455708971 | 580605189 | 396E-05 | 0000360094 |
| Gm26532 | ENSMUSG00000097296 | 2467055245 | 0802503694 | 5487726192 | 491E-05 | 0000411891 |

| | | | | | | |
|---|---|---|---|---|---|---|
| 4932422M17Rik | ENSMUSG00000105881 | 2474510139 | 1479199866 | 7322007539 | 161E-05 | 0000208431 |
| Lrrn4 | ENSMUSG00000043110 | 2476869398 | 0322076721 | 5584289106 | 460E-05 | 0000395437 |
| Prss16 | ENSMUSG00000006179 | 2543599657 | 2198216074 | 1105873408 | 310E-06 | 776E-05 |
| Ankrd29 | ENSMUSG00000057766 | 2583922169 | 0570988504 | 3879032459 | 0000178345 | 0000973075 |
| Fam83d | ENSMUSG00000027654 | 2666183959 | 2852850687 | 3883688881 | 0000177572 | 0000971041 |
| Neat1 | ENSMUSG00000092274 | 2717484193 | 6884294637 | 1196438952 | 225E-06 | 652E-05 |
| Efr3b | ENSMUSG00000020658 | 2825122467 | 0785111853 | 4466626213 | 0000106406 | 0000687954 |
| Echdc2 | ENSMUSG00000028601 | 2909826808 | 063095186 | 1085557429 | 334E-06 | 805E-05 |
| Slc29a4 | ENSMUSG00000050822 | 2927885223 | 1000401567 | 5824409065 | 391E-05 | 000035799 |
| Pkd1l2 | ENSMUSG00000034416 | 2972607011 | 0538813066 | 6619897461 | 239E-05 | 0000263882 |
| Espnl | ENSMUSG00000049515 | 3179702044 | 0373396954 | 4895249805 | 756E-05 | 0000545738 |
| Fam241b | ENSMUSG00000020083 | 3469489291 | 02312176 | 8423180213 | 926E-06 | 0000150564 |
| Slc7a11 | ENSMUSG00000027737 | 3546179024 | 1238448564 | 4509085601 | 0000102743 | 0000671637 |
| Ccl17 | ENSMUSG00000031780 | 3569405325 | 0156583902 | 8647060056 | 834E-06 | 000014101 |
| Dll4 | ENSMUSG00000027314 | 3788947994 | 1080278727 | 1244421376 | 192E-06 | 622E-05 |
| Slc6a4 | ENSMUSG00000020838 | 3872473318 | 1026548944 | 641565388 | 270E-05 | 000028196 |
| Insrr | ENSMUSG00000005640 | 390883012 | 0087702466 | 7514211556 | 145E-05 | 0000195077 |
| Cacna1b | ENSMUSG00000004113 | 3984667803 | 1203972077 | 4988923978 | 704E-05 | 0000518436 |
| Timp3 | ENSMUSG00000020044 | 4018660157 | 0990012626 | 6725579579 | 225E-05 | 0000254976 |
| Gm10800 | ENSMUSG00000075014 | 4202469589 | 100308536 | 8458439646 | 911E-06 | 0000149486 |
| Peg10 | ENSMUSG00000092035 | 422322139 | 1308356638 | 7823741865 | 124E-05 | 0000176836 |
| Gm10602 | ENSMUSG00000073985 | 4245824886 | 0091305308 | 4375845418 | 0000114794 | 0000725555 |
| Syt13 | ENSMUSG00000027220 | 437764237 | 0608811084 | 171698712 | 509E-07 | 330E-05 |
| Gm28874 | ENSMUSG00000100813 | 4415604775 | 1439842718 | 7230351331 | 169E-05 | 0000214813 |
| Igsf5 | ENSMUSG00000000159 | 4557723432 | 0508607596 | 1178185614 | 240E-06 | 683E-05 |
| Zfp941 | ENSMUSG00000060314 | 4964491439 | 0701711995 | 5147237532 | 626E-05 | 0000481642 |
| Gm16045 | ENSMUSG00000086050 | 785164856 | 0126927575 | 1541076168 | 797E-07 | 390E-05 |
| Gzme | ENSMUSG00000022156 | 1138123162 | 1318663419 | 5572300431 | 680E-07 | 353E-05 |

**TABLE S6**

| gene_id | gene_name | logFC | logCPM | F | PValue | FDR |
|---|---|---|---|---|---|---|
| ENSMUSG00000092341.6 | Malat1 | -10032268 | 829051423 | 294471465 | 113E-13 | 284E-09 |
| ENSMUSG00000006058.11 | Snf8 | -17590961 | 608181762 | 874015424 | 742E-09 | 933E-05 |
| ENSMUSG00000090877.4 | Hspa1b | 322754044 | 477808595 | 781034037 | 189E-08 | 000015885 |
| ENSMUSG00000096768.9 | Gm47283 | 159401137 | 529507053 | 696578685 | 481E-08 | 000030269 |
| ENSMUSG00000091971.4 | Hspa1a | 314048541 | 472686925 | 665156601 | 696E-08 | 000035047 |
| ENSMUSG00000114203.2 | Gm35279 | 476290568 | 043620143 | 48283631 | 812E-07 | 000340621 |
| ENSMUSG00000105263.2 | Gm42427 | 16221625 | 511037729 | 390631297 | 368E-06 | 001321738 |
| ENSMUSG00000005483.11 | Dnajb1 | 121454344 | 641497923 | 38308279 | 420E-06 | 001322504 |
| ENSMUSG00000097141.3 | Gm10524 | 139790094 | 345424646 | 373471362 | 500E-06 | 001398078 |
| ENSMUSG00000121042.1 | Gm33887 | 146744881 | 531086338 | 355084752 | 702E-06 | 001767938 |
| ENSMUSG00000024644.18 | Cndp2 | -11846287 | 335801579 | 325912352 | 123E-05 | 002825061 |
| ENSMUSG00000119904.1 | n-R5s125 | -2805354 | 0948882 | 31037164 | 169E-05 | 003544061 |
| ENSMUSG00000057897.15 | Camk2b | 156841857 | 201886496 | 306021741 | 185E-05 | 003578047 |
| ENSMUSG00000032766.10 | Gng11 | -23471341 | 112130661 | 300491013 | 207E-05 | 003727618 |
| ENSMUSG00000092609.2 | Gm20481 | 335471598 | 053201616 | 28975465 | 260E-05 | 004366374 |
| ENSMUSG00000110537.2 | Gm4316 | 12984562 | 586401355 | 279125662 | 327E-05 | 005152143 |
| ENSMUSG00000104876.2 | Trdc | -09081753 | 420431245 | 271078914 | 391E-05 | 005791964 |

**TABLE S7**

| gene_id | gene_name | logFC | logCPM | F | PValue | FDR |
|---|---|---|---|---|---|---|
| ENSMUSG00000092341.6 | Malat1 | -1289106146 | 5674821317 | 5857651359 | 199E-20 | 486E-16 |
| ENSMUSG00000054072.13 | Iigp1 | 3767629758 | 3786289724 | 1422355372 | 364E-13 | 445E-09 |
| ENSMUSG00000079339.8 | Ifit1bl1 | 6328851815 | 2488418422 | 1263949385 | 164E-12 | 134E-08 |
| ENSMUSG00000074896.4 | Ifit3 | 5371600466 | 4032397223 | 1230451984 | 230E-12 | 141E-08 |
| ENSMUSG00000033355.7 | Rtp4 | 3864489448 | 4966775479 | 11867457 | 362E-12 | 177E-08 |
| ENSMUSG00000025498.16 | Irf7 | 3131399139 | 4326155065 | 1148590877 | 543E-12 | 221E-08 |
| ENSMUSG00000056071.13 | S100a9 | 3482650173 | 4922650349 | 855747814 | 185E-10 | 646E-07 |
| ENSMUSG00000027514.15 | Zbp1 | 2215053664 | 5399032568 | 8078975478 | 358E-10 | 109E-06 |
| ENSMUSG00000056054.10 | S100a8 | 3760887254 | 5240087683 | 7645228956 | 668E-10 | 181E-06 |
| ENSMUSG00000066677.13 | Ifi208 | 5715341324 | 1184393399 | 7438370586 | 908E-10 | 222E-06 |
| ENSMUSG00000121042.1 | Gm33887 | 2810203647 | 4597596639 | 7141926601 | 142E-09 | 316E-06 |
| ENSMUSG00000062488.10 | Ifit3b | 4813740431 | 2189299234 | 6653894388 | 308E-09 | 627E-06 |
| ENSMUSG00000104713.5 | Gbp6 | 2536515931 | 3073555636 | 6164532152 | 694E-09 | 131E-05 |
| ENSMUSG00000078921.4 | Tgtp2 | 2097281225 | 575959896 | 6085491869 | 795E-09 | 139E-05 |
| ENSMUSG00000097141.3 | Gm10524 | 3025980648 | 2906415969 | 5876082025 | 115E-08 | 187E-05 |
| ENSMUSG00000029561.18 | Oasl2 | 3747577558 | 1373065819 | 5751207057 | 143E-08 | 218E-05 |
| ENSMUSG00000105263.2 | Gm42427 | 2616072628 | 4498204073 | 5610749559 | 184E-08 | 254E-05 |
| ENSMUSG00000073491.12 | Ifi213 | 3800646202 | 1552514275 | 5603079142 | 187E-08 | 254E-05 |
| ENSMUSG00000037849.9 | Ifi206 | 4529394484 | 0910800397 | 5407257579 | 268E-08 | 344E-05 |
| ENSMUSG00000046031.8 | Calhm6 | 3217092653 | 1997699356 | 5289274891 | 334E-08 | 408E-05 |
| ENSMUSG00000034459.9 | Ifit1 | 3147102258 | 1806127404 | 5253417465 | 357E-08 | 416E-05 |
| ENSMUSG00000032690.17 | Oas2 | 5759140949 | 0567690894 | 5129200571 | 453E-08 | 483E-05 |
| ENSMUSG00000078922.10 | Tgtp1 | 1928418755 | 5898104481 | 5128077255 | 454E-08 | 483E-05 |
| ENSMUSG00000040809.11 | Chil3 | 3633011395 | 3644257685 | 5063586006 | 515E-08 | 524E-05 |
| ENSMUSG00000049723.15 | Mmp12 | 292350576 | 3475065596 | 4927274422 | 672E-08 | 654E-05 |
| ENSMUSG00000006281.9 | Tep1 | 2713111387 | 2203372195 | 4909953964 | 695E-08 | 654E-05 |
| ENSMUSG00000019467.16 | Arhgef25 | -6419910504 | 0757659969 | 4575588943 | 137E-07 | 0000123899 |
| ENSMUSG00000110537.2 | Gm4316 | 2001313696 | 5553962229 | 4524137988 | 152E-07 | 000012924 |
| ENSMUSG00000039236.19 | Isg20 | 1882175619 | 3699612045 | 4520854634 | 153E-07 | 000012924 |
| ENSMUSG00000101939.2 | Gm28438 | 1569762798 | 6770168481 | 4445648275 | 179E-07 | 0000146138 |
| ENSMUSG00000064360.1 | mt-Nd3 | 1567168046 | 6770726004 | 4430457881 | 185E-07 | 0000146138 |
| ENSMUSG00000078606.9 | Gvin2 | 2071883631 | 3703061331 | 4128223555 | 355E-07 | 0000271535 |
| ENSMUSG00000079363.8 | Gbp4 | 16650887 | 4215095916 | 4103074441 | 376E-07 | 0000278321 |
| ENSMUSG00000056290.17 | Ms4a4b | 1933106168 | 5228251265 | 4074418964 | 400E-07 | 0000287825 |
| ENSMUSG00000045868.13 | Gvin1 | 2114354408 | 3334606091 | 3968223159 | 508E-07 | 0000354513 |
| ENSMUSG00000054942.14 | Miga1 | 2110743503 | 4156735888 | 3735792814 | 865E-07 | 0000587144 |
| ENSMUSG00000079362.6 | Gm43302 | 2406350712 | 1713278276 | 3659509968 | 103E-06 | 000068321 |
| ENSMUSG00000070427.5 | Il18bp | 2588402968 | 1512755588 | 3605085263 | 118E-06 | 0000746963 |
| ENSMUSG00000037921.16 | Ddx60 | 410864186 | 0575440904 | 3599661858 | 119E-06 | 0000746963 |
| ENSMUSG00000001128.8 | Cfp | 2265179748 | 217603307 | 3584233007 | 124E-06 | 0000755578 |
| ENSMUSG00000032484.9 | Ngp | 4416128291 | 0838175665 | 3567163313 | 129E-06 | 0000757114 |
| ENSMUSG00000091144.3 | Phf11c | 3337010001 | 1471520905 | 3562977654 | 130E-06 | 0000757114 |
| ENSMUSG00000039997.17 | Ifi203 | 2475996049 | 2542666135 | 351782565 | 145E-06 | 0000824272 |
| ENSMUSG00000028270.13 | Gbp2 | 1554737469 | 5749385236 | 3486561813 | 156E-06 | 0000868792 |
| ENSMUSG00000078763.3 | Slfn1 | 1741141891 | 393735808 | 3462428981 | 166E-06 | 0000900759 |
| ENSMUSG00000094345.3 | Igkv14-126 | 2162923517 | 6128026135 | 3346643146 | 220E-06 | 000117099 |
| ENSMUSG00000032596.15 | Uba7 | 1949151914 | 2655209503 | 3304455101 | 245E-06 | 0001272873 |
| ENSMUSG00000038357.11 | Camp | 4375873708 | 0496686786 | 3263637492 | 271E-06 | 0001380487 |
| ENSMUSG00000092021.10 | Gbp11 | 3467639366 | 0774242395 | 3230648379 | 295E-06 | 0001469511 |
| ENSMUSG00000105373.2 | Gm42429 | 3937602508 | 0660771696 | 3217144977 | 305E-06 | 0001490152 |
| ENSMUSG00000106734.5 | Gm20559 | 1950422017 | 2364912057 | 318744452 | 329E-06 | 0001575323 |
| ENSMUSG00000078490.11 | Cfap74 | 1540083142 | 3473249139 | 3130082044 | 381E-06 | 0001780948 |
| ENSMUSG00000037902.19 | Sirpa | 5082879504 | 0044885618 | 3124442467 | 386E-06 | 0001780948 |
| ENSMUSG00000020009.14 | Ifngr1 | 1384841609 | 4829910068 | 3062964653 | 453E-06 | 0002049026 |
| ENSMUSG00000097113.9 | Gm19705 | 1764609789 | 3667179834 | 3028418564 | 495E-06 | 0002201292 |
| ENSMUSG00000020388.13 | Pdlim4 | 2041501216 | 2706415565 | 2991181382 | 546E-06 | 0002383732 |
| ENSMUSG00000078920.4 | Ifi47 | 1265592968 | 7556694053 | 2974258274 | 571E-06 | 0002448683 |
| ENSMUSG00000024675.20 | Ms4a4c | 2058953392 | 2573406454 | 2953800249 | 603E-06 | 0002497832 |
| ENSMUSG00000076934.3 | Iglv1 | 2394745908 | 4572666889 | 295369114 | 603E-06 | 0002497832 |

| | | | | | | |
|---|---|---|---|---|---|---|
| ENSMUSG00000067149.7 | Jchain | 313619349 | 1633495552 | 2915831852 | 667E-06 | 0002716077 |
| ENSMUSG00000120570.1 | Gm57321 | 179616725 | 3170465785 | 269311256 | 122E-05 | 0004894829 |
| ENSMUSG00000069792.6 | Wfdc17 | 3612773702 | 1097451474 | 2677899586 | 127E-05 | 0005023741 |
| ENSMUSG00000096393.2 | Fam240a | 1805770068 | 2585897459 | 2671203653 | 130E-05 | 0005037 |
| ENSMUSG00000043263.14 | Ifi209 | 1998852182 | 2033526184 | 2645854532 | 139E-05 | 0005196476 |
| ENSMUSG00000078122.6 | F630028O10Rik | 4233996877 | 0862768203 | 2643781801 | 140E-05 | 0005196476 |
| ENSMUSG00000076940.4 | Iglv2 | 1876773149 | 4522554523 | 2643322439 | 140E-05 | 0005196476 |
| ENSMUSG00000033460.15 | Armcx1 | -4236215013 | -0052233103 | 263800915 | 142E-05 | 0005196476 |
| ENSMUSG00000029298.16 | Gbp9 | 1834585926 | 2267045129 | 2611747889 | 153E-05 | 0005501108 |
| ENSMUSG00000006058.11 | Snf8 | -1129759181 | 6886207998 | 260726859 | 155E-05 | 0005501108 |
| ENSMUSG00000101483.3 | 1700016L21Rik | 3127813829 | 1421971885 | 2564231905 | 175E-05 | 0006124711 |
| ENSMUSG00000105906.2 | Iglc1 | 2746174123 | 4848294368 | 2552117587 | 182E-05 | 0006250127 |
| ENSMUSG00000108414.3 | Snhg1 | 1149682521 | 105661604 | 2538144408 | 189E-05 | 0006413821 |
| ENSMUSG00000060675.14 | Plaat3 | 108707064 | 5716010301 | 2518840605 | 200E-05 | 000664619 |
| ENSMUSG00000018927.4 | Ccl6 | 2245893549 | 353444685 | 2513459329 | 203E-05 | 000664619 |
| ENSMUSG00000051748.3 | Wfdc21 | 3583108845 | 0755359064 | 2511447431 | 204E-05 | 000664619 |
| ENSMUSG00000023349.15 | Clec4n | 2955170811 | 3125796753 | 2487287142 | 219E-05 | 0007030685 |
| ENSMUSG00000024677.14 | Ms4a6b | 1143881011 | 6831976323 | 2479124664 | 224E-05 | 0007104725 |
| ENSMUSG00000117465.2 | Gm49980 | 1737172933 | 3326236833 | 2444623454 | 247E-05 | 0007750816 |
| ENSMUSG00000076937.4 | Iglc2 | 2149891116 | 5354347979 | 240968662 | 274E-05 | 0008473496 |
| ENSMUSG00000001270.10 | Ckb | -1275798918 | 5344717519 | 2380406415 | 298E-05 | 0009118333 |
| ENSMUSG00000088252.3 | Snord13 | 1132640631 | 6350725984 | 2366684919 | 311E-05 | 0009309255 |
| ENSMUSG00000018906.15 | P4ha2 | -2370956007 | 1486720811 | 2364997721 | 312E-05 | 0009309255 |
| ENSMUSG00000095079.7 | Igha | 3328511103 | 0929478445 | 2331525171 | 345E-05 | 0010155533 |
| ENSMUSG00000091649.4 | Phf11b | 1628569489 | 2965599316 | 2320003196 | 357E-05 | 0010384533 |
| ENSMUSG00000014195.17 | Dnajc7 | 1056252305 | 7675370929 | 2297765534 | 381E-05 | 0010966627 |
| ENSMUSG00000079017.4 | Ifi27l2a | 1052311243 | 8873580934 | 2284938748 | 396E-05 | 0011263636 |
| ENSMUSG00000096108.3 | Ighv11-2 | 2252088912 | 3404425424 | 2271499442 | 413E-05 | 0011592728 |
| ENSMUSG00000121353.1 | ENSMUSG00000121353 | 1206540702 | 4343540194 | 226347776 | 423E-05 | 0011741046 |
| ENSMUSG00000058470.11 | Gm8369 | 1058081668 | 5971612034 | 2245699242 | 446E-05 | 0012248897 |
| ENSMUSG00000026104.15 | Stat1 | 106959741 | 5435303272 | 2224596971 | 475E-05 | 0012912382 |
| ENSMUSG00000105096.2 | Gbp10 | 2030598145 | 1313951783 | 2218686008 | 484E-05 | 0012916292 |
| ENSMUSG00000102037.2 | Bcl2a1a | 1336110515 | 4294193688 | 2217262652 | 486E-05 | 0012916292 |
| ENSMUSG00000070501.15 | Ifi214 | 2657766088 | 096218522 | 2207950898 | 500E-05 | 0013144659 |
| ENSMUSG00000044703.6 | Phf11a | 3031397256 | 0678021522 | 2202736426 | 508E-05 | 0013213145 |
| ENSMUSG00000018899.18 | Irf1 | 1108192238 | 4365144981 | 2195134476 | 520E-05 | 0013380901 |
| ENSMUSG00000003484.5 | Cyp4f18 | 4029324576 | 0773168554 | 2191048817 | 526E-05 | 0013407845 |
| ENSMUSG00000110386.2 | Gm42031 | 1329202423 | 6573547624 | 2172376282 | 557E-05 | 0014050532 |
| ENSMUSG00000034634.8 | Ly6d | 2492898654 | 3189149116 | 2159846284 | 579E-05 | 0014452957 |
| ENSMUSG00000069516.9 | Lyz2 | 2742346321 | 3044208514 | 2145777946 | 605E-05 | 0014869014 |
| ENSMUSG00000037683.15 | Armc3 | 2742589348 | 0695691286 | 2144078816 | 608E-05 | 0014869014 |
| ENSMUSG00000044827.11 | Tlr1 | 2742965562 | 0720561646 | 2119650046 | 656E-05 | 001587767 |
| ENSMUSG00000030584.15 | Dpf1 | -5236048449 | -0239460278 | 2111649706 | 672E-05 | 0016117389 |
| ENSMUSG00000087627.3 | A230059L01Rik | 113296583 | 573496273 | 2106343098 | 684E-05 | 0016226407 |
| ENSMUSG00000045932.13 | Ifit2 | 233269187 | 0634661558 | 2099828724 | 698E-05 | 0016322267 |
| ENSMUSG00000070327.15 | Rnf213 | 1389711342 | 2763261745 | 2098273672 | 701E-05 | 0016322267 |
| ENSMUSG00000017734.16 | Dbndd2 | 347448606 | 0078491089 | 20718091 | 761E-05 | 0017562037 |
| ENSMUSG00000056763.17 | Cspp1 | 138610197 | 3366136219 | 2064059654 | 780E-05 | 0017825859 |
| ENSMUSG00000103349.2 | Gm36888 | 2131983143 | 107147874 | 2056999353 | 798E-05 | 0018056788 |
| ENSMUSG00000094796.3 | BC147527 | 2161236774 | 118158437 | 2049238403 | 817E-05 | 0018333275 |
| ENSMUSG00000024644.18 | Cndp2 | -1223208792 | 4279889819 | 2045342947 | 827E-05 | 0018390861 |
| ENSMUSG00000069307.8 | H2bc23 | -1005340354 | 5643768058 | 2040255425 | 841E-05 | 0018519834 |
| ENSMUSG00000032870.9 | Smap2 | 133215998 | 3272381665 | 2021938753 | 891E-05 | 0019395619 |
| ENSMUSG00000024659.16 | Anxa1 | 2192638751 | 2684968719 | 2019988774 | 896E-05 | 0019395619 |
| ENSMUSG00000037649.11 | H2-DMa | 1211277846 | 6537081458 | 2011970908 | 920E-05 | 0019652761 |
| ENSMUSG00000002227.16 | Mov10 | 1388446975 | 3878359968 | 2008919822 | 929E-05 | 0019652761 |
| ENSMUSG00000069303.8 | H2bc24 | -0993190162 | 5652827567 | 2007576192 | 932E-05 | 0019652761 |
| ENSMUSG00000118012.2 | Gm46620 | 2400869213 | 1062978714 | 2003733153 | 944E-05 | 0019724184 |
| ENSMUSG00000031125.3 | 3830403N18Rik | 240869758 | 1502457785 | 1992738625 | 978E-05 | 002025347 |
| ENSMUSG00000118026.3 | Gm50335 | 2243416491 | 0736326613 | 1976456243 | 000010297 | 0021154656 |
| ENSMUSG00000120167.1 | Gm56655 | 1122699325 | 5220125347 | 196751468 | 0000105959 | 002158742 |
| ENSMUSG00000067297.6 | Ifit1bl2 | 5619098784 | -0826282727 | 2012492699 | 0000110994 | 002227599 |

| | | | | | | |
|---|---|---|---|---|---|---|
| ENSMUSG00000021538.4 | Il9 | 2094767243 | 5349614272 | 1952578978 | 0000111161 | 002227599 |
| ENSMUSG00000026822.15 | Lcn2 | 4492672397 | 0146783888 | 1939083557 | 0000116097 | 0023062763 |
| ENSMUSG00000090272.10 | Mndal | 1757091706 | 2344662317 | 1936748184 | 0000116974 | 0023062763 |
| ENSMUSG00000031104.3 | Rab33a | -2507343051 | 1965317427 | 193032662 | 0000119424 | 0023357378 |
| ENSMUSG00000038642.11 | Ctss | 1376548065 | 655703106 | 1896603047 | 000013322 | 0025848914 |
| ENSMUSG00000097893.10 | 1700034P13Rik | 1570990693 | 2369806819 | 189333484 | 0000134645 | 0025919649 |
| ENSMUSG00000059498.14 | Fcgr3 | 2292584218 | 1675312215 | 188871612 | 0000136686 | 0026107042 |
| ENSMUSG00000117485.3 | Gm19696 | -3886620066 | 0459856893 | 1879923611 | 0000140664 | 0026658548 |
| ENSMUSG00000105703.3 | Gm43305 | 125809689 | 554791243 | 1858782677 | 0000150743 | 0028272314 |
| ENSMUSG00000050921.13 | P2ry10 | 1040437172 | 4428320038 | 1857143937 | 0000151556 | 0028272314 |
| ENSMUSG00000029610.14 | Aimp2 | -0956731125 | 5880587019 | 1854957188 | 0000152648 | 0028272314 |
| ENSMUSG00000054892.15 | Txk | 1515243342 | 3284917635 | 1851307537 | 000015449 | 0028398213 |
| ENSMUSG00000116504.3 | I730030J21Rik | -1264523359 | 4031626692 | 1844510786 | 0000157982 | 0028734249 |
| ENSMUSG00000024673.10 | Ms4a1 | 3158046437 | 3358964458 | 1840788354 | 0000159931 | 0028734249 |
| ENSMUSG00000017652.17 | Cd40 | 1501952999 | 3901327594 | 1840750665 | 0000159951 | 0028734249 |
| ENSMUSG00000060441.16 | Trim5 | 4224971568 | -0617697291 | 1838730445 | 0000161019 | 0028734249 |
| ENSMUSG00000028268.15 | Gbp3 | 1428086357 | 2466102986 | 182505646 | 0000168454 | 0029843146 |
| ENSMUSG00000097705.3 | Gm26740 | 2294068424 | 1739197914 | 1821841069 | 0000170255 | 0029945206 |
| ENSMUSG00000021880.8 | Rnase6 | 2314545528 | 1518916135 | 1784494346 | 0000192747 | 0033659176 |
| ENSMUSG00000055865.9 | Tafa3 | 2984276376 | -0068066958 | 1761340378 | 0000208271 | 0036112121 |
| ENSMUSG00000120084.1 | Gm57127 | 1764368151 | 1300375947 | 1755447869 | 0000212431 | 0036574091 |
| ENSMUSG00000071552.6 | Tigit | -124457105 | 4046542252 | 1751545165 | 0000215236 | 0036797762 |
| ENSMUSG00000090024.2 | Gm16350 | -2157081297 | 1605455594 | 1745132269 | 000021993 | 0037339195 |
| ENSMUSG00000040253.16 | Gbp7 | 1539826282 | 1802219673 | 174098076 | 0000223027 | 0037395419 |
| ENSMUSG00000034422.15 | Parp14 | 1776239816 | 1373598952 | 1739677726 | 0000224009 | 0037395419 |
| ENSMUSG00000045092.9 | S1pr1 | 174793801 | 1863575207 | 1738566113 | 000022485 | 0037395419 |
| ENSMUSG00000005125.14 | Ndrg1 | -1745937863 | 2522122059 | 1733820418 | 0000228479 | 0037742224 |
| ENSMUSG00000026946.10 | Nmi | 1009718556 | 5520400099 | 1730334295 | 0000231184 | 0037932872 |
| ENSMUSG00000076532.4 | Igkv4-91 | 1915607488 | 2974109456 | 1724359781 | 0000235901 | 0038448785 |
| ENSMUSG00000026921.21 | Egfl7 | -1107084586 | 3801872211 | 1721808457 | 0000237947 | 0038525345 |
| ENSMUSG00000018387.13 | Shroom1 | -1683127441 | 2926397022 | 1690793081 | 0000264388 | 0042524646 |
| ENSMUSG00000044583.14 | Tlr7 | 254937799 | 1040357819 | 1676896107 | 0000277238 | 0044190659 |
| ENSMUSG00000046727.14 | Cystm1 | -1557471184 | 342625639 | 167571488 | 0000278361 | 0044190659 |
| ENSMUSG00000106567.3 | 2010309G21Rik | 2773100152 | 4620968727 | 1670133698 | 0000283731 | 0044752646 |
| ENSMUSG00000115681.3 | Gm2682 | 1059299441 | 5140270631 | 1666081152 | 00002877 | 0045087774 |
| ENSMUSG00000105186.2 | Gm43778 | 1009739185 | 5533662378 | 1660587552 | 0000293175 | 0045653199 |
| ENSMUSG00000022475.20 | Hdac7 | -1127917615 | 4940601267 | 1657897767 | 0000295897 | 0045785338 |
| ENSMUSG00000073430.7 | Gm10505 | 1417061305 | 3649815816 | 1652220181 | 000030173 | 0046207972 |
| ENSMUSG00000059142.17 | Zfp945 | 1728233805 | 1856297349 | 1651567671 | 0000302408 | 0046207972 |
| ENSMUSG00000027398.14 | Il1b | 1771733472 | 2048859345 | 1648308187 | 0000305821 | 0046312331 |
| ENSMUSG00000027776.13 | Il12a | 1981763301 | 1675892531 | 1647304405 | 000030688 | 0046312331 |
| ENSMUSG00000020108.5 | Ddit4 | -111103581 | 3433458224 | 1642213843 | 0000312312 | 0046677885 |
| ENSMUSG00000027333.19 | Smox | -146122446 | 3156970642 | 1641464097 | 0000313121 | 0046677885 |
| ENSMUSG00000114608.4 | Gm36161 | 4155523995 | -0624683155 | 1637836407 | 0000317065 | 0046979469 |
| ENSMUSG00000034656.19 | Cacna1a | 175565529 | 1276894395 | 1611065104 | 0000347871 | 0051233446 |
| ENSMUSG00000035692.8 | Isg15 | 1192145637 | 5066392576 | 1606332074 | 0000353643 | 005157649 |
| ENSMUSG00000062991.10 | Nrg1 | 2466128528 | 1139686602 | 1605701283 | 000035442 | 005157649 |
| ENSMUSG00000024679.11 | Ms4a6d | 1900643829 | 097356747 | 1597027233 | 000036529 | 0052724057 |
| ENSMUSG00000032359.15 | Ctsh | 2086530314 | 3453136874 | 1594451641 | 0000368586 | 0052724057 |
| ENSMUSG00000097134.3 | 1110002J07Rik | 4039135667 | -0736694602 | 1594304573 | 0000368775 | 0052724057 |
| ENSMUSG00000120432.1 | Gm56600 | 2147730056 | 1847722775 | 1586318098 | 0000379205 | 0053850439 |
| ENSMUSG00000057143.16 | Trim12c | 1607461379 | 1345871311 | 1584703522 | 0000381351 | 0053850439 |
| ENSMUSG00000029530.17 | Ccr9 | 5985185858 | -0622806999 | 1623208932 | 0000384061 | 0053850439 |
| ENSMUSG00000079293.13 | Clec7a | 2819265158 | 0161010725 | 1581637111 | 0000385464 | 0053850439 |
| ENSMUSG00000109251.2 | E230032D23Rik | 175913182 | 1187537599 | 1579937209 | 0000387764 | 0053864006 |
| ENSMUSG00000024990.14 | Rbp4 | 6416758914 | -0278102092 | 1616603904 | 0000392641 | 0054100652 |
| ENSMUSG00000037548.16 | H2-DMb2 | 1988122759 | 3856671415 | 1574748565 | 0000394877 | 0054100652 |
| ENSMUSG00000074466.14 | Gm15417 | -1034229845 | 9245022608 | 1573278367 | 0000396918 | 0054100652 |
| ENSMUSG00000020610.18 | Amz2 | -1366046535 | 456447099 | 1572272957 | 000039832 | 0054100652 |
| ENSMUSG00000003541.7 | Ier3 | -1555978471 | 4793838406 | 1569383607 | 0000402378 | 0054349953 |
| ENSMUSG00000008193.14 | Spib | 2878833041 | 0842639708 | 1563509626 | 0000410766 | 005517808 |
| ENSMUSG00000119262.1 | Snord3b1 | 1002259256 | 4785085935 | 1557537846 | 0000419486 | 0055987551 |

| | | | | | | |
|---|---|---|---|---|---|---|
| ENSMUSG00000068874.14 | Selenbp1 | -2423133753 | 4518546341 | 1556263298 | 0000421372 | 0055987551 |
| ENSMUSG00000046794.10 | Ppp1r3b | -2066558935 | 1888687322 | 1552992499 | 0000426255 | 00563302 |
| ENSMUSG00000040751.13 | Lat2 | 1508676923 | 3359229572 | 1541040243 | 000044462 | 005844118 |
| ENSMUSG00000030410.17 | Dmwd | -1714454096 | 1399651754 | 1537493949 | 000045023 | 0058561142 |
| ENSMUSG00000052749.11 | Trim30b | 3169611807 | -0159188961 | 1537435489 | 0000450323 | 0058561142 |
| ENSMUSG00000037443.14 | Cep85 | 1137588208 | 4675143909 | 1533395866 | 0000456807 | 0059090093 |
| ENSMUSG00000113866.2 | Gm47450 | 1020404217 | 5281727733 | 1531389177 | 0000460066 | 0059198343 |
| ENSMUSG00000037463.15 | Fbxo27 | 5379809324 | -0967610603 | 1562874405 | 0000470603 | 0059966314 |
| ENSMUSG00000033577.19 | Myo6 | 1027107363 | 5829425639 | 1524005357 | 000047227 | 0059966314 |
| ENSMUSG00000094347.3 | Or6c208 | 1277014571 | 2852170217 | 1523336782 | 0000473392 | 0059966314 |
| ENSMUSG00000029816.11 | Gpnmb | 2802302458 | 0395981599 | 1513970611 | 0000489415 | 0061676382 |
| ENSMUSG00000105033.2 | Gm43423 | -1442113016 | 7313834802 | 1502512327 | 000050981 | 0063917049 |
| ENSMUSG00000087175.9 | Gm15133 | 2075202429 | 1018905833 | 1501067062 | 0000512446 | 006391978 |
| ENSMUSG00000008540.12 | Mgst1 | 3738285842 | -0375792543 | 1488786231 | 0000535445 | 0066449509 |
| ENSMUSG00000028943.19 | Espn | -1816253361 | 2771775719 | 1485569341 | 000054165 | 0066880096 |
| ENSMUSG00000079597.3 | Cstdc4 | 2501404647 | 0814279606 | 1477085673 | 0000558387 | 0068600179 |
| ENSMUSG00000025163.7 | Cd7 | 1732303353 | 1262811396 | 1474058728 | 0000564491 | 00687276 |
| ENSMUSG00002074955.1 | Snord3b4 | 0985908783 | 4822042059 | 1473785413 | 0000565046 | 00687276 |
| ENSMUSG00000111083.2 | Gm48274 | 3508535764 | -0556362968 | 1467628371 | 0000577699 | 0069741411 |
| ENSMUSG00002076650.1 | Snord3b1 | 0988376127 | 4800856195 | 1466028007 | 0000581037 | 0069741411 |
| ENSMUSG00002075931.1 | Snord3b2 | 0970373055 | 48107481 | 1465596992 | 0000581939 | 0069741411 |
| ENSMUSG00000000753.16 | Serpinf1 | -1420998089 | 4537009622 | 1464046437 | 0000585198 | 0069789906 |
| ENSMUSG00000039883.6 | Lrrc17 | 1446449305 | 7384035009 | 1462291954 | 000058891 | 00698916 |
| ENSMUSG00000041481.18 | Serpina3g | 108353793 | 3466749226 | 1459603615 | 0000594646 | 006992719 |
| ENSMUSG00000106874.3 | Gm20186 | 0806904502 | 7204901789 | 145947105 | 000059493 | 006992719 |
| ENSMUSG00000119648.1 | Snord3b3 | 0984091242 | 4794644271 | 1452209339 | 0000610733 | 0071441183 |
| ENSMUSG00000006651.9 | Aplp1 | -2572103082 | -020123225 | 1449897985 | 0000615857 | 0071697491 |
| ENSMUSG00000060586.12 | H2-Eb1 | 171575738 | 6336213082 | 144321717 | 0000630927 | 0072951391 |
| ENSMUSG00000019122.9 | Ccl9 | 1215497997 | 2986392993 | 1442488159 | 0000632596 | 0072951391 |
| ENSMUSG00000030142.11 | Clec4e | 1741832609 | 2638053852 | 1438177822 | 0000642556 | 0073752134 |
| ENSMUSG00000060591.10 | Ifitm2 | -0944323397 | 5528573429 | 1429109116 | 0000664064 | 0075660245 |
| ENSMUSG00000023084.12 | Lrrc71 | -2625931878 | -0165222165 | 142856885 | 0000665369 | 0075660245 |
| ENSMUSG00000041538.17 | H2-Ob | 2321638981 | 2200749547 | 1426142252 | 0000671267 | 0075977455 |
| ENSMUSG00000004814.11 | Ccl24 | 1421844741 | 48661556 | 1424721168 | 0000674746 | 0076019352 |
| ENSMUSG00000038085.14 | Cnbd2 | 10867955 | 4345756082 | 1415513671 | 0000697764 | 0078252038 |
| ENSMUSG00000035299.17 | Mid1 | 1747515907 | 1514001622 | 141234217 | 0000705886 | 0078801415 |
| ENSMUSG00000047735.15 | Samd9l | 2627451735 | 0151837275 | 1407773384 | 0000717765 | 0079763272 |
| ENSMUSG00000021057.16 | Akap5 | 1354189168 | 7385253654 | 1400685093 | 000073662 | 0081488157 |
| ENSMUSG00000027199.15 | Gatm | -1923690268 | 1624288098 | 1397483413 | 000074531 | 0081973724 |
| ENSMUSG00000119851.1 | Snord3b4 | 0985311345 | 4801379878 | 1396604128 | 0000747715 | 0081973724 |
| ENSMUSG00000068245.15 | Phf11d | 2560237728 | 0040723845 | 1391022981 | 000076318 | 0083295616 |
| ENSMUSG00000036594.16 | H2-Aa | 199580005 | 6893880099 | 1385615949 | 0000778489 | 0083798281 |
| ENSMUSG00000030148.16 | Clec4a2 | 5341364942 | -0979595768 | 1415831534 | 0000782809 | 0083798281 |
| ENSMUSG00000037161.15 | Mgarp | -1758567259 | 3909098589 | 1384000655 | 0000783126 | 0083798281 |
| ENSMUSG00000036570.15 | Fxyd1 | -5187817011 | -1221861734 | 1414251189 | 0000787188 | 0083798281 |
| ENSMUSG00000121137.1 | Gm56555 | 082988645 | 5459884939 | 1382509419 | 0000787433 | 0083798281 |
| ENSMUSG00000036435.14 | Exoc1 | -0976486986 | 7003092181 | 138219267 | 0000788351 | 0083798281 |
| ENSMUSG00000073902.6 | Gvin3 | 2763250445 | -0200298849 | 1379110877 | 0000797344 | 008438728 |
| ENSMUSG00000020646.18 | Mboat2 | -2242132395 | 0936982297 | 1377123889 | 00008032 | 0084640679 |
| ENSMUSG00000099974.2 | Bcl2a1d | 1064089215 | 7204871167 | 1371017627 | 0000821487 | 0086196251 |
| ENSMUSG00000078616.3 | Trim30c | 3088372877 | -0474916483 | 1366331955 | 0000835823 | 00873256 |
| ENSMUSG00000021831.10 | Ero1a | -1262609736 | 4673815414 | 1356002187 | 000086838 | 0090341068 |
| ENSMUSG00000119761.1 | Snord3b2 | 093918269 | 481295691 | 1350148253 | 0000887431 | 0091931834 |
| ENSMUSG00000074141.14 | Il4i1 | 1603848906 | 3312590505 | 1347118138 | 0000897468 | 0092579275 |
| ENSMUSG00000017754.14 | Pltp | 2619427142 | 1798510842 | 1340772102 | 0000918884 | 0094390198 |
| ENSMUSG00000076939.3 | Iglv3 | 2352409626 | 1165003114 | 1335533298 | 0000936976 | 0095845935 |
| ENSMUSG00000105547.2 | Iglc3 | 2088340266 | 5109162986 | 132540427 | 0000973047 | 0099121027 |

**TABLE S8**

| Cluster number | Cluster color | Protein name |
|---|---|---|
| 1 | Red | Ddx60 |
| 1 | Red | Gbp2 |
| 1 | Red | Gbp3 |
| 1 | Red | Gbp6 |
| 1 | Red | Gbp7 |
| 1 | Red | Ifi209 |
| 1 | Red | Ifi27l2a |
| 1 | Red | Ifi47 |
| 1 | Red | Ifit1 |
| 1 | Red | Ifit1bl1 |
| 1 | Red | Ifit1bl2 |
| 1 | Red | Ifit2 |
| 1 | Red | Ifit3 |
| 1 | Red | Ifit3b |
| 1 | Red | Ifngr1 |
| 1 | Red | Iigp1 |
| 1 | Red | Il12a |
| 1 | Red | Il18bp |
| 1 | Red | Il9 |
| 1 | Red | Irf1 |
| 1 | Red | Irf7 |
| 1 | Red | Isg15 |
| 1 | Red | Isg20 |
| 1 | Red | Ms4a4b |
| 1 | Red | Ms4a4c |
| 1 | Red | Ms4a6b |
| 1 | Red | Nmi |
| 1 | Red | Oas2 |
| 1 | Red | Oasl2 |
| 1 | Red | Parp14 |
| 1 | Red | Phf11d |
| 1 | Red | Rnf213 |
| 1 | Red | Rtp4 |
| 1 | Red | Samd9l |
| 1 | Red | Stat1 |
| 1 | Red | Uba7 |
| 1 | Red | Zbp1 |
| 2 | Brown | Anxa1 |
| 2 | Brown | Camp |
| 2 | Brown | Ccl24 |
| 2 | Brown | Ccl6 |
| 2 | Brown | Ccl9 |
| 2 | Brown | Ccr9 |

| | | |
|---|---|---|
| 2 | Brown | Cfp |
| 2 | Brown | Chil3 |
| 2 | Brown | Clec4e |
| 2 | Brown | Clec4n |
| 2 | Brown | Ctsh |
| 2 | Brown | Ctss |
| 2 | Brown | Fcgr3 |
| 2 | Brown | H2-Aa |
| 2 | Brown | H2-DMa |
| 2 | Brown | H2-DMb2 |
| 2 | Brown | H2-Eb1 |
| 2 | Brown | H2-Ob |
| 2 | Brown | Iglc2 |
| 2 | Brown | Il1b |
| 2 | Brown | Jchain |
| 2 | Brown | Lat2 |
| 2 | Brown | Lcn2 |
| 2 | Brown | Lyz2 |
| 2 | Brown | Ms4a1 |
| 2 | Brown | Ms4a6d |
| 2 | Brown | Ngp |
| 2 | Brown | S100a8 |
| 2 | Brown | S100a9 |
| 2 | Brown | Sirpa |
| 2 | Brown | Spib |
| 2 | Brown | Tlr7 |
| 2 | Brown | Txk |
| 2 | Brown | Wfdc17 |
| 3 | Dark Golden | Akap5 |
| 3 | Dark Golden | Myo6 |
| 3 | Dark Golden | Pdlim4 |
| 4 | Green | Cyp4f18 |
| 4 | Green | Plaat3 |
| 5 | Green 2 | Trim12c |
| 5 | Green 2 | Trim5 |
| 6 | Cyan | Tgtp1 |
| 6 | Cyan | Tgtp2 |
| 7 | Dark Cyan | Bcl2a1a |
| 7 | Dark Cyan | Bcl2a1d |
| 8 | Blue | Phf11a |
| 8 | Blue | Phf11b |
| 9 | Medium Blue | Ifi203 |
| 9 | Medium Blue | Mndal |

**TABLE S9**

| Gene_ID | FDR_DTU | Gene name | Number of isoforms | Number of differentially expressed isoforms (KO vs WT) |
|---|---|---|---|---|
| ENSMUSG00000097354.9 | 0 00076427 | 2310001H17Rik | 38 | 3 |
| ENSMUSG00000053063.12 | 0 03906294 | Clec12a | 2 | 2 |
| ENSMUSG00000014195.17 | 1 43E-11 | Dnajc7 | 19 | 7 |
| ENSMUSG00000033373.17 | 0 01076268 | Fntb | 5 | 1 |
| ENSMUSG00000053332.15 | 0 00168573 | Gas5 | 149 | 5 |
| ENSMUSG00000054435.17 | 0 01403486 | Gimap4 | 6 | 1 |
| ENSMUSG00000043505.9 | 0 01762127 | Gimap5 | 4 | 2 |
| ENSMUSG00000001663.11 | 0 03129099 | Gstt1 | 2 | 2 |
| ENSMUSG00000027718.9 | 0 00026601 | Il21 | 2 | 2 |
| ENSMUSG00000026420.17 | 0 00927377 | Il24 | 4 | 2 |
| ENSMUSG00000000869.13 | 0 03329888 | Il4 | 4 | 1 |
| ENSMUSG00000054611.18 | 0 02890363 | Kdm2a | 5 | 1 |
| ENSMUSG00000031683.17 | 0 09584018 | Lsm6 | 4 | 3 |
| ENSMUSG00000021606.9 | 0 | Ndufs6 | 5 | 3 |
| ENSMUSG00000030815.12 | 0 08244566 | Phkg2 | 9 | 1 |
| ENSMUSG00000060675.14 | 0 08639127 | Plaat3 | 3 | 2 |
| ENSMUSG00000023048.14 | 0 04679064 | Prr13 | 5 | 1 |
| ENSMUSG00000006058.11 | 0 01584275 | Snf8 | 5 | 3 |
| ENSMUSG00000108414.3 | 0 | Snhg1 | 26 | 13 |
| ENSMUSG00000085385.9 | 0 09778325 | Snhg17 | 74 | 1 |
| ENSMUSG00000005656.10 | 0 00047815 | Snx6 | 5 | 2 |
| ENSMUSG00000066258.14 | 0 00077081 | Trim12a | 6 | 2 |
| ENSMUSG00000039701.12 | 0 0116619 | Usp53 | 3 | 2 |
| ENSMUSG00000040483.16 | 0 0018117 | Xaf1 | 5 | 1 |
| ENSMUSG00000022100.15 | 0 0398128 | Xpo7 | 4 | 0 |
